# Modelling Subject Variability in the Spatial and Temporal Characteristics of Functional Modes

**DOI:** 10.1101/544817

**Authors:** Samuel J. Harrison, Janine D. Bijsterboch, Andrew R. Segerdahl, Sean P. Fitzgibbon, Seyedeh-Rezvan Farahibozorg, Eugene P. Duff, Stephen M. Smith, Mark W. Woolrich

**Affiliations:** FMRIB, Wellcome Centre for Integrative Neuroimaging, University of Oxford, Oxford, UK; OHBA, Wellcome Centre for Integrative Neuroimaging, University of Oxford, Oxford, UK; Translational Neuromodeling Unit, University of Zurich & ETH Zurich, Zurich, Switzerland; Department of Radiology, Washington University Medical School, Saint Louis, USA; Department of Paediatrics, University of Oxford, Oxford, UK

## Abstract

Recent work has highlighted the scale and ubiquity of subject variability in observations from functional MRI data (fMRI). Furthermore, it is highly likely that errors in the estimation of either the spatial presentation of, or the coupling between, functional regions can confound cross-subject analyses, making accurate and unbiased representations of functional data essential for interpreting any downstream analyses.

Here, we extend the framework of probabilistic functional modes (PFMs) [Harrison et al. 2015] to capture cross-subject variability not only in the mode spatial maps, but also in the functional coupling between modes and in mode amplitudes. A new implementation of the inference now also allows for the analysis of modern, large-scale data sets, and the combined inference and analysis package, PROFUMO, is available from git.fmrib.ox.ac.uk/samh/profumo. Using simulated data, resting-state data from 1,000 subjects collected as part of the Human Connectome Project [Van Essen et al. 2013], and an analysis of 14 subjects in a variety of continuous task-states [Kieliba et al. 2019], we demonstrate how PFMs are able to capture, within a single model, a rich description of how the spatio-temporal structure of resting-state fMRI activity varies across subjects.

We also compare the new PFM model to the well established independent component analysis with dual regression (ICA-DR) pipeline. This reveals that, under PFM assumptions, much more of the (behaviorally relevant) cross-subject variability in fMRI activity should be attributed to the variability in spatial maps, and that, after accounting for this, functional coupling between modes primarily reflects current cognitive state. This has fundamental implications for the interpretation of cross-sectional studies of functional connectivity that do not capture cross-subject variability to the same extent as PFMs.

## 1 Introduction

One of the key changes to the landscape of the analysis of functional connectivity via rfMRI in recent years has been the proliferation of large population-level studies [Van Essen et al. 2012b; Breteler et al. 2014; Bamberg et al. 2015; Miller et al. 2016] and multi-site data-sharing initiatives [Biswal et al. 2010; Scott et al. 2011; Mennes et al. 2013; Poldrack et al. 2013; Thompson et al. 2014; Kennedy et al. 2016; Gorgolewski et al. 2017]^1^. This has allowed investigations into the population-level correlates of fine-grained changes in functional connectivity [E. A. Allen et al. 2011; Dubois and Adolphs 2016], with several studies already finding strong links with a variety of behavioural, genetic and lifestyle factors [Finn et al. 2015; Smith et al. 2015; Colclough et al. 2017; Elliott et al. 2018]; together, these findings augur well for the search for clinically relevant, personalised predictions from functional neuroimaging data [Insel and Cuthbert 2015; Dubois and Adolphs 2016; Abraham et al. 2017; Stephan et al. 2017]. In sum, there has been a shift in what is required of analysis techniques, namely that they must be both interpretable and sensitive to subject-level variability, and at the same time they need to scale to meet the computational demands posed by large data sets.

### 1.1 Implications of variability over subjects

In this paper, we are primarily interested in the interpretation of—and characterisation of the subject variability in—static functional connectivity^2^. Ultimately, static functional connectivity is encapsulated by the dense connectome—by which we mean the time-averaged voxels-by-voxels connectivity matrix, as defined by the statistical relationships between time courses as extracted from functional data [Friston et al. 1993; Friston 2011]. However, dense connectomes are cumbersome computationally, and the natural spatial scale of the functional data is likely to be much lower than the several hundred thousand voxels present in a typical fMRI acquisition [Van Essen et al. 2012a]. In practice, what we are seeking is a parsimonious summary of the static functional connectivity that is both readily interpretable and captures key forms of variability.

The canonical approach for analyses of static functional connectivity is to summarise the high-dimensional data in terms of a comparatively small number of either parcels or functional systems^3^. These are usually defined in terms of their spatial configuration, at which point it is possible to extract representative time courses from functional data and analyse these. There will naturally be variability in functional connectivity in several domains, though based on the above framework we will focus on two key ones here: firstly, we will refer to variability in the size, shape and location of functional regions as *subject variability in spatial organisation*; secondly, we will use *subject variability in temporal features* to denote the changes in summary measures based on said time courses—in particular, the strength of functional connectivity between regions (i.e. functional connectomes). Finally, note that for clarity we will use the term *functional coupling* to specifically refer to the functional connectivity between regions as described by these low-dimensional connectomes^4^.

The assumption that is implicit in either the parcel or system-level analyses is that registration to a common space means that the time courses we extract based on group-level spatial descriptions are an accurate, or at least unbiased, description of each subject’s data. However, given that it is by no means uncommon to observe three-fold variation in the areal extent of regions of primary visual cortex across subjects [Andrews et al. 1997; Dougherty et al. 2003]; or that non-homeomorphic morphological changes, such as subjects exhibiting different number of gyri and sulci, are prevalent [Shackman et al. 2011; Amiez and Petrides 2014] even in identical twins [Bartley et al. 1997; Hasan et al. 2011]; or that macroscale anatomical features are poor predictors of cytoarchitectonic borders [Amunts et al. 2007]; then we should expect there to be substantial disparities in the presentation of functionally homologous regions across subjects, even after nonlinear registration [Brett et al. 2002; Devlin and Poldrack 2007; Van Essen and Dierker 2007; Mueller et al. 2013]. Recent observations have confirmed this for functional data, where it has been shown that this subject variability in spatial organisation ‘can give rise to divergent connectivity estimates from the same seed region in different subjects’ [Gordon et al. 2017a]—with the results from several studies also suggesting that reorganisations of functionally homologous regions that cannot be represented by diffeomorphic warps seem to be commonplace [Hacker et al. 2013; Harrison et al. 2015; Laumann et al. 2015; Glasser et al. 2016a; Gordon et al. 2016; Braga and Buckner 2017; Gordon et al. 2017b; Kong et al. 2018]. Furthermore, these differences have a substantial impact on the data: cross-subject differences in static functional connectivity have been shown to be much larger than either cross-site effects [S. Noble et al. 2017] or cross-condition, within-subject changes [Gratton et al. 2018].

Loosely speaking, these spatial differences in functional connectivity after registration can arise for four reasons: there will naturally be some errors in the registration process, resulting in structural features that are not brought into correspondence; there will be locations where anatomical landmarks bear little relation to functional subdivisions, meaning structural similarity is not a sufficient condition for accurate registration; there will be genuine non-homotopic reorganisations, whereby the standard registration approaches based on diffeomorphic warps could never succeed^5^; and there will be dynamic—either moment-to-moment or state-dependent—changes in the functional connectivity structure [Buckner et al. 2013; Krienen et al. 2014; Salehi et al. 2020]. If these different sources of variability in spatial organisation are not accounted for, then one expects the inferred mode time courses to be a farrago of contributions from the underlying ‘true’ set of modes [Smith et al. 2011; E. A. Allen et al. 2012]. Worse still, if the structural differences capture meaningful cross-subject differences—which they almost certainly will do [Llera et al. 2019]—then the amount of misalignment, and hence the quality of the extracted time courses, will reflect information that is anatomical rather than functional in origin [Bijsterbosch et al. 2018]. This breaks the central tenet of investigations into subject variability in temporal features, as we can no longer assume that a group-level description of the functional architecture is a reliable description of individual subjects, or even that we can use these to extract unbiased estimates of functional coupling. How then, do we proceed from here?

The first approach we could take is to improve the registrations, and hope that better algorithms and utilising a richer feature set to drive the alignment will push individual subjects ever closer towards the group description [Robinson et al. 2014; Tong et al. 2017; Robinson et al. 2018]. Notably however, the multiple recent observations that single functional regions can be manifested as multiple disjoint regions in some subjects, is something that not even advanced functional registration algorithms reliant on diffeomorphic warps can correct for. The minimum requirement for this approach is therefore the use of advanced registration techniques that can non-homotopically reorganise the spatial layout of functional regions, as, for example, introduced by Conroy et al. [2013] and Guntupalli et al. [2016, 2018], or Langs et al. [2010].

The alternative approach, and the one that we take in this paper, is to build algorithms that can extract estimates of subject variability in temporal features while simultaneously accounting for the variable presentation of functional regions at the subject level. Several methods have been proposed to do exactly this, using both hierarchical models of functional systems [Varoquaux et al. 2011; Abraham et al. 2013; Harrison et al. 2015; Li et al. 2017] and parcels [Liu et al. 2012; Langs et al. 2016; Kong et al. 2018]. We provide a more fulsome description of these, and their counterparts that extract subject-specific information given a fixed group template, in Appendix A. However, the majority of these methods have what is potentially a major limitation: the flow of information is almost exclusively from group to subject. In other words, there are only relatively rudimentary efforts to tap into what we might hope is a virtuous cycle: we should be able to use our group-level estimates to infer accurate subject-level information, but, crucially, we should also be able to utilise the observed variability at the subject level to refine our group-level parameterisations. Furthermore, the same process should hold within subjects, such that accurate estimation of the individual spatial presentations should improve evaluation of the temporal information, and vice versa.

Finally, while we have tended to focus on connectomes as the principal temporal feature of interest in the above discussion, there are other types of variability we are interested in. Recent work has shown that, for example, amplitudes—by which we mean any metric which represents the amount of fluctuation in activity of a functional region over time—carry a substantial amount of information about subjects [Zang et al. 2007; Duff et al. 2008; Zou et al. 2008; Miller et al. 2016; Bijsterbosch et al. 2017], provided we are sufficiently careful in how we distinguish changes from those in functional coupling [Duff et al. 2018], and then how we interpret said changes [Qing and Gong 2016]. Amplitudes are therefore another type of subject-specific information that we would hope analysis methods could identify, and more importantly disambiguate from, the types of subject variability we have already discussed. This is an illustrative example of the complexity of the task of characterising functional connectivity: at every level of any perceptual hierarchy of features we impose (i.e. separation into spatial and temporal features, or subdivision of temporal features into amplitudes and coupling), we expect there to be multiple ways to identify the different features, and substantial cross-subject variability that is correlated across the different categories.

### 1.2 Outline

For the rest of this paper, we will outline our approach for simultaneously inferring group- and subject-level descriptions of functional systems. We use the term *mode* to describe our mathematical description of a given system.

To begin with, we present our probabilistic model for these modes, including the way we parameterise subject variability in both spatial and temporal features, and our approach for inference. This is a significant extension of the proof-of-concept method [Harrison et al. 2015] in several key ways: we introduce a new hierarchical model to better capture the functional coupling between modes, incorporate a model for mode amplitudes to engender a cleaner separation between different types of functional variability, and we overhaul the entire implementation to help the inference scale to large data sets.

We then compare the performance of our method with existing approaches. We do this using both simulated and empirical data, namely the complete set of rfMRI data as released by the Human Connectome Project and ‘active-state’ fMRI data from a more conventionally sized study. Finally, we offer some brief discussions as to the significance of our results.

## 2 Model

Our approach infers subject-level probabilistic functional modes (PFMs)—each of which can be thought of as being described by a subject-specific spatial map and a set of time courses—across the whole cohort simultaneously. Ensuring that there is correspondence between the inferred modes across the cohort is a challenge [Esposito et al. 2005], especially on resting-state data where we cannot assume any common temporal structure.

However, we can use the information at the group-level to inform the subject-specific decompositions: both the subject-specific spatial maps and the low-dimensional, between-mode functional connectomes are constrained to vary around their group-level descriptions, and we can also leverage the expected properties of the hæmodynamic response to further constrain the time courses. Moreover, we can use the subject-specific modes to learn about the variability of all these properties, thereby allowing us to not only describe typical patterns of activity, but to also quantify the extent to which observed patterns are atypical. We do this by building, and then inferring upon, a hierarchical probabilistic model for rfMRI data as described by a set of modes, and it is this that we outline in the following section.

### 2.1 Matrix factorisation models

Defining a mode in terms of a spatial map and time course means that it is fundamentally a matrix factorisation approach, a mathematical formulation which underpins principal component analysis, independent component analysis, non-negative matrix factorisation, dictionary learning and several other of the well established methods for extracting modes from rfMRI data. For completeness, we briefly introduce our notation for this class of models before introducing our extensions.

Firstly, each subject, *s*, from a cohort of *S* subjects, is scanned *R*_*s*_ times. Note that we do not assume that each of the runs for a given subject (i.e. *r* ∈ {1, …, *R*_*s*_}) are identical from a modelling standpoint: they could, for example, represent different time points in a longitudinal study, or different conditions^6^, and we may therefore want to treat them differently. The fMRI data are acquired in *V* voxels and at *T* time points, which we reshape into a data matrix ***D***^(*sr*)^ ∈ ℝ^*V* ×*T*^. We do all our analyses after the data has been registered into a common space, so the number of voxels is constant across subjects. We do however allow the number of time points per run to vary (i.e. 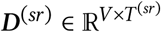), but for notational simplicity we drop any superscripts on *T*.

The problem we are faced with is defining an extension to the standard matrix factorisation approach to account for these multiple data. In the spatial domain, as discussed in the Introduction, we expect between-subject variability in the locations of functional regions, even after registration, and we expect these effects to be amongst the dominant sources of functional variability. We make the pragmatic decision to focus on differences in the static configuration of functional systems specifically, and we target our spatial approach towards what are essentially misalignments.

Therefore, as in Harrison et al. [2015], we model subject and run variability within the matrix factorisation framework as follows. We are looking for a set modes, and we assume that the subject variability in spatial organisation we observe across subjects, by virtue of it being driven primarily by cortical reorganisations, is consistent across all runs for a given subject. This gives a set Of subject-specific spatial maps, ***P***^(*s*)^ ∈ ℝ^*V* ×*M*^, that will potentially be observed multiple times. Furthermore, each run will have its own unique set of time courses, ***A***^(*sr*)^ ∈ ℝ^*M*×*T*^, as well as a set of mode amplitudes, ***h***^(*sr*)^ ∈ ℝ^*M*^. For convenience we adopt the following convention: ***H***^(*sr*)^ ∈ ℝ^*M*×*M*^ ≡ diag(***h***^(*sr*)^). Finally, note that in general we infer a small number of PFMs relative to *V* and *T*, which gives a parsimonious description of the data. However, this means that the factorisation will not be exact, so we express the data as the contribution from the PFMs and a noise term, ***ε***^(*sr*)^ ∈ ℝ^*V* ×*T*^. This set of assumptions allows us to describe the complete model for one run as

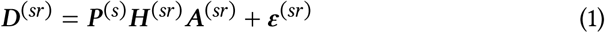

In the following sections, we describe how we model the dependencies between these run-specific decompositions, as well as the key properties of rfMRI data that we are trying to capture. For reference, a full graphical model is provided in the Supplementary Material.

### 2.2 Spatial Model

The spatial model remains conceptually similar to the approach we used in Harrison et al. [ibid.]. For each mode, there is a rich group-level description capturing the mean group maps and typical subject variability around these; as Van Essen and Dierker [2007] discuss, in light of subject variability, it is essential that ‘[regions are] represented probabilistically whenever possible, in a way that reflects variability in cortical convolutions and in [their] size, location, and internal (e.g., topographic) organization’. Similarly, subject maps are parameterised such that they retain the key characteristics of the group maps, but allow for genuine variability while being robust to spurious correlations induced by noise.

A key modification we make to the previous model is to change the way we model the spatial map distribution, by relaxing the delta-Gaussian mixture model to a double-Gaussian mixture model. Previously, the weights in voxels which were inferred to be outside of a given mode were set to exactly zero. In reality however, essentially all voxels will exhibit a *weak* correlation with a given mode time course^7^, and, particularly in studies like the Human Connectome Project with thousands of time points per subject, there is often sufficient evidence *a posteriori* to model this noise as small, but nevertheless non-zero, weights^8^. The new model allows for exactly this type of ‘spurious’ (i.e. statistically but not biologically significant) correlation by including a noise distribution to capture small deviations from zero in the spatial map weights. While we are not interested in these small weights per se, if we do not include a more explicit noise model then the model will erroneously include them as signal thereby hindering our ability to detect genuine ‘neural’ signal.

This contamination by noise happens for three main reasons. Firstly, as Bright and Murphy [2015] recently showed, even well-characterised functional modes can be identified from noise processes like subject motion. Conversely, this implies that even accurately identified modes may well correlate with non-neural processes. Secondly, given the complex, long-range spatial autocorrelations present in fMRI data [Kriegeskorte et al. 2008], fMRI noise processes have a non-trivial structure. This is heightened by spatial smoothing, which is an often used pre-processing step for fMRI data (though less so for modern high spatial and temporal resolution data [Glasser et al. 2016b]). This is advantageous as it ameliorates the problem of residual spatial mis-alignment after registration, but induces heightened spatial correlations in the noise. While it would be possible to model this, estimating—and then correcting for—the true number of spatial degrees of freedom in the data is notoriously difficult [Worsley et al. 1996; Eklund et al. 2016], and would be computationally expensive over a large number of voxels. Finally, in the section on the noise model itself, we demonstrate how unstructured noise can have a stabilising effect on matrix factorisation models. Therefore, we make the pragmatic decision to account for these effects in the spatial model, rather than trying to incorporate a more complex mechanistic model for the noise.

The resulting model takes the following form. For voxel *v* in mode *m*, the subject-specific spatial weights are distributed as follows:

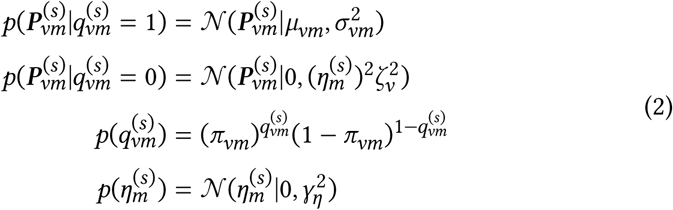

Where 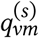 is a binary indicator variable which represents whether a given voxel’s weight is drawn from the signal or the noise component.

This distribution is defined in terms of several group-level hyperparameters: the probability that a given weight is drawn from the signal rather than the noise distribution, *π*_*vm*_; the mean and standard deviation of the signal component, *μ*_*vm*_ and *σ*_*vm*_ respectively; and the new parameters, the standard deviation of the noise component, which we parameterise as 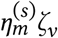 for reasons which we explain in detail later.

Note how much richer this description is than the single set of group-level means that most currently used techniques infer. For example, the *σ*_*vm*_ parameters can capture the types of spatial non-uniformity in subject variability observed by Mueller et al. [2013]. Therefore, when inferring subject maps, the inference will automatically be informed by the data more than the group mean in regions inferred to exhibit high functional heterogeneity over subjects, and vice versa for regions with low subject-to-subject variability.

The model also includes the set of distributions over the group-level hyperpriors (see the Supplementary Material for the way these, and all subsequent, hyper-parameters are specified). Starting with the hyperpriors on the ‘signal’ component, we place a mixture model prior over the group means, which, as in the previous work, is inspired by the spike-slab distribution [Mitchell and Beauchamp 1988; George and McCulloch 1993; Ishwaran and Rao 2005; Titsias and Lázaro-Gredilla 2011]. This encourages sparsity in the group-level spatial maps, thereby encoding ideas about functional segregation, as well as allowing more flexibility when specifying the distribution of the non-zero weights. However, we introduce an extension and model the non-zero weights with a combination of two Gaussians with different variances. This allows the group-level distribution of non-zero spatial weights to have heavier tails than the single Gaussian used in the previous incarnation of the model.

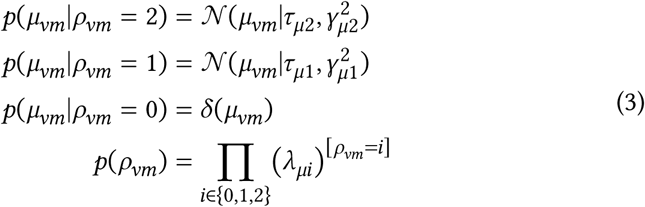

Where *ρ*_*vm*_ is the probability that a voxel in the group map is drawn from each of the three distributions, and [*ρ*_*vm*_ = *i*] is the Iverson bracket.

The group signal standard deviations, *σ*_*vm*_, take an inverse-gamma hyperprior:

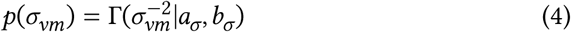

Returning to the hyperpriors on the ‘noise’ component, in Equation 2, the standard deviation of the noise component of the subject-specific spatial map distribution is parameterised as 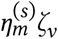. The *ζ* _*v*_ parameter encodes spatial inhomogeneity in the noise variance: for example, we expect more structured noise due to motion around the edges of the brain; similarly, we expect more physiological noise in the brainstem. This group noise standard deviation, *ζ*_*v*_, also takes an inverse-gamma hyperprior:

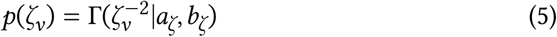

However, we also expect different signal-to-noise ratios, both across subjects and modes. Therefore, we include an extra parameter,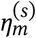, which captures variations in the noise level^9^. We place a weak prior on 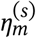, as we want the overall scale of each spatial map to be determined by the signal rather than the noise, as this makes cross-subject analyses more informative:

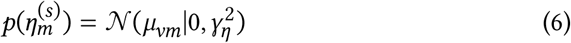

Finally, the last hyperprior to specify is that on the group membership probabilities. This follows a beta distribution:

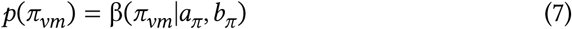

In summary, the model has rich descriptions of the spatial maps, both at the group and subject level, and allows us to encode typical patterns of variability. Furthermore, while we have included a weak sparsity constraint at the group-level, there is no explicit constraint on, for example, orthogonality of the spatial maps. Therefore, the model can capture modes that are highly spatially overlapping in what is arguably a more natural way than independent component analysis—even despite a historic tendency to overstate those criticisms [Beckmann et al. 2005; Smith et al. 2012; Calhoun et al. 2013].

One last point to note is that when we present our results, the group maps we show are the marginal posterior means of the whole spatial distribution, rather than *μ* the parameters themselves. The group-level maps are therefore E [*π*_*vm*_*μ*_*vm*_|𝒟], which has the nice property that it incorporates the uncertainty about whether each voxel is is drawn from the signal or the noise component.

### 2.3 Temporal Model

In the temporal domain, the unconstrained nature of rfMRI data means that we can say relatively little about the time courses from a given run, as there are no external events from which we can search for consistent time-locked patterns of mode activation. However, functional connectomics has shown that, as well as having a consistent group structure, both the interactions between modes and simple amplitude measures encode interesting information about subjects. Similarly, the hæmodynamic processes lend neural processes a distinct temporal signature. That being the case, we wish to formulate a model that primarily captures these two phenomena.

However, we expect the inferred time courses to be corrupted by noise, even if we properly make allowances for the global noise process ***ε***^(*sr*)^. As mentioned in the Spatial Model section, there are likely to be structured noise processes that violate our hæmodynamic assumptions. This needs to be accounted for before we can introduce the targeted models of the BOLD signal.

Analogously to the the spatial model, we extend the model from Harrison et al. [2015] by making the pragmatic decision to allow noisy time courses. Therefore, our time course model contains two terms: the first represents the clean BOLD time courses, ***B***^(*sr*)^, while the second represents the noise that corrupts these, ***ξ***^(*sr*)^. This gives:

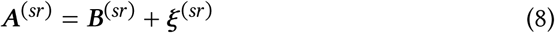

There is an additional benefit of this explicit parameterisation of the BOLD time courses. Recent work has claimed that the [fractional] Amplitude of Low Frequency Fluctuations ([f]ALFF) [Zang et al. 2007; Zou et al. 2008; Zuo et al. 2010a], as derived from rfMRI data, captures aspects of subject variability related to disease. Our parameterisation allows us to derive arelated quantity, which we term the fractional amplitude of BOLD time courses (fABT). This is simply defined as the power in the clean BOLD time courses ***B***^(*sr*)^, relative to the power in the noise time courses ***ξ***^(*sr*)^, calculated for each mode and each run individually. Conceptually, this is very closely related to fALFF, but it has the clear advantage that it does not require defining ‘low’ frequencies in terms of an arbitrary threshold; rather, the signal of interest is based on an explicit model of the HRF. Secondly, the calculated fABT measures specifically relate to the activity in different functional systems which makes the measure more interpretable.

#### 2.3.1 Hæmodynamic model

We use the hæmodynamic response function (HRF) based model that we introduced in Harrison et al. [2015]. This is a relatively simple, computationally efficient, linear model that captures the gross properties of the HRF via the temporal auto-correlations that it induces in the data. We assume a white noise ‘neuronal’ process convolved with a canonical HRF^10^, whose autocorrelation function we can capture using a full covariance matrix, ***K***_***B***_ ∈ ℝ^*T*×*T*^, for all the time points in a given run. As the overall variance of the time courses is arbitrary given the explicit amplitude parameters, we simply ensure that ***K***_***B***_ is scaled such that all entries on the main diagonal are unity.

#### 2.3.2 Subject-level mode interactions

The major extension relative to the previous model is an explicit parameterisation of the functional coupling between modes. As discussed earlier, we expect to observe temporal interactions between modes, and this will lend some structure to the mode time courses. We define these interactions in terms of the precision matrix between the mode time courses. In other words, we combine the HRF-derived autocorrelation structure with a prior on the between-mode precision matrix, ***α***^(*sr*)^ ∈ ℝ^*M* ×*M*^, in a matrix normal distribution.

The combined prior on the hæmodynamic time course for all the PFMs in a given run is then:

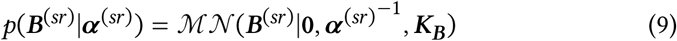

#### 2.3.3 Group-level mode interactions

The temporal interactions between modes have been characterised as having a consistent structure across the group [Shehzad et al. 2009], so we introduce a hierarchical model to capture this. Subject- or run-level variability will manifest itself as deviations from this set of group interactions. This formulation we use is, in essence, the same model as that proposed by Marrelec et al. [2006], but where we have two principal advantages: firstly, inference is informed by the full posteriors on the rest of the model (i.e. rather than point estimates); and, secondly, that the regularisation that arises from these priors will inform the inference of the rest of the model parameters.

Starting at the subject level, we estimate the subject/run-specific temporal precision matrix ***α***^(*sr*)^ to keep track of the functional connectivity between modes. These precision matrices follow a Wishart distribution, and we introduce a hyper-parameter, ***β*** ∈ ℝ^*M*×*M*^, that encourages the interactions to be consistent across subjects and/or runs. This takes the form of a hyperprior on the subject-specific scale matrices, and again this follows a Wishart distribution.

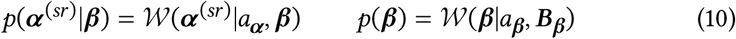

Furthermore, we can also place restrictions on the type of variability we want the model to capture. If, for example, subjects are scanned multiple times but always under the same conditions, then it may well be appropriate to generate a consensus set of interactions for that subject by pooling over runs. We can do this straightforwardly by setting ***α***^(*sr*)^ ≡ ***α***^(*sr*)^. Alternatively, if the runs vary across the group in a consistent way (e.g. ‘before’ and ‘after’ scans) then we may want to explicitly model these conditions as separate entities. We can do this by introducing a family of group-level interactions, 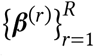, and selectively using these as the hyperpriors on ***α*** as appropriate. This gives us enormous flexibility and allows us to increase our statistical power by making targeted assumptions about the key modes of variation.

#### 2.3.4 Time course specific noise model

The noise time course of mode*m* at time 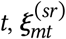, is simply drawn from a Gaussian distribution with precision 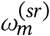. This gives

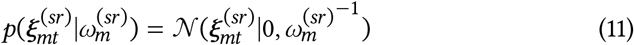

Each 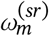 takes a gamma hyperprior:

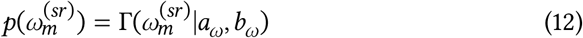

### 2.4 Amplitude Model

Again, the amplitude model is an extension to our previous work. This has a straightforward formulation, with these parameters simply designed to account for the run-to-run variations in the overall activity of each mode. These are parameterised in terms of ***H***^(*sr*)^ ≡ diag(***h***^(*sr*)^), and follow a Gaussian distribution:

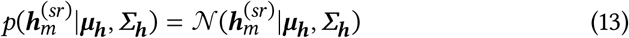

The group-level parameters, ***μ***_***h***_ and ***Σ***_***h***_ capture any consistent cross-subject relationships between the mode amplitudes. For example, Bijsterbosch et al. [2017] recently reported that the amplitudes of sensorimotor modes are correlated with one another, as are the amplitudes of cognitive networks. It is exactly these types of effects that these hyperpriors are able to capture.

The hyperpriors are formulated as follows:

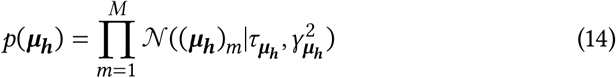

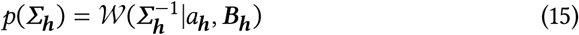

Furthermore, we impose a post-hoc positivity constraint on these variables as part of the inference procedure. As there is a multiplicative ambiguity as to the signs of the components in a matrix factorisation model, we can do this without loss of generality.

### 2.5 Noise Model

The final part of the model left to specify is the noise process,***ε***^(*sr*)^, which we assume is zero-mean, white Gaussian noise, with an overall precision for each run,*ψ*^(*sr*)^. This specifies the likelihood:

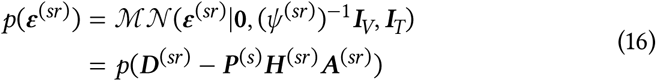

This noise precision then takes a standard gamma hyperprior:

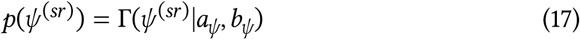

This relatively simple structure assumes that the noise variance is the same in every voxel, which is particularly useful as it allows us to exploit the properties of the matrix normal distribution, leading to very computationally efficient inference [Stegle et al. 2011]. We can preprocess the data in such a way that this is a reasonable assumption to make, and this is discussed in Appendix B.

What is perhaps more problematic is that this model does not acknowledge the spatial smoothness of fMRI data, which means that the noise is not truly independent over voxels. It would be possible to model this, for example by inferring a full spatial covariance matrix for the noise that acknowledged the dependencies between voxels that smoothing introduces. Again, we decide that the benefits of this more complex model are outweighed by the increased computational burden, and again we discuss a way in which we can mitigate the effects of this model misspecification via a relatively straightforward adjustment for the spatial degrees of freedom introduced by Groves et al. [2011], as discussed in Appendix D.

#### 2.5.1 Spatially and temporally specific noise models

One of the key changes to the model as introduced here and its previous incarnation is the way we model noise on the spatial maps and timecourses, as well as the overall noise described above. Interestingly, these different sources of noise can be beneficial for matrix factorisation models even in the absence of the fMRI-specific effects we postulated.

To demonstrate this, we use a simple, single-run version of our generative model, ***D*** = ***PA*** + ***ε***, and we assume the maps and timecourses are full rank to simplify the derivations below. The ordinary-least-squares single-regression estimator for the spatial maps, 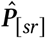, given the ground-truth timecourses is:

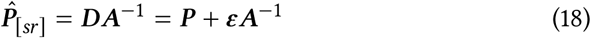

If we instead run dual regression—using the Woodbury matrix identity for the key rearrangements—we find a different estimator for 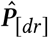:

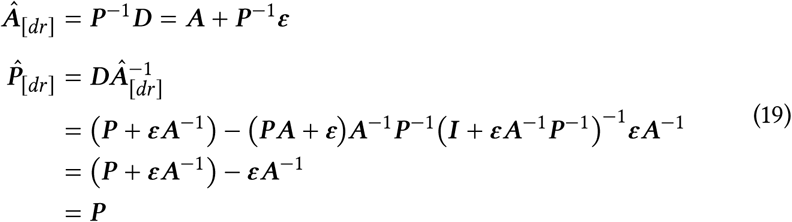

What is surprising is that the dual regression estimator is closer to the ground truth, even though the intermediate timecourses, 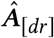, are noisy. This unintuitive behaviour occurs because dual regression involves two regressions on the same noise, and this has concrete implications for the PFM model. When we fit the hæmodynamic model to the timecourses, we exclude the temporally specific noise terms from the estimation of the functional coupling between modes. However, we need to include the temporal noise terms when using the timecourses to estimate the subject-specific spatial maps, as removing it could increase the variance of the inferred maps. The situation is directly analagous with the model for spatial noise: while it is not a quantity of interest for cross-subject modelling, its inclusion can improve the stability of the overall estimation.

In sum, the PROFUMO approach uses the spatially and temporally specific noise where the stabilising effect on matrix factorisation models means that it is expedient to do so, but seeks to avoid letting it confound cross-subject analyses. By way of contrast, dual regression is not naturally able to separate these types of noise.

### 2.6 Inference Approach

We use a computationally efficient variational Bayesian approach to infer upon the probabilistic model outlined above. This technique is well established for graphical models that have a conjugate-exponential structure, as is the mean-field approximation that renders the inference tractable [Attias 2000; MacKay 2003; Winn and Bishop 2005; Blei et al. 2017]; as such, we will not cover the details of that here. In the Appendices, we outline several of the implementation details, including our data preprocessing pipeline, the way we handle large data sets, tweaks to the model and the initialisation procedure.

The combined inference and analysis package, PROFUMO (from PRObabilistic FUnctional MOdes) is available from git.fmrib.ox.ac.uk/samh/profumo and is compatible with FSL [Jenkinson et al. 2012]. All subsequent analyses were performed with version 0.11.1.

The model clearly has a large number of hyperparameters, but as described in the Supplementary Material we can drastically reduce the effective number given that the overall variance of the data is fixed by the internal preprocessing. Furthermore, the vast majority of the parameters that need setting govern the group-level hyperpriors and, as such, are several steps removed from the subject-level decompositions. This means that we can use the same default hyperpriors for all the analyses presented here, and that the inference generalises well across simulated, volumetric, and surface-based data, as well as datasets with very different numbers of subjects.

### 2.7 Model Summary

In summary, we explicitly model many of the properties of rfMRI data within the PROFUMO framework. In the spatial domain, we have a complex group-level model that captures both mean effects and typical patterns of variability, and use these to regularise the subject-specific spatial maps. The temporal model is based around the physiological properties of the BOLD signal, and includes another hierarchical model for the functional coupling between modes. Similarly, we capture differences in the overall activity levels of modes via the amplitude parameters. Finally, we can generate additional summaries by combining parameters as desired, which includes, for example, the measures related to the fractional amplitudes of the BOLD signal.

## 3 Results

Here, we demonstrate the performance of PROFUMO using a set of simulated data and two empirical datasets. All comparisons are with spatial independent component analysis and dual regression (ICA-DR) [Calhoun et al. 2001; Beckmann et al. 2005; Zuo et al. 2010b; Nickerson et al. 2017], as this is what has been used in previous publications on the empirical data.

### 3.1 Simulations

The simulation framework is explicitly designed to be challenging, such that it tests the various ways in which the assumptions the different models make are most likely to be violated. This includes spatial and temporal correlations between components; spatial variability, including a model for misalignments; amplitude variability across subjects and components; a (weakly) nonlinear HRF that varies over both subjects and space; and spatial and temporal smoothness in the residuals. This extends previously published analyses [Harrison et al. 2015; Bijsterbosch et al. 2019], and all simulation code is available from git.fmrib.ox.ac.uk/samh/PFM_ Simulations.

Specifically, we simulate data containing 15 components in a group of 40 subjects, each with two runs containing 10,000 voxels and 500 timepoints at a TR of 2.0 s. A more detailed overview of the data generation procedure is provided in Appendix F. We then test how well PROFUMO and ICA-DR can recover the ground-truth parameters, pooling results across 10 different simulated datasets. Finally, to give more detailed insights into the performance of ICA-DR we include several intermediate steps: firstly, to separate the performance of ICA and dual regression, we include a dual regression analysis starting from the ground-truth spatial maps (GTg-DR); secondly, we include the thresholded variant of dual regression proposed by Bijsterbosch et al. [2019] which is designed to reduce the observed bias in functional coupling (ICA-DRt, GTg-DRt).

Four key performance metrics are shown in Figure 1, and a much more detailed set of comparisons is included in the Supplementary Material. PROFUMO is able to accurately recover spatial maps, amplitudes and functional coupling network matrices (netmats), and much more so than either ICA-DR or the improved thresholded variant (ICA-DRt).

**Figure 1:**
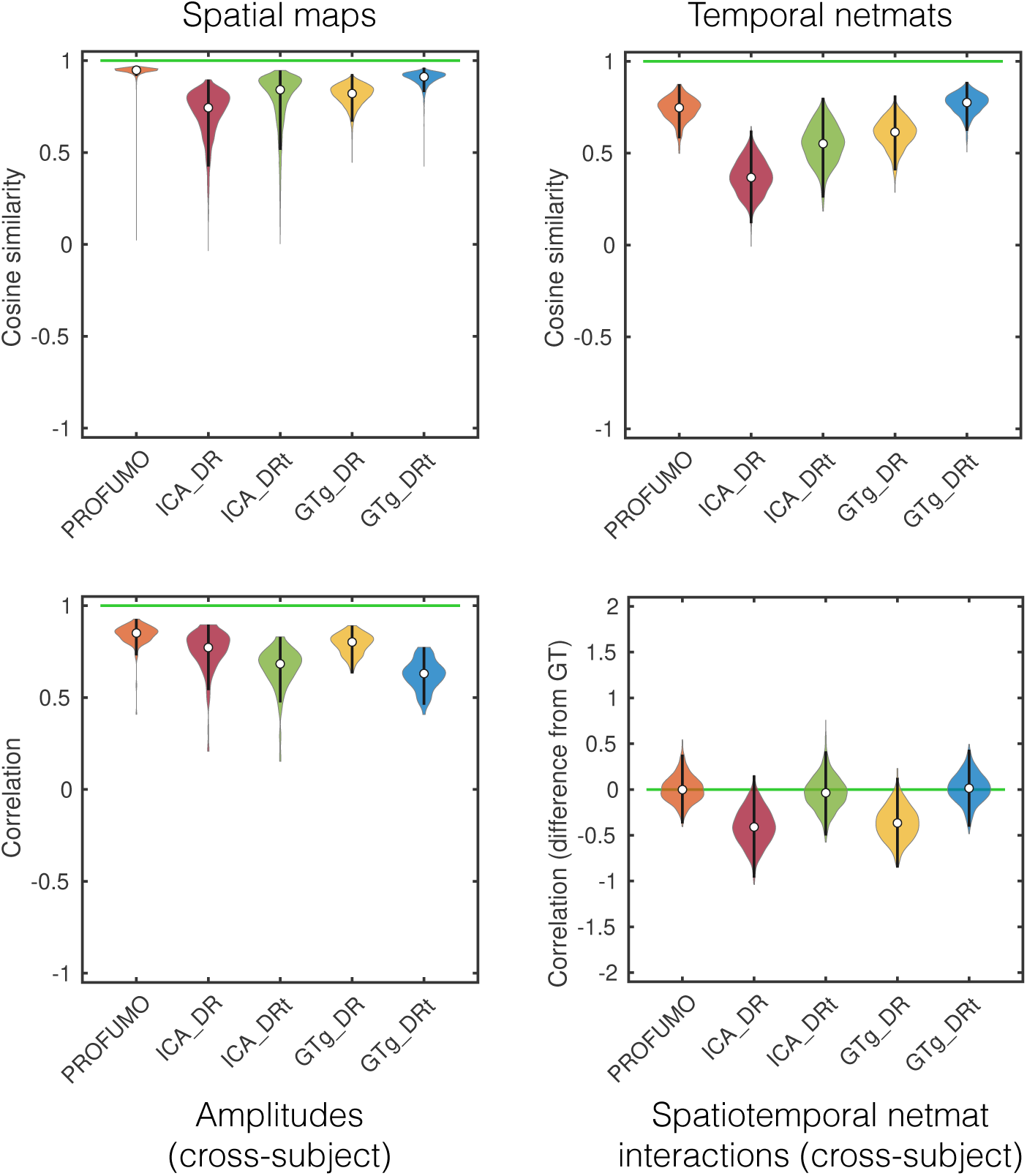
Performance of different algorithms on simulated data. For each metric, optimal performance is shown by the horizontal green line. The metrics are accuracy in recovery of the subject-specific spatial maps, recovery of the run-specific network matrices (netmats), recovery of cross-subject differences in amplitudes (as different approaches normalise the data differently, we look at relative changes in amplitudes across subjects), and any biases in the recovered temporal correlations towards the spatial correlation structure. As well as PROFUMO and ICA-DR, we test dual regression starting with the ground-truth spatial maps (GTg-DR) and thresholded dual regression (ICA-DRt, GTg-DRt).

Crucially, the inferred PFMs are also unbiased in the presence of spatio-temporal correlations between components, unlike ICA-DR. What Bijsterbosch et al. [ibid.] demonstrated was that inaccurate estimation of the group-level spatial correlation structure—an inevitable consequence of the orthogonality constraints of ICA—leads to biased estimates of functional coupling. What we show here is a stronger result: this effect is present even when starting from the correct group-level spatial maps (GTg-DR). In this case, the effect is driven by the mismatch between the true subject-level spatial correlations and those between the group-level maps. In other words, this bias will be present for all dual regression analyses, however the group-level maps are generated.

Furthermore, in the Supplementary Material, we repeat the simulations but with the addition of structured noise, including subject-specific artefacts that can be either spatially specific or global. While the differences between methods are less pronounced, there are still clear benefits to using PROFUMO. However, performance does suffer, and, as such, we strongly recommend ICA-based artefact removal before running PROFUMO, as is the case for the two empirical datasets presented here.

### 3.2 Human Connectome Project data

To evaluate the ability of PROFUMO to detect subtle subject-specific variations in functional connectivity, we use data from the Human Connectome Project (HCP) [Van Essen et al. 2012b; 2013]. This is for two main reasons. Firstly, the most recent data release includes high-quality functional data from over 1,000 subjects and, as such, is an ideal test for methods that purport to be suitable for population-level studies as mentioned in the Introduction. Secondly, the functional pipeline has been published [Smith et al. 2013a] and the results are available to download—thereby offering a comparison that is independently verifiable. The pipeline uses spatial ICA and dual regression to characterise subject variability in both spatial and temporal features. While it would also be possible to examine the equivalent pipeline based on temporal ICA, this has not been used so extensively: for example, the HCP’s MegaTrawl analyses are based on the spatial ICA pipeline^11^. Similarly, this pipeline does not make use of the new thresholded variant of dual regression. Based on the simulated data, this would improve the results slightly, though PROFUMO still outperforms this variant on essentially all of the metrics we tested. Again, the aim is to use the existing, publicly available results as a baseline.

A key aim of modern, large-scale studies of functional connectivity is to relate neurobiological changes to individual differences in genetic, lifestyle and behavioural factors. Using the HCP data also allows us to do this by comparing our results with a wide range of information about subjects. The data involves a battery of cognitive tests, and also records a range of metrics based on health and lifestyle: we will refer to differences in these as *subject variability in behavioural measures*. We can indirectly assess the effects of genetics and environement by calculating the heritability of key imaging metrics; we do this by utilising the fact that many twins and siblings were involved in the study. Finally, we can examine *subject variability in structural measures* by relating functional measures to the thicknesses, areas and volumes of key cortical and subcortical structures as derived from the structural MRI scans [Glasser et al. 2013]. In this way, we can quantify to what extent different methods are able to capture key aspects of functional variability, and if there are meaningful relationships with other measures.

A more detailed overview of the data, and the tests we carry out here, can be found in Appendix G.

#### 3.2.1 Analyses

Both PROFUMO and spatial ICA were run at a dimensionality of 50, at which point the modes were reordered for visualisation and noise components—or, in the case of PROFUMO, modes eliminated by the Bayesian model complexity penalties— were removed. Even on the extensive and high-quality HCP data, PROFUMO does not identify more than 50 PFMs: when run at higher dimensionalities, more PFMs are simply eliminated from the model. We discuss why PROFUMO is likely to be conservative in this regard in more detail later.

For the full HCP data, PROFUMO therefore infers the posterior over approximately 25,000,000,000 parameters (1,000 subjects, 100,000 grayordinates, 50 modes, 5 parameters per grayordinate). In terms of computational requirements, this analysis took approximately 110 hours using 18 cores on a single compute node, and memory usage peaked at 350 GB.

Finally, note that subsequent figures display spatial maps on the cortical surface for simplicity and concision. However, all grayordinates (comprising approximately 60,000 cortical vertices and 30,000 subcortical voxels [Glasser et al. 2013]) were used in all analyses.

#### 3.2.2 Overview of the PFM spatial model

To begin with, in Figures 2 and 3 we show examples of the group- and subject-level spatial maps for four PFMs in order to demonstrate the richness of information contained within the PFM model. We do this to emphasise that PROFUMO is able to identify PFMs with strong spatial relationships with one another (in terms of overlap and anti-correlations), while at the same time being able to identify complex, subject-specific reorganisations of the group templates.

**Figure 2:**
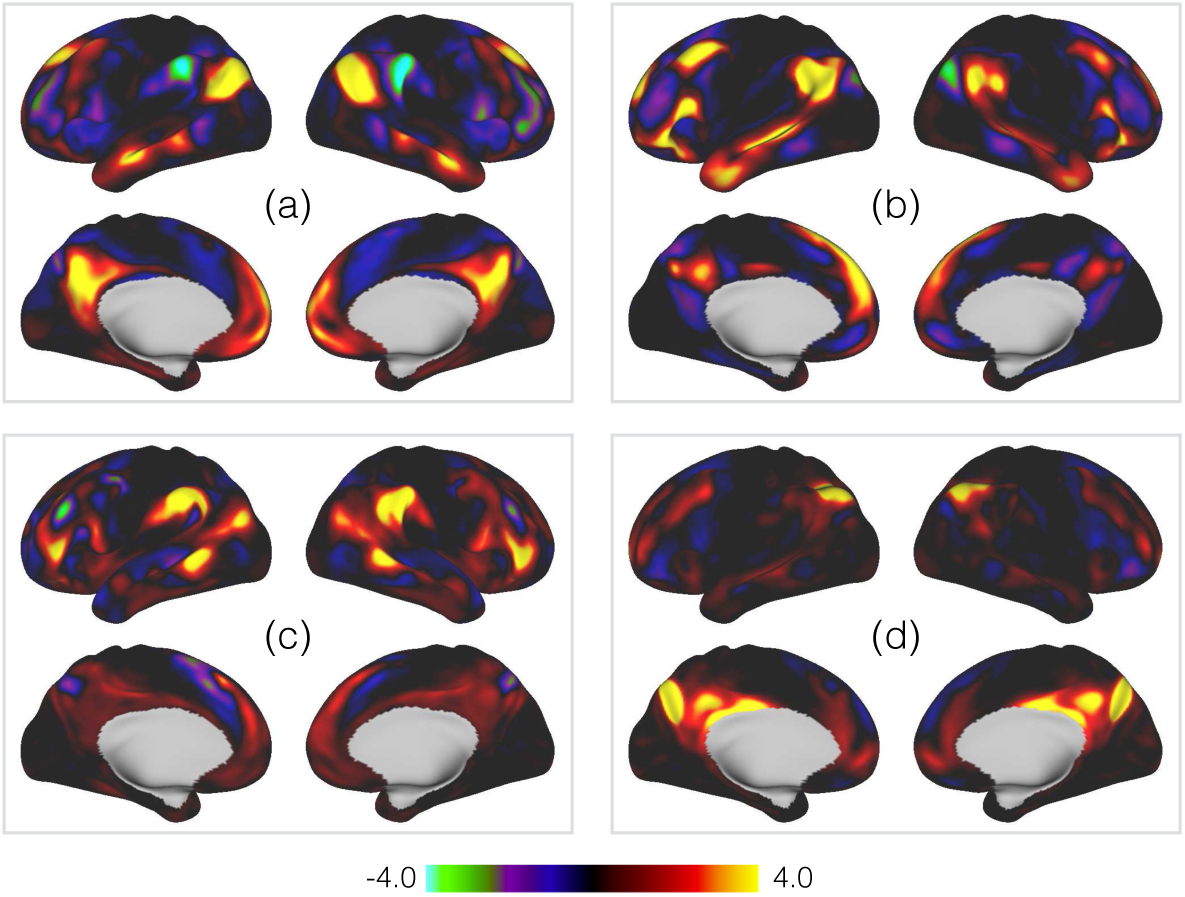
Group-level spatial maps for four example PFMs, as inferred from the HCP data. The PFMs are **(a)** the default mode network (DMN) [Shulman et al. 1997; Raichle et al. 2001; Greicius et al. 2003; Buckner et al. 2008]; **(b)** a mode described as a variant of the DMN by Braga and Buckner [2017]; **(c)** a mode with strong spatial anticorrelations with the DMN; and **(d)** the mode containing functional activity within POS2 [Glasser and Van Essen 2011].

**Figure 3:**
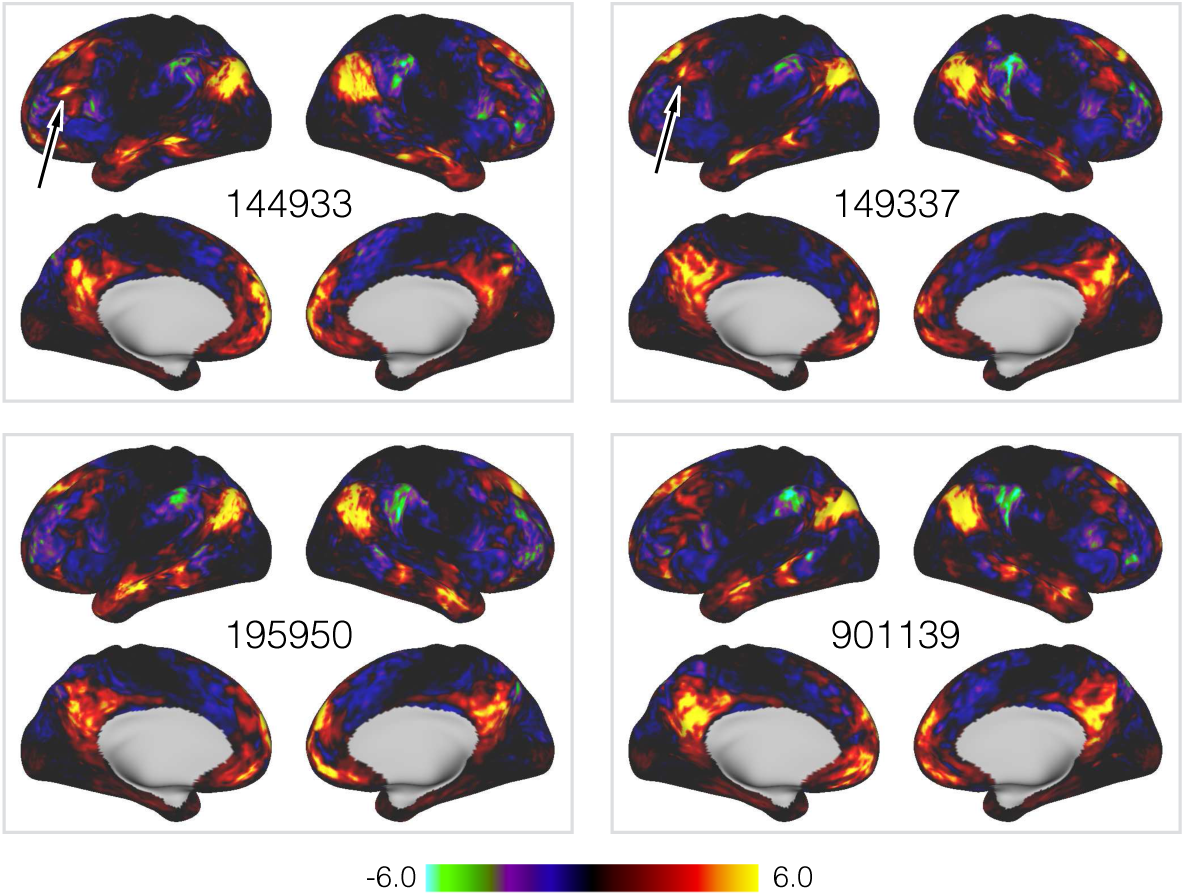
Subject-level equivalents of the default mode network shown in Panel of Figure 2.

The most striking feature of the subject maps in Figure 3 is simply how much variability relative to the group maps there is. These results are from data already aligned using surface-based registration driven by functional features, which arguably represent the current ‘gold-standard’ for warp-based registrations [Glasser et al. 2016b; Coalson et al. 2018]. Despite this, and as we and several others have demonstrated, there are pronounced differences between subjects, with both shifts in the relative location of functional regions over surprisingly large distances, and complex, non-homotopic splittings and reorganisations of the regions themselves. Furthermore, as highlighted in the figure, even though the PFM itself is large, there are several subject-specific features that are too small to be accurately represented at the typical spatial scale of parcellations applied to fMRI.

However, while the descriptions of modes in terms of the mean group- or subject-level spatial maps are familiar, a key advantage of the PFM framework is the more detailed group-level parameterisation. In other words, we can go beyond simply noting the degree of subject variability: we can now quantify it in detail on a per-mode level. In Figure 4 we again take the default mode network [Figures 2 and 3] as an example and plot the four key group-level spatial parameters: the probability that a given voxel belongs to the DMN, the mean and variability over subjects of the signal component of the DMN’s voxelwise weights, and the standard deviation of the spatial noise component. The information encoded by the mean weights is familiar, but the other parameters add novel and complementary information.

**Figure 4:**
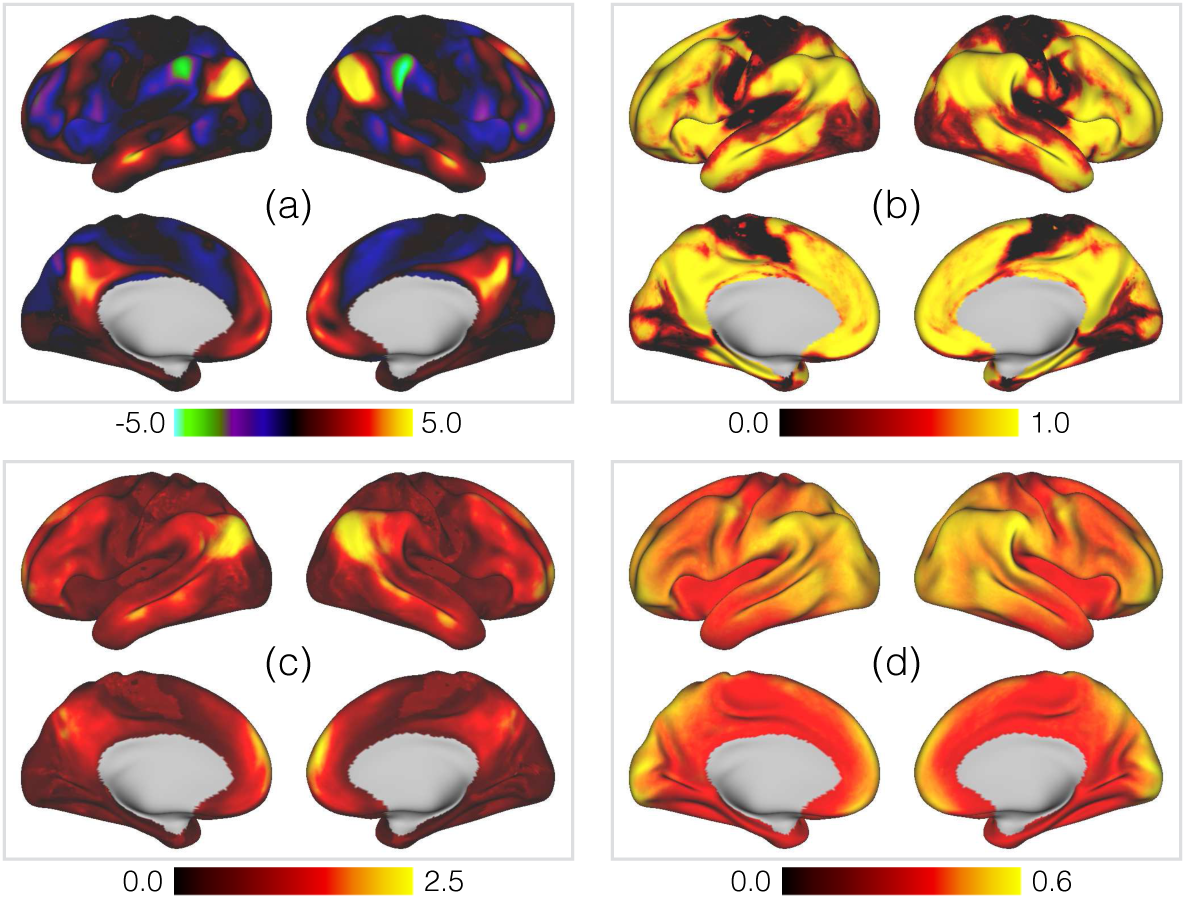
Example of the key group-level spatial parameters for the PFM representing the default mode network [Panel (a) of Figure 2 and Figure 3], as inferred from the HCP data. The parameters are the **(a)** posterior means of the signal of the signal component, *μ*_*vm*_; **(b)** posterior memberships, *π*_*vm*_; **(c)** posterior standard deviations of the signal component, *σ*_*vm*_; **(d)** posterior standard deviations of the noise component, *ζ*_*v*_.

For example, the memberships [Panel (b)] demonstrate that default mode activity is distributed over a surprisingly large area, with consistently detected activity across much of the lateral prefrontal cortex. This is an effect that has been captured by several recent, high-powered single-subject analyses [Gonzalez-Castillo et al. 2012; Laumann et al. 2015; Poldrack et al. 2015; Huth et al. 2016]. However, while the activity is widespread, it is also distinct: the areas of high and low probability are sharply delineated. Similarly, the standard deviations [Panel (c)] add extra information by telling us about variability in the size of the weights–that is, in the strength of the detected activity—and we can see that, in this instance, the activity in the inferior parietal lobule is much more variable in strength across subjects than that in the precuneus.

This detailed characterisation of non-homogeneous variability across the cortex is a key advantage of the more complex group-level model we have adopted, and we expand upon this in Figure 5. This summarises the membership probabilities and weight standard deviations across all modes. There is a clear pattern whereby association cortex contains more overlapping modes than sensory cortices [Panel (a)], and that the spatial weights are also more variable in association cortex [Panel (b)]—note how this is in agreement with the results of Mueller et al. [2013]. Finally, the uncertainty in the memberships themselves [Panel (c)] tells us about shifts in locations between subjects. For example, note the very clear area of variability in medial frontal cortex between SMA and pre-SMA [Johansen-Berg et al. 2004]. This metric is presumably particularly sensitive to this region because variability here tends to manifest itself as relatively simple anterior-posterior shifts of the SMA/pre-SMA boundary, whereas more complicated 2D rearrangements of overlapping PFMs are present elsewhere.

**Figure 5:**
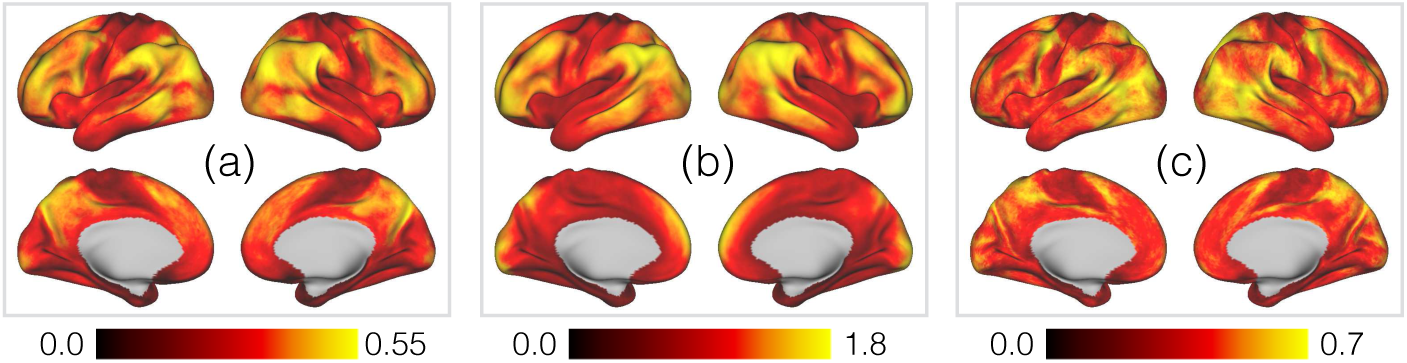
Summaries of the group-level spatial parameters encoding different aspects of variability across subjects. The panels are **(a)** mode overlap; **(b)** variability in mode strength; **(c)** variability in mode memberships.Mode overlap is defined as the posterior memberships averaged across all modes, 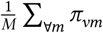Variability in mode strength is captured by the weighted average of the posterior st1andard deviations, (∑_∀*m*_ *π*_*vm*_*σ*_*vm*_)/(∑_∀*m*_ *π*_*vm*_). Finally, variability inmode memberships is given by the average entropy, in bits, of the membership distributions, 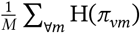

In summary, the PFM spatial model captures familiar group-level modes, and exhibits many of the complex subject-specific rearrangements already described in the literature. However, the key advantage is the way in which we have parameterised this model. Crucially, the richness of the group description allows us to make specific claims about the patterns of variability across the population that are ordinarily hard to tease apart.

#### 3.2.3 Comparison with spatial ICA

To begin with, we examine the performance of the different models in terms of their inference of the group-level spatial descriptions. In Figure 6 we plot the similarity between these group-level descriptions.

**Figure 6:**
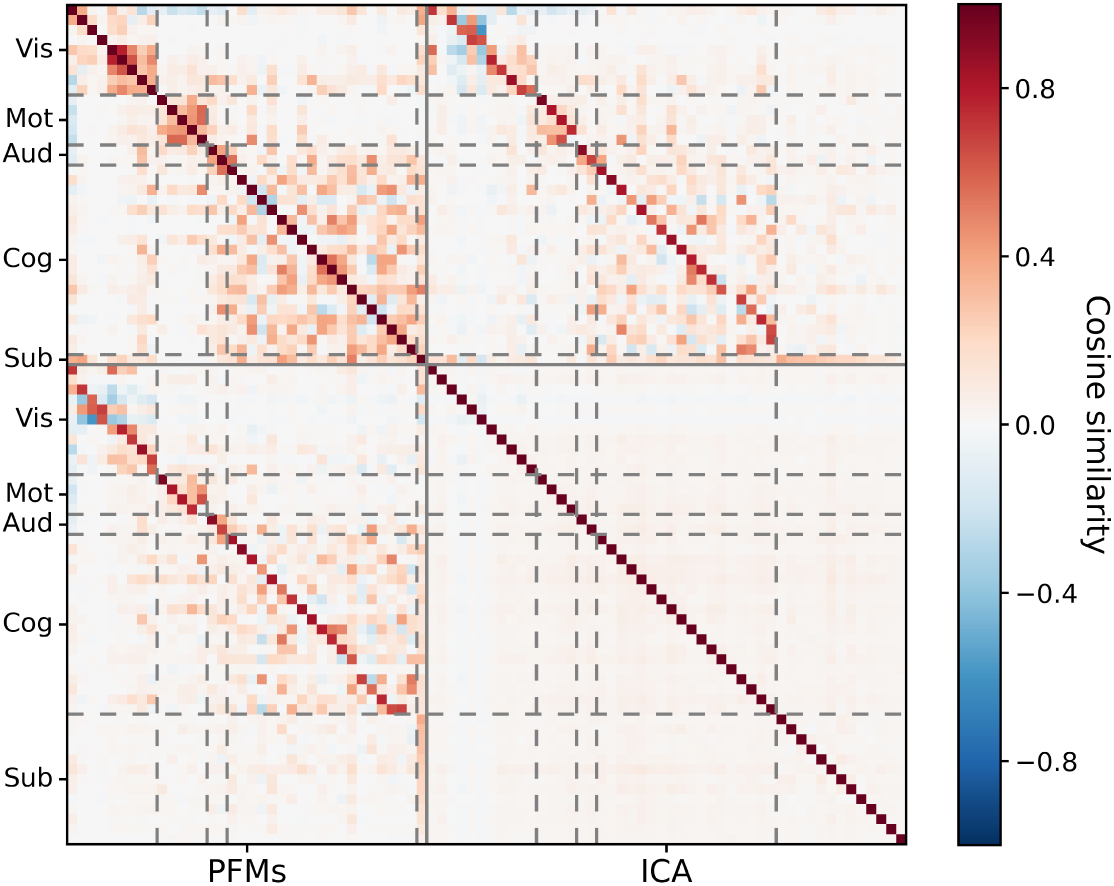
Spatial similarity between the sets of group-level spatial maps as inferred by PROFUMO and ICA. Modes were split into five categories and reordered: visual (Vis); motor (Mot); auditory (Aud); cognitive (Cog); and subcortical (Sub). This ordering is used for all subsequent sections.

There are several key points to note. Firstly, there are strong spatial correlations between the PFM maps, especially within the different categories. By way of contrast, the independence assumptions in spatial ICA preclude this. Secondly, PROFUMO is relatively conservative: it only infers 36 signal modes compared to the 48 found by ICA, and the difference is particularly pronounced in the subcortical regions. This subcortical difference is predominantly driven by the different signal properties of the HCP data between cortical and subcortical grayordinates, and the different data normalisation strategies the two algorithms use. The result is that ICA tends to find subcortical regions appearing in components without much cortical involvement, whereas PROFUMO tends to find subcortical regions appearing in components with cortical involvement. Finally, despite the above differences, there is fundamentally a strong relationship between the two sets of maps. Most cortical modes appear in both decompositions, and often look fairly similar; this is encouraging, as we do not expect a radically different patterns of functional connectivity at the group level given how many published methods have converged on similar descriptions.

#### 3.2.4 Properties of subject variability in spatial organisation

Given that the group-level descriptions are fairly similar between PFMs and sICA, the obvious question are to what extent does the extra group-level information in the PFM model regularise the subject-specific decompositions, and in what ways do the subject-specific maps diverge from the group-level representations? We deal with the former first, and in Figure 7 we look at that the consistency of the subject maps as inferred by PROFUMO and the ICA-DR pipeline. As expected given the regularisation from the more complex group-level priors, the PFM maps are much more consistent across subjects.

**Figure 7:**
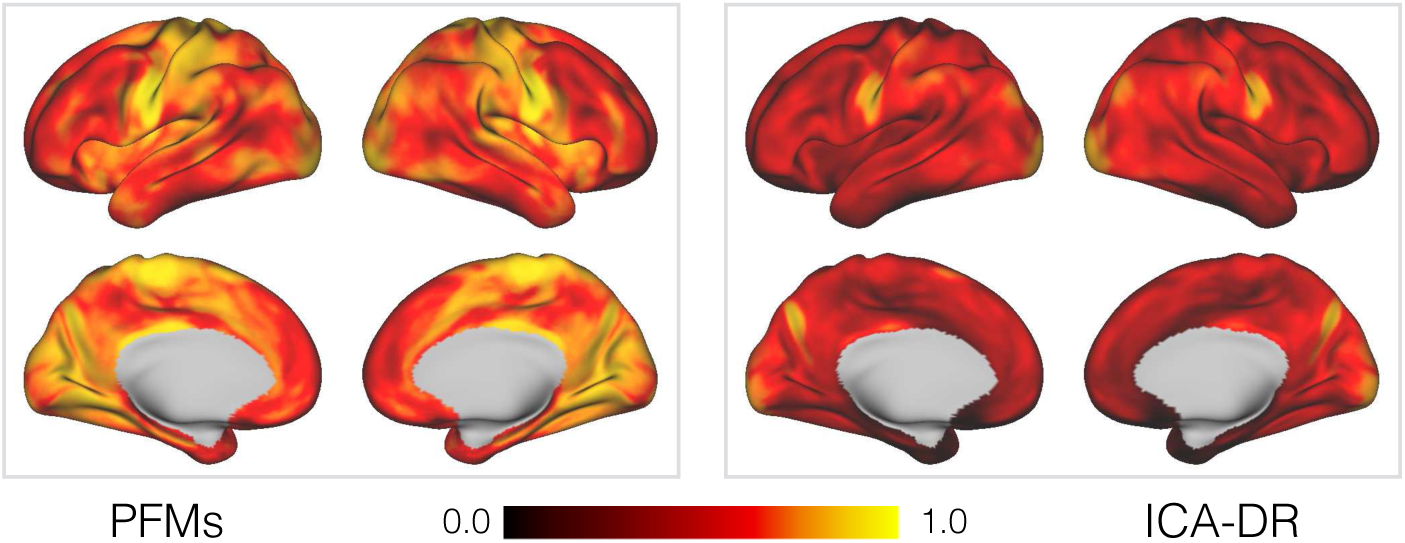
Similarity between the subject-specific spatial maps, for both PFMs and ICA-DR, as inferred from the HCP data. For each voxel and in every pair of subjects, we compute the Pearson correlation coefficient between the two *M*-dimensional vectors of mode weights. The maps shown here are the correlation coefficients averaged over every pair of subjects.

However, this increase in consistency could also be explained if the subject-specific PFM spatial maps were simply pushed closer to the group maps by the priors, thereby being less faithful to the ‘true’ patterns of functional connectivity at the subject level. While this does not appear to be the case for the exemplar subject maps, what we really want to quantify is whether they are capturing ‘interesting’ aspects of subject variability in spatial organisation. In other words, are the differences between the approaches meaningful, and do they make different predictions about the subjects themselves?

To investigate this, we use the fact that the HCP includes data from twins and siblings to investigate the influence of genetics and environment. We estimate the voxelwise broad-sense heritability of the subject-specific spatial maps we observe: in each voxel and each subject, we extract the vector of PFM or ICA map weights, and look to see if these weight vectors are more consistent in monozygotic than dizygotic twins (see Appendix G for full methodological details). The results of this analysis are shown in Figure 8.

**Figure 8:**
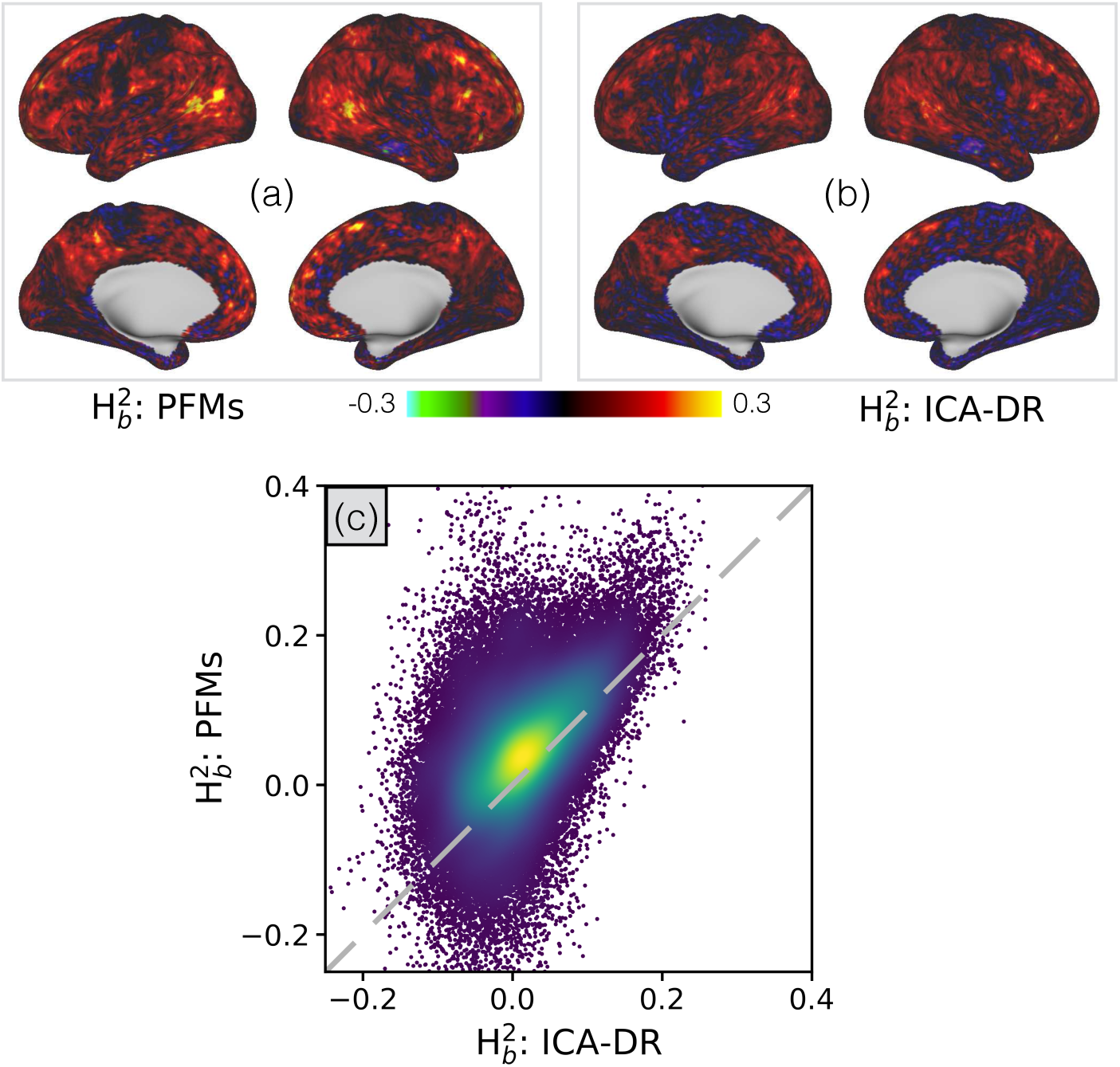
Analyses of the heritability of the subject-specific mode maps, for both PFMs and ICA-DR, as inferred from the HCP data. In **(a)** and **(b)** we display the voxelwise estimates of broad-sense heritability 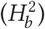and in **(c)** we compare the two as a scatter plot.

The results show a clear increase in heritability for the PFM spatial maps, suggesting that they are more sensitive to subject variability that we can attribute to genetic factors. Furthermore, this is not simply attributable to a reduction in noise or as the result of the priors pushing the subject maps closer to the group. While the PFM maps are more consistent across subjects than ICA-DR [Figure 7], the heritability relates to the difference in consistency between monozygotic and dizygotic twins and, as such, a global increase in consistency is not enough to explain the increased heritability.

We can also gain further insights into this observation by utilising the HCP’s retest data. 46 subjects underwent the full HCP imaging and behavioural testing protocol twice, of which there is full rfMRI data from 42. This allows us to examine how the algorithms perform on the hitherto unseen retest scans. The group-level representations from the full data (i.e. the ICA spatial maps, and the group-level PFM posteriors) were used to derive new subject maps from the independently acquired retest data.

In Figure 9 we compare subject-specific realisations of the language mode as derived by PROFUMO and the ICA-DR pipeline. This particular mode was chosen because a characteristic split in area 55b in some subjects was reported and examined in some detail by Glasser et al. [2016a]. In terms of a comparison between PROFUMO and ICA-DR, both are clearly sensitive to the same gross re-organisations that occur. For example, both can detect the rearrangement of area 55b in the original and retest data for the subject shown here. However, the most marked difference is in the noise-level and appearance of anticorrelations. Relative to ICA-DR, the PFMs show much reduced background noise in regions not associated with the networks, and do not exhibit anticorrelations (indicated by negative weights, shown in blue) tightly interposed between positive weights. This is presumably a simple consequence of dual regression’s inability to separate signal from noise, as we discussed in the section on noise modelling. By way of contrast, the information encoded by the group-level parameters in the PFM model suppresses the background noise in regions that are not part of the language network, but in a way that does not preclude inferring complicated rearrangements of functional regions.

**Figure 9:**
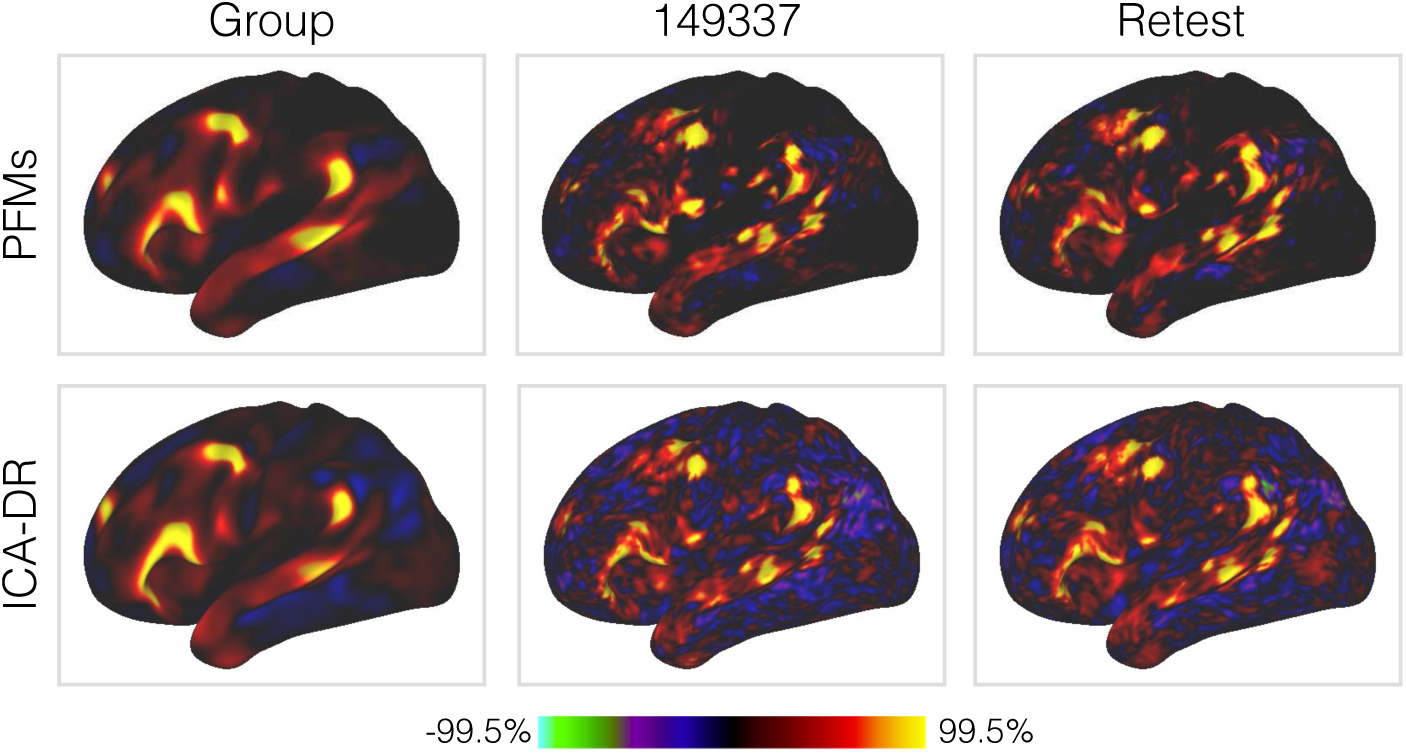
Example spatial maps for the language mode, for both PFMs and ICA-DR, as inferred from the full HCP data and the HCP retest data for subject 149337. Only the left lateral surface is shown.

To assess the reliability of the different decompositions on the retest data more quantitatively, we assess the specificity of the inferred spatial maps as ‘fingerprints’ that uniquely identify different subjects [Finn et al. 2015; Horien et al. 2019]. This is shown in Figure 10.

**Figure 10:**
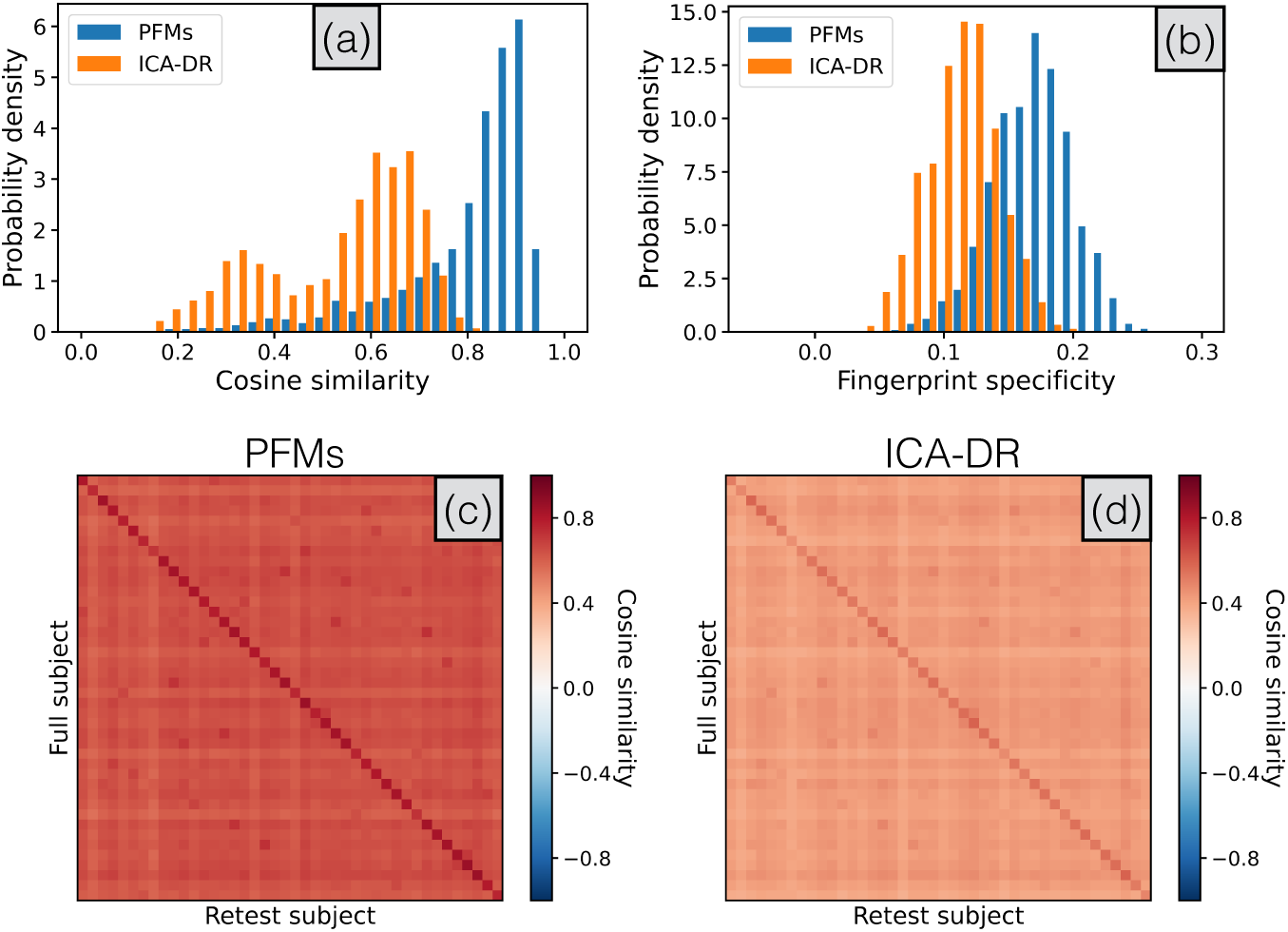
Specificity of the subject-level spatial maps as inferred from both the original and retest HCP data by PROFUMO and ICA-DR. The group results from the full data are used to derive subject-specific spatial maps in the unseen retest data. In **(a)** we show the similarity of the inferred maps in the same subject, seperately for each mode. In **(b)** we calculate the fingerprint specificity, or how much more similar the maps in the same subject are as compared to maps from non-matching pairs of subjects, averaged over modes. This is equivalent to the difference between the diagonal and the off-diagonal elements (calculated for each column separately) in the full simmilarity matrices as shown in **(c)** and **(d)**.

Firstly, we compute the spatial similarity between the new subject-specific spatial maps from the retest data, and the original set from the full data, for every pair of subjects. We pool these retest results over all modes and subjects, and this is shown in Panel (a). Again, the subject-specific PFM maps are much more consistent across the two acquisitions.

Secondly, we assess whether this leads to more specific fingerprints. In Panel (b) we show that the fingerprint specificity (i.e. the amount by which the two sets of maps from the same subject are more similar than paired maps from different subjects) is also higher for the PFMs. In other words, not only are the maps generally more consistent across subjects, but there is an increase in subject specificity above and beyond this effect.

In summary, the comparisons with ICA-DR have demonstrated that while the group-level descriptions are similar, the more complex hierarchical modelling in PROFUMO allows us to infer spatial maps that are more consistent—on both the original data and the held-out retest data—as well as being more specific and capturing more informative aspects of cross-subject variability.

#### 3.2.5 Overview of the PFM temporal model

Here, we briefly give a summary of the key temporal parameters—that is, the amplitudes and the functional coupling between modes—as inferred by PROFUMO on the HCP data. Note again that these are new parameters: in other words, it was only possible to investigate these in a post-hoc fashion based on the previous PFM model. Firstly, in Figure 11 we plot the cross-subject correlations between the mode amplitudes, as captured by the ***Σ***_***h***_ parameter. Encouragingly, we see a clear replication of the results of Bijsterbosch et al. [2017], who reported strong correlations between the amplitudes of sensorimotor modes, as well as between cognitive modes, but relatively weak correlations across the two categories. However, the crucial difference between this result and the original observation is that this behaviour was initially demonstrated from a purely post-hoc analysis of the ICA-DR results, whereas it is explicitly parameterised and inferred within the PFMs model. What this means is that this knowledge of the systematic relationship between mode amplitudes is available *during* inference, and it is therefore naturally incorporated as an extra factor regularising the subject-specific decompositions.

**Figure 11:**
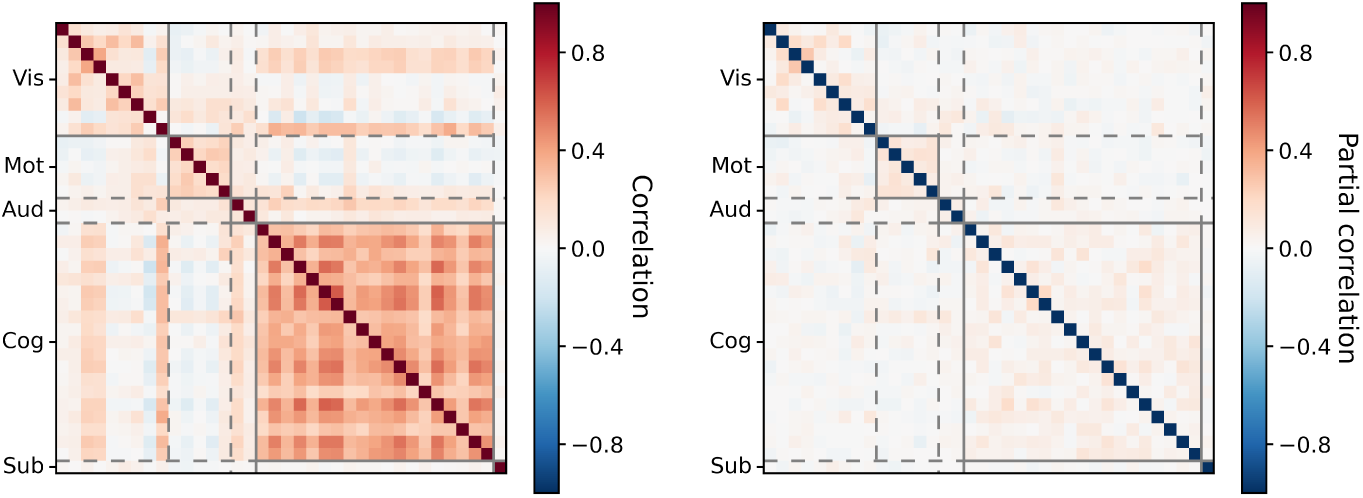
Cross-subject relationships between the amplitudes of the PFMs, as inferred from the HCP data. For visualisation purposes, we display the posterior precision matrix, ***Σ***_***h***_, after transforming it to both full and partial correlation coefficients.

Secondly, in Figure 12 we plot the PFM functional coupling parameters, ***β*** and ***α*** ^(*s*)^ (these represent the group- and subject-level temporal network matrices respectively). What is striking is how weak the functional coupling is between modes in the group-level network matrix (netmat), especially given that we have an explicit hierarchical model to allow for just these interactions. This is not trivial to explain away as a spatial effect either: despite the fact that these interactions are more similar to what we would expect from temporal ICA, the PFM spatial maps are similar to those inferred by spatial ICA which typically infers strong functional coupling between modes. We quantify the implications of this different view on functional coupling from the PFM model in the following section.

**Figure 12:**
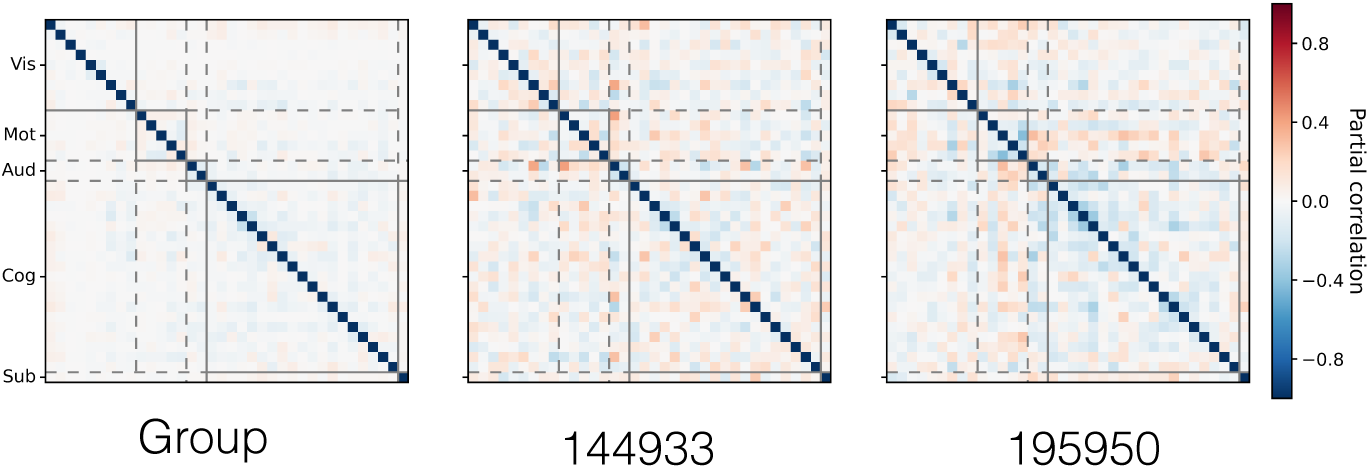
Group- and subject-level functional coupling between the PFMs, as inferred from the HCP data. For visualisation purposes, we display the posterior parameters ***β*** (group-level) and ***α***^(*s*)^ (subject-level) as partial correlation coefficients. As in Figures 3 and 9, subject 149337 is chosen as the exemplar.

#### 3.2.6 Multivariate relationships with behavioural variables

How then, are we to interpret the differences between the PFM and ICA-DR approaches? Do they simply represent a different trade-off between sensitivity and specificity in the spatial and temporal domains, or are they telling us something fundamentally different about brain activity?

To probe this further, we performed a series of multivariate analyses to investigate the different ways in which the two models encode cross-subject information. Like in Smith et al. [2015], canonical correlation analysis (CCA)—a multivariate analysis technique used to find the linear relationships between sets of variables [Hotelling 1936]—was used to summarise the key correspondences (see Appendix G for methodological details). Furthermore, as some sets exhibit more than one strong linear relationship, we use the RV coefficient [Robert and Escoufier 1976] to give a principled summary of the multivariate information reported by the CCA. In Figure 13, we examine the full set of pairwise relationships between the behavioural and structural variables from the HCP, and the spatial maps, amplitudes and network matrices from both PROFUMO and ICA-DR.

**Figure 13:**
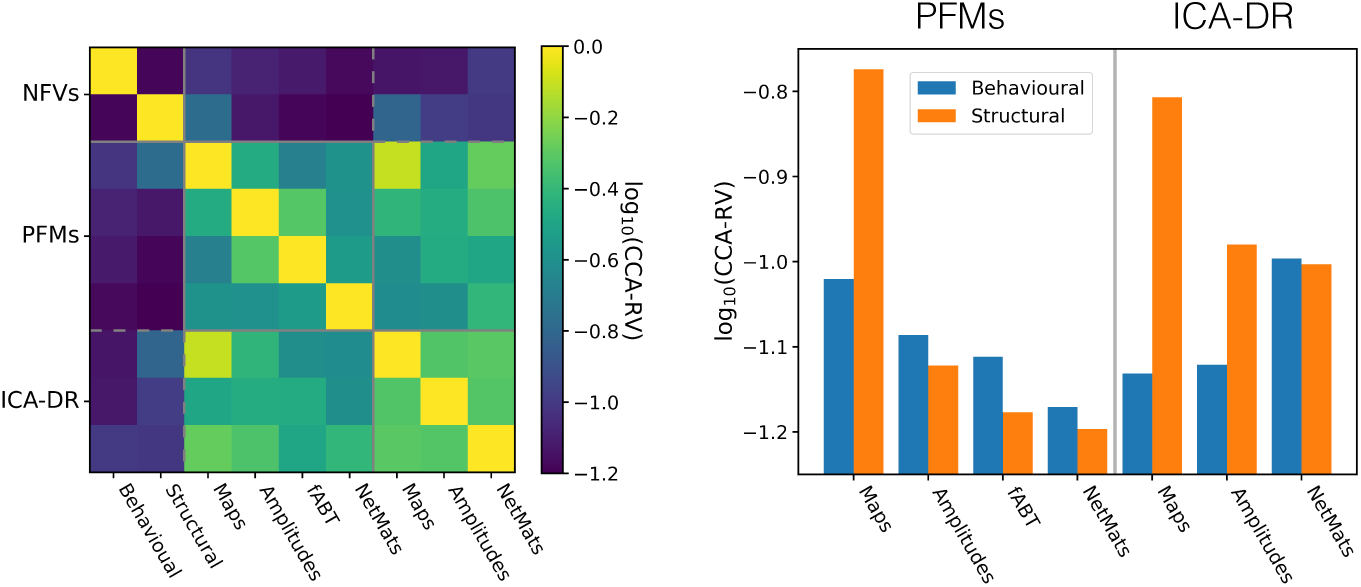
Relationships between the cross-subject information encoded by different analyses. The non-functional variables (NFVs) have been separated into variables from the HCP’s battery of behavioural tests, and variables derived from structural MRI relating to brain size and morphology. On the left we plot the log RV coefficient calculated between the subspaces of the top ten CCA components as calculated between every pair of sets of variables, and on the right we reproduce the relationships with the non-functional variables (i.e. the top two rows / two leftmost columns) as a bar chart for ease of visualisation. Higher values of the RV coefficient indicate that more similar cross-subject information is being captured.

There are several key results we can glean from this analysis. Firstly, the cross-subject information captured by the different aspects of the PFM model is relatively distinct. Comparing the similarity between the PFM measures with those for the ICA-DR variables (i.e. the on-diagonal blocks), we can see that the scores are typically lower for the PFMs. In other words, the temporal measures derived from the PFMs carry relatively different information from the spatial measures about the subjects themselves, at least compared to their ICA-DR equivalents.

Secondly, if we examine the relationships with the behavioural and structural measures in the bar graph on the right, there are several striking differences between the methods. As we would expect from our earlier analyses, the PFM spatial maps are the best predictors of structural variables. They are also good predictors of the behavioural variables, though slightly less so than the ICA-DR netmats. However, the stories for the temporal information are very different. The PFM amplitudes, fABT and netmats are relatively poor predictors of both behavioural and structural variables, though, intriguingly, they are better predictors of behaviour than structure. By way of contrast, the ICA-DR amplitudes and netmats are better behavioural predictors, though surprisingly they are also good predictors of structure (e.g. one can predict the sizes and thicknesses of cortical areas better than behavioural measures from the ICA-DR amplitudes).

Given the simulation results, the interpretation is relatively straightforward: the ICA-DR pipeline contains inherent biases that conflate spatial and temporal information. Furthermore, even though we do not explicitly test it here, it is interesting to note that using the thresholded version of dual regression to correct this bias also reduces the correlation between temporal netmats and behaviour [Bijsterbosch et al. 2019]. In other words, and consistent with the results on simulated data, thresholded dual regression is an improvement on ICA-DR but is less accurate than the full PROFUMO model. The question that remains however, is what information, if any, is the PFM temporal model capturing if not the trait-like behavioural variables examined here?

#### 3.2.7 Summary

Given the full set of results presented on the HCP data, the implication is that the PFMs, by virtue of the improved spatial modelling in particular, are better able to capture interesting information about cross-subject variability in spatial organisation. However, this does not address the relative lack of information encoded in the various temporal measures that PFMs capture. We address this point using another data set in the following section.

### 3.3 Active-state data

Given the way that subject variability in spatial and temporal features simultaneously co-varies with a wide range of non-imaging derived subject measures, it is very challenging to conclusively disambiguate them from studies like the HCP. However, if we manipulate the functional connectivity at the subject level, for example by changing the cognitive state [Shirer et al. 2011; Krienen et al. 2014; Vanderwal et al. 2017; Gratton et al. 2018; Kieliba et al. 2019; Salehi et al. 2020], then we can begin to examine temporal differences in more detail. Crucially, by looking at multiple conditions for the same subject we essentially eliminate the influence of structural variability from the functional data.

To do this, we use a dataset collected where subjects were scanned when in different active states—these are induced by performing simple, continuous tasks in the scanner, of which rest (i.e. eyes-open fixation) is just one [Duff et al. 2018; Kieliba et al. 2019; Sala-Llonch et al. 2019]. There are five runs for every subject, each collected under different steady-state conditions: a standard resting-state acquisition (`Rest`); a finger-tapping based motor task (`Mot`); a passive visual condition (`Vis`); an independent combination of the visual stimulus and motor task (`V-M`); and a condition where the specifics of the motor task changed based on the visual stimulus (`V+M`). A more detailed descriptions of the tasks and data itself can be found in Kieliba et al. [2019]. Furthermore, this dataset offers a validation of our method on data acquired using a more conventional sequence and scan duration than the HCP, with fewer subjects, shorter scan durations, and all analyses performed on volumetric rather than surface-based data.

#### 3.3.1 Analyses

As per the modelling assumptions, PROFUMO infers one consensus spatial map per subject, but a separate set of time courses per run. We choose to infer run-specific temporal precision matrices, ***α***^(*sr*)^, with aconsistent group-level hyperprior, ***β***, which is shared across all conditions. Note that we could have chosen to use condition-specific group-level priors, 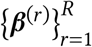, but this has the side-effect of invalidating the assumptions behind any subject-level statistics where we compare between conditions. In short, it reduces the cross-subject, within-condition variance which invalidates the typical null hypothesis we use. We leave the problem of performing statistical inference on these types of models for future investigations.

We infer 30 modes for both PROFUMO and ICA-DR, which again seems to be close to the upper limit for PROFUMO on this relatively small dataset. Again, artefactual modes were eliminated and those remaining were reordered for visualisation. In terms of computational requirements, the PROFUMO analysis took approximately 12 hours using 15 cores on a single compute node, and memory usage peaked at 25 GB. Compared to the HCP analysis, the demands are higher than expected given the number of subjects for two reasons: firstly, the volumetric analysis contains over twice as many voxels as grayordinates; secondly, we do not do within-subject data reduction for this analysis.

For the ICA-DR pipeline, we use MELODIC [Beckmann and Smith 2004; Beckmann et al. 2005] to infer a set of group maps, followed by dual regression to generate the run-specific time courses.

#### 3.3.2 Overview of the PFM model

In Figure 14 we show the group-level properties of the default mode as inferred from this data set. This is directly comparable with Figure 4, and simply demonstrates that we are able to infer similar summaries of the mode itself, and heterogeneous variability, from fourteen subjects rather than one thousand.

**Figure 14:**
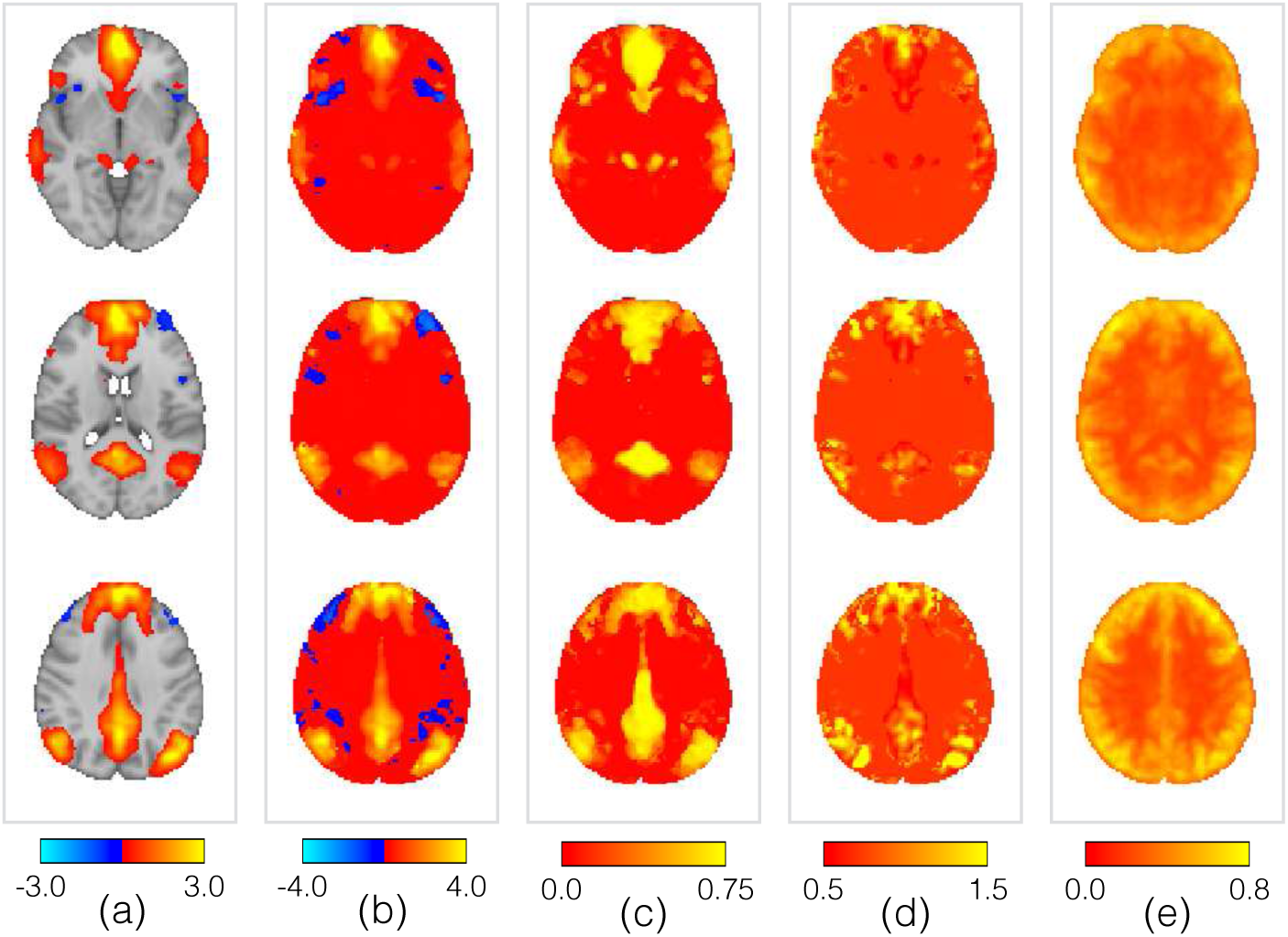
Example of the key group-level spatial parameters for the PFM representing the default mode network, as inferred from the active-state data. The parameters are as per Figure 4, along with the group map. The panels are the (a) group map; **(b)** posterior means, *μ*_*vm*_; **(c)** posterior memberships, *π_vm_*; **(d)** posterior standard deviations, *σ_vm_*; **(e)** posterior noise standard deviations, *ζ*_*v*_.

In Figure 15, we demonstrate some of the properties of the inferred time courses from the PFMs. This data is more challenging than the HCP in that the runs are shorter, and the data has not benefited from resampling onto the cortical surface. Nevertheless, the HRF-based prior constraint results in a temporally smooth timecourse, which we are able to cleanly separate from the high-frequency noise which contaminates them. Furthermore, this is stable when we undo the temporal blurring that the HRF induces, with straightforward estimation of the underlying ‘neural’ process via whitening with respect to the autocorrelation induced by the HRF.

**Figure 15:**
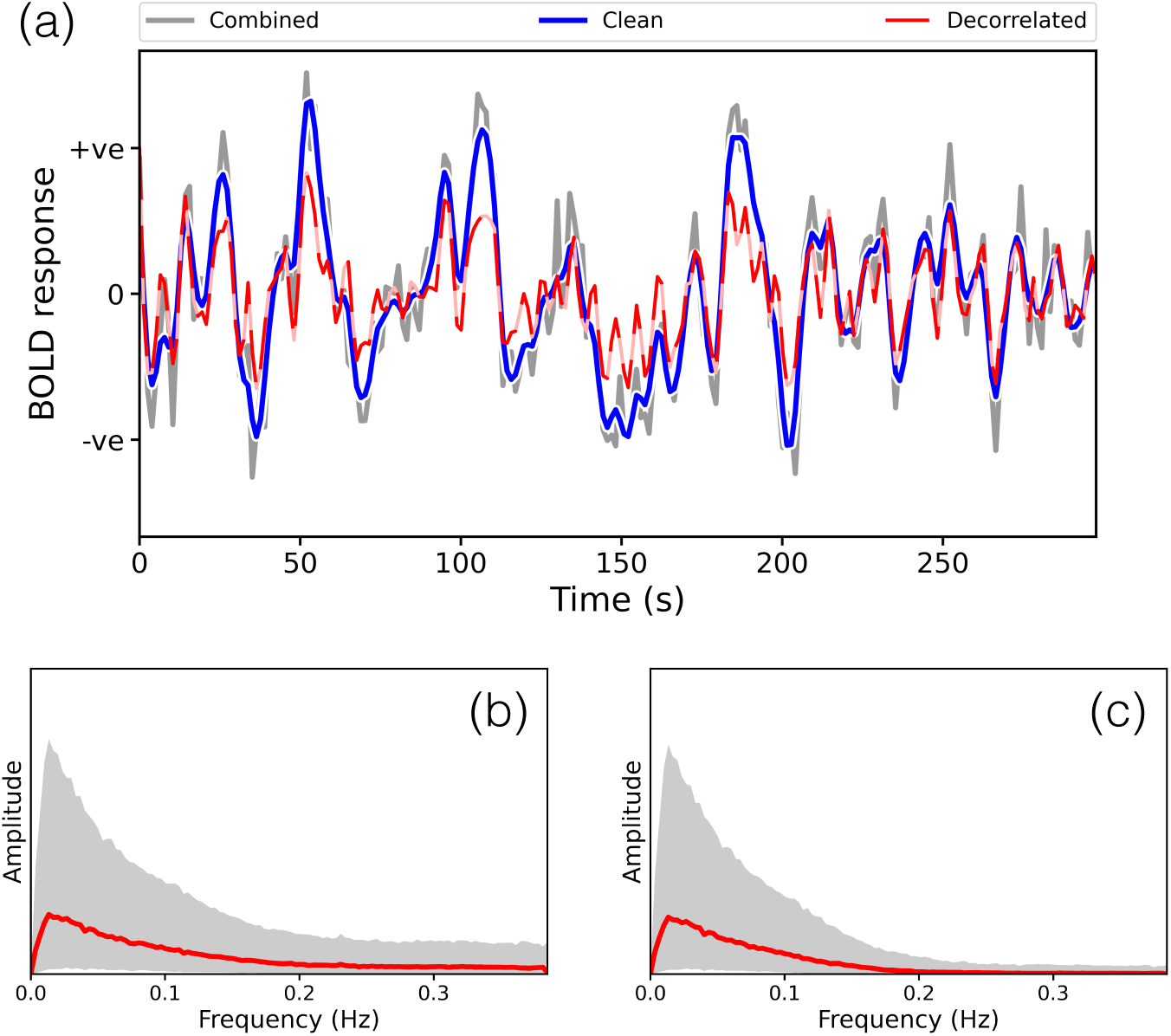
Example PFM time courses, and observed frequency content, from the active-state data. **Panel (a):** Example time course for one mode in one run.’Combined’ refers to the time course which includes the noise terms (***A***^(*sr*)^ = ***B***^(*sr*)^ + ***ξ***^(*sr*)^), ‘clean’ refers to the BOLD portion specifically (***B***^(*sr*)^), while ‘decorrelated’ refers to the clean time course after correcting for the temporal autocorrelation induced by the HRF 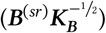. **Panels (b) & (c):** Frequency content of the combined and clean time courses respectively, pooled over all runs and subjects. The magnitude of the DFT coefficients are calculated for each time course, and for visualisation purposes, we fit a gamma distribution to the histogram of observed magnitudes for each frequency bin. The mode of this distribution is plotted in red, and the grey region represents the 95 % highest density interval [Kruschke 2014].

Finally, in Figure 16, we display examples of the network matrices to illustrate the typical patterns of, and subject variability in, the functional coupling between PFMs. Interestingly, in this data, PROFUMO infers PFMs with much stronger functional coupling between them than at the run level from the HCP data.

**Figure 16:**
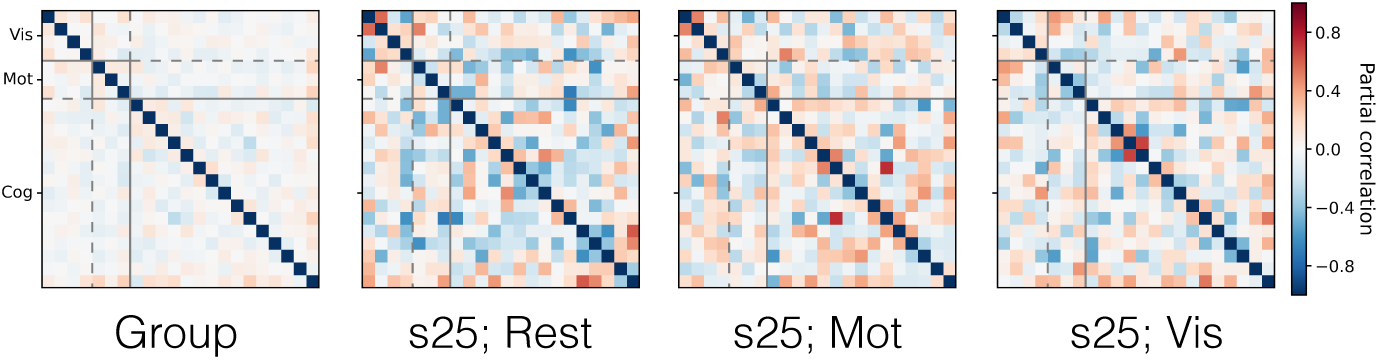
Example PFM network matrices, capturing the functional coupling between the mode timeseries. We display the group network matrix alongside the network matrices from subject 25 in the rest, motor and visual conditions. As in Figure 12, we display the posterior precision matrices (i.e. for the ***β*** group level and ***α***^(*sr*)^ at the run level) as partial correlations. Modes were split into three categories and reordered for visualisation of the network matrices: visual (Vis); motor (Mot); and cognitive (Cog).

#### 3.3.3 Comparison with ICA-DR

One would hope that the PFM model allows us to more accurately infer the true functional coupling between modes. To begin with, we look at the relationships between the condition-specific network matrices as inferred by PROFUMO and ICA-DR. These are shown in Figure 17. While the PFM network matrices are less consistent between conditions and subjects than their ICA-DR counterparts, there is some indication that there is condition-specific modulation across subjects (as indicated by the block diagonal). By way of contrast, the ICA-DR network matrices are dominated by the subjects themselves (i.e. the multiple strong off-diagonal lines in the ICA-DR plot), with no real indication of condition-specific modulations.

**Figure 17:**
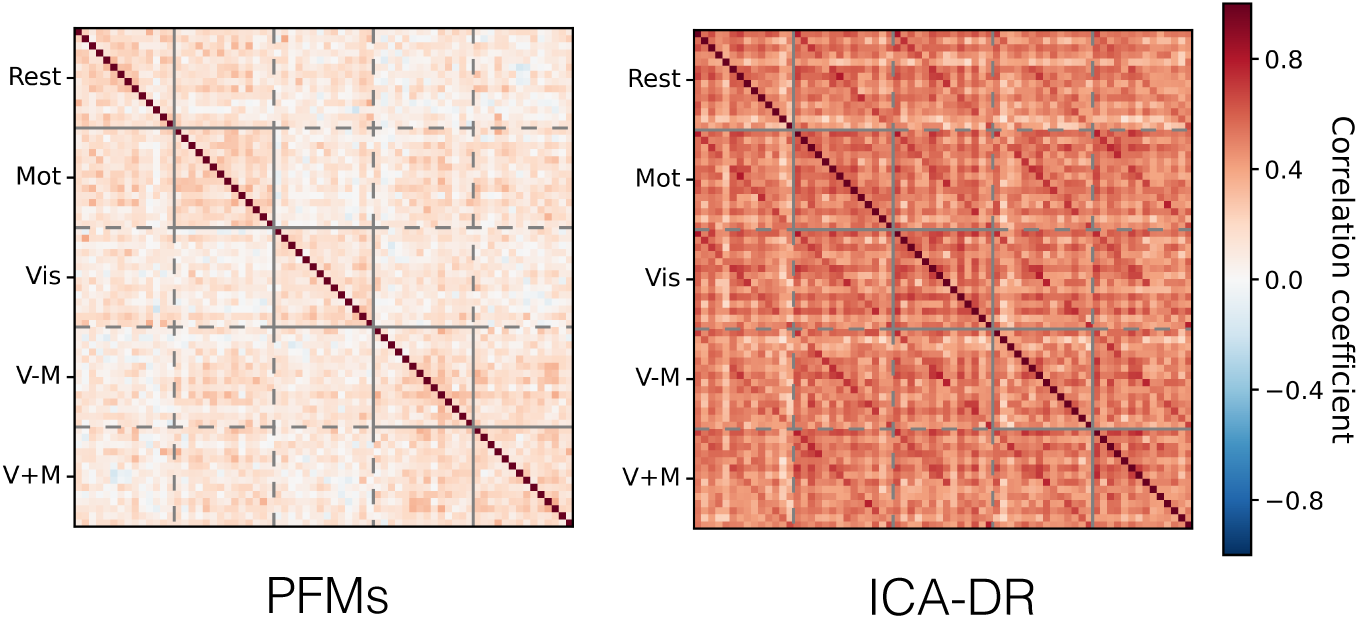
Correlations between the network matrices, for both PFMs and ICA-DR, as inferred from the active-state data. The network matrices are grouped by condition, and the subjects have a consistent ordering within each block. Correlation is the Pearson correlation coefficient between the unwrapped upper-triangle of the network matrices.

In summary, ICA-DR computes netmats that are more similar within subjects than they are within conditions across subjects. By way of contrast, PROFUMO infers netmats that are somewhat more similar within conditions than within subjects. Again, this suggests that the different models for subject variability in spatial organisation have a profound influence on downstream estimates of functional connectivity.

Next, we test whether the different conditions induce focal changes to the between-mode patterns of functional connectivity. The results of a statistical analysis that looks for modulations at the level of individual network matrix edges are shown in Figure 18. Both the PFM and ICA-DR pipelines detect changes in the coupling of visual regions induced by the visual stimulus, and it appears they both have similar sensitivity to the changes in coupling induced by the changes in cognitive state. There are some differences between the methods: for example, the visual changes detected by PROFUMO are more consistent across the three conditions with visual stimuli than for ICA-DR. Similarly, the types of changes for the combined visuo-motor condition are somewhat different, with ICA-DR finding changes in amplitude predominantly, whereas there are more changes in coupling for PROFUMO.

**Figure 18:**
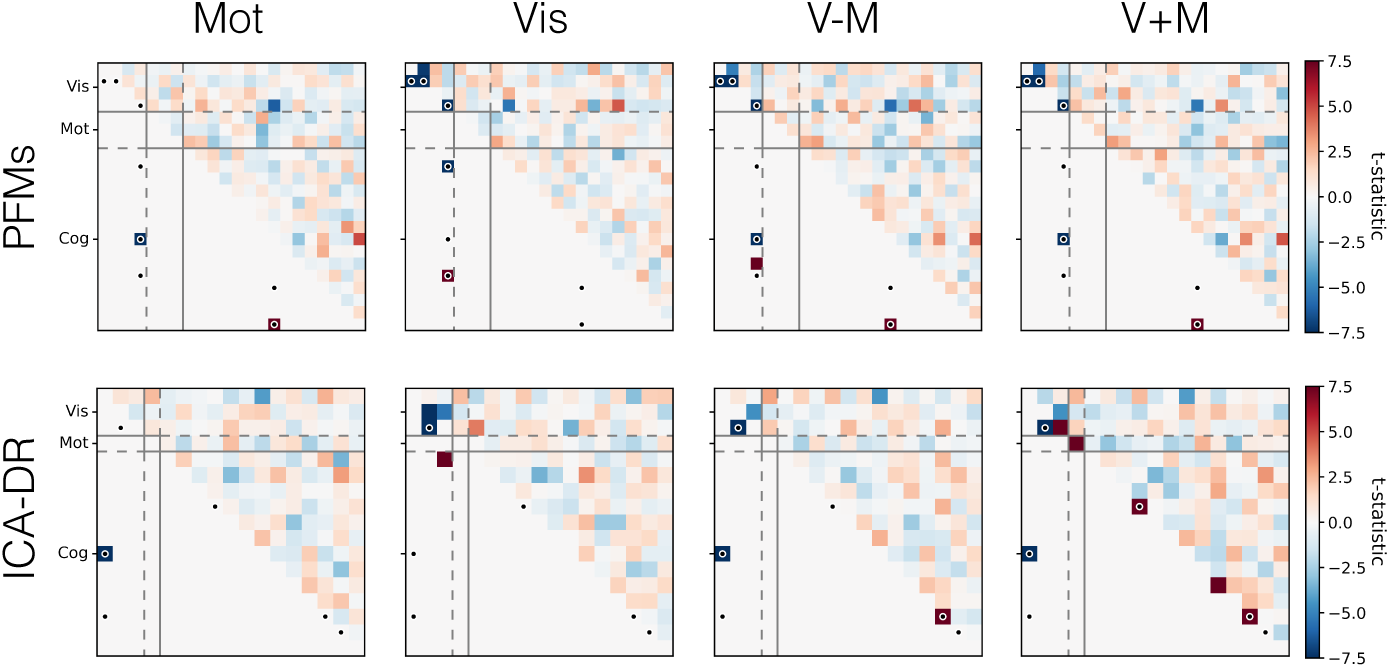
Changes in between-mode functional connectivity as induced by different active states relative to the rest condition. The raw difference between the group mean network matrix during the active condition and during rest is shown above the diagonal, and any significant changes (p<0.05) are highlighted by blue or red squares, for increases and decreases in coupling respectively, below the diagonal. Changes in amplitudes are shown on the diagonal. All tests were family-wise error corrected and computed using the accelerated permutation inference in PALM [Winkler et al. 2014; 2016]. The black dots denote elements that were significant under an f-test over all contrasts. As per Figure 16, modes were split into three categories and reordered for visualisation of the network matrices: visual (Vis); motor (Mot); and cognitive (Cog).

However, the results are fundamentally fairly similar and the numbers of edges that exhibit significant changes is relatively low—and, perhaps, lower than we might expect given the strong manipulations of cognitive state^12^—suggesting that the statistical power might be the limiting factor here, especially given that there are only 14 subjects included in this analysis. Finally therefore, we do one further set of tests to probe whether the multivariate information in the network matrices and amplitudes captures condition-specific information. Repeating the analysis of Sala-Llonch et al. [2019], we investigate whether a support vector machine (SVM) can be trained to distinguish between network matrices from different conditions. The accuracy of the SVM classification is tested using a leave-one-subject-out cross-validation framework [Varoquaux et al. 2017], of which we provide more methodological details in Appendix H.

As well as comparing PROFUMO and ICA-DR in this way, we additionally examine the effect that the hæmodynamic model has on the temporal information that we infer. In other words, can the changes to estimates of functional connectivity be attributed to the advanced spatial modelling alone, or does the regularisation in the time domain improve our estimates too? As well as the explicitly inferred PFM network matrices, we do a post-hoc estimation of the temporal network matrices based on both the BOLD time courses and the combined time courses (i.e. ***A***^(*sr*)^, which includes both the BOLD and noise time courses) to assess what, if any, effect the modelling hierarchy has.

The results from the SVM analysis are presented in Figure 19. The SVM achieves a significantly better classification accuracy when trained on the PFM netmats, as opposed to those estimated by ICA-DR. Again, this suggests that by correcting for subject variability in spatial organisation the PFM framework allows us to estimate state-induced changes in functional coupling with greater fidelity. By way of contrast, the conflation of spatial and temporal information by ICA-DR masks these more subtle state-related changes in functional coupling. Finally, there appears to be a distinct performance improvement when using the inferred PFM network matrices, suggesting that the hierarchical temporal modelling is advantageous and that we are not discarding relevant information by focusing on the predominantly low-frequency HRF-derived time courses.

**Figure 19:**
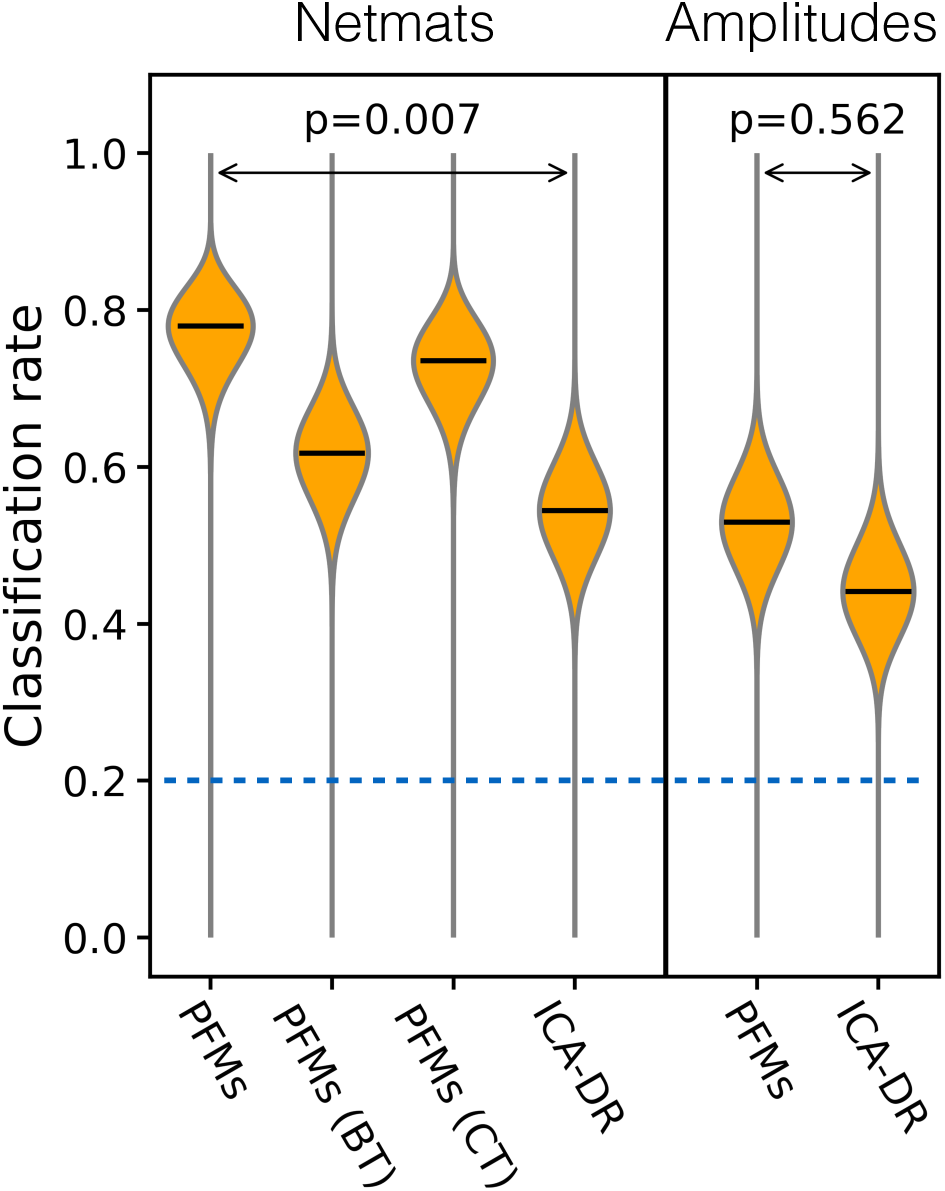
Posterior classification rates for a multi-class SVM trained to distinguish between the different active-state conditions. The results on the left are when the off-diagonal elements of the network matrices are fed in, and the results on the right are when the amplitudes are used as features. Posterior densities are based on the number of correct and incorrect classifications out of the full set of 70 tests (14 subjects; 5 conditions), combined with Haldane’s uninformative beta prior [Haldane 1932]. The modes of the distributions are shown by the black bars, and the chance level is shown by the dashed blue line. The two p-values are calculated via McNemar’s test (mid-p variant) and Bonferroni corrected [Fagerland et al. 2013]. For the PFM netmats, the variants are: *PFMs*: network matrices inferred as part of the PFM model, ***α***^(*sr*)^ *PFM (BT)*: network matrices estimated as the partial correlations between the PFM BOLD time courses ***B***^(*sr*)^. *PFM (CT)*: network matrices estimated as the partial correlations between the combined time courses ***A***^(*sr*)^ = ***B***^(*sr*)^ + ***ξ***^(*sr*)^.

## 4 Discussion

In summary, the results presented above demonstrate three key attributes of PROFUMO. Firstly, the algorithm is applicable to modern, large-scale data, whereby it is exquisitely sensitive to cross-subject variability in spatial organisation. Secondly, the joint inference framework allows estimation of subject variability in temporal features that does not appear to be confounded by spatial differences, which at times leads to a radically different view of functional connectivity. Finally, the implication of these results is that after accounting for spatial variability, the functional coupling between modes is much more reflective of current cognitive state rather than trait-like qualities.

Furthermore, we have shown that there is significant value added in terms of interpretability from the practitioner’s point of view in using models of this form. To give a few concrete examples, we can only make the claims pertaining to the dissociation of non-homogeneous spatial variability—as illustrated in Figure 5—if we can both consistently identify equivalent functional systems across multiple subjects and model the different ways in which variability can arise. Similary, the ability to capture cross-subject amplitude effects (Figure 11) or use the model to define alternatives to, for example, fALFF-type measures (Figure 13) means that many of what would have been post-hoc analyses can be simplified and made more interpretable.

### 4.1 Group-versus subject-level approaches

The comparisons in this paper have been with ICA-DR, as this is probably the most common method for finding functional modes from resting-state data and is a key part of the HCP’s pipelines. However, while PROFUMO and ICA-DR try and infer on many of the same quantities, they make fundamentally—and not necessarily compatible—assumptions about the data itself.

The key difference between the two is the way PROFUMO entails a holistic model for group- and subject-level representations, whereas ICA-DR assumes they are separately estimable. The majority of group-level ICA methods assume all subjects are in a common space, and proceed to analyse the data without recourse to individual decompositions. This formulation gives much more flexibility for the group-level decomposition to utilise the extra statistical power that concatenating over subjects affords, which means that the ICA modes depart—at times fairly radically—away from what we can resolve at the subject level. As such, ICA seems to be able to identify up to several hundred plausible components, that ultimately begin to resemble a parcellation [Kiviniemi et al. 2009; Smith et al. 2013b].

However, what we show here is that group-level representations are not enough. In the simulated data, even if the ground truth is known at the group-level, the subject-level information inferred by dual regression will be biased and noisy. What PROFUMO attempts to do is to model as many different facets of multi-subject rfMRI data together as is plausible. Here, we expand on two concrete implications of this approach as compared to other methods.

Firstly, the implication of the joint subject-level modelling in PROFUMO is that for a mode to appear at the group-level it has to be resolvable in the majority of individual subjects. Therefore, this engenders a fundamentally different view on what the dimensionality of the data is. The Bayesian model complexity penalties seem to result in no more than thirty or forty PFMs being identified, essentially regardless of the pre-specified model dimensionality. While more subjects do offer increased regularisation of the subject-level modes, this can only do so much. This is why the inferred number of PFMs is on the same order as the number of signal components as inferred by ICA-FIX (23.3 ± 6.6 at the run-level for HCP data [Marcus et al. 2013]).

Secondly, we have demonstrated the importance of modelling different characteristics of the data together. In the simulated data, even if the ground truth is known and thresholded dual regression is used to reduce the inherent spatiotemporal biases [Bijsterbosch et al. 2019], PROFUMO is still more accurate than ICA-DR like approaches. Similarly, in the classification of the active-state data, there are clear performance benefits from modelling the netmats hierarchically, even after the spatial variability has been accounted for. This is not to say that the PROFUMO model is perfect, as it clearly contains many simplifying assumptions. However, it is at least an internally consistent framework within which one can begin to explore the implications of different modelling decisions.

### 4.2 Spatial representations

One of the key messages from this work, in line with other recent reports [Hacker et al. 2013; Harrison et al. 2015; Laumann et al. 2015; Glasser et al. 2016a; Gordon et al. 2016; Braga and Buckner 2017; Gordon et al. 2017a; b; Kong et al. 2018], is that complex rearrangements of functional regions in individual subjects are ubiquitous and of a surprisingly large spatial scale. Figures 3 and 9 provide reasonable examples of these effects. Even after the advanced multi-modal, surface-based registration employed by the HCP, one often observes spatial rearrangements where subject-specific features are shifted relative to the group by many millimetres.

The difficulty we face when working at the group-level is that the summary features we extract are not necessarily representative of those at the subject-level; they are, and should always be thought of as, probabilistic representations [Van Essen and Dierker 2007]. As discussed in the previous section, we cannot automatically expect that it will be straightforward to project group-level results back to meaningful characterisations of functional connectivity at the subject level. Furthermore, the characteristic size of misalignments probably represents a limit in terms of the size of functional features we can project from the group back to the subject-level; while the native resolution of the subject-level data may well be higher, methods that work on the functional data alone like ICA-DR or PROFUMO will always struggle in the absence of additional constraints if the misalignments are large enough to mean some regions do not overlap with their group-level homologues at all.

In other words, misalignments are now often larger in scale than the fundamental resolution limits imposed by the physics and physiology that governs the properties of the data itself. Subject-level representations are limited by the properties of the data itself: 2 mm isotropic voxels are now common, and the spatial characteristics of the HRF do not appear to blur much beyond this [Shmuel et al. 2007]; at the group-level, the effective resolution of the data relates to the characteristic size of these residual misalignments between subjects, and these are likely to be larger. What this means is that functional MRI currently occupies an interesting liminal space, where the spatial resolution of high-powered single-subject analyses can now surpass that of studies that employ multitudes of subjects. This probably explains the recent resurgence of exploratory studies based on small numbers of subjects [Gonzalez-Castillo et al. 2012; Raemaekers et al. 2014; Laumann et al. 2015; Poldrack et al. 2015; Huth et al. 2016; Braga and Buckner 2017; Gordon et al. 2017b; Salehi et al. 2020]. Fortunately, recent work has suggested that there is scope to further reduce the size of the residual misalignments [Guntupalli et al. 2018], and use multi-modal data to help identify regions at the subject-level [Glasser et al. 2016a], both of which will be essential parts of the push towards finer spatial scales.

Finally, these observed spatial differences also have implications for parcel-based analyses. Given the many fine-scale variations in the spatial maps and the amount of overlap between PFMs, it may be that we need multivariate analysis techniques that go beyond one summary time course per parcel to capture the richness of the functional data at sub-parcel spatial scales [Geerligs et al. 2016; Anzellotti and Coutanche 2018; Haak et al. 2018].

### 4.3 Interpreting spatiotemporal connectivity patterns

One of the striking differences between PROFUMO and ICA-DR is their inferred patterns of functional coupling between regions. Not only do these suggest fundamentally different group-level coupling strengths, but the predictive power at the subject and run level is also different. Whereas ICA-DR netmats primarily correlate with trait-like properties, PROFUMO netmats are more sensitive to changes in cognitive state. Here, we expand on these observations as a final discussion point.

Clearly, there is a complicated relationship linking spatial variability and the functional coupling between modes, and indeed concerns about the interpretability of functional connectivity in the presence of anatomical variability are far from new [Brett et al. 2002]. The effect that subject variability in spatial organisation might have on its temporal counterpart has been noted in simulation studies. For example, E. A. Allen et al. [2012] observed a sharp decrease in the ability of a variant of ICA-DR to detect subject-specific modes in the presence of subject variability in spatial organisation, an effect which was compounded by spatial overlap between modes^13^. This links to functional coupling via the work of Smith et al. [2011], who noted that if ROIs were misspecified such that the time courses contained a range of contributions from the true underlying regions, then ‘[t]he results are extremely bad’. It is the latter result in particular which is particularly shocking: if we do not extract accurate subject-level estimates of functional regions then it is essentially impossible to characterise the functional coupling between them.

Furthermore, a key claim of the related recent work on subject variability in functional connectivity by Bijsterbosch et al. [2018] is that it is not possible to make meaningful claims about what drives cross-subject changes in functional coupling between regions if said regions are not properly delineated at the subject level. In other words, spatial variabiality does not simply make it harder to estimate functional coupling, it can also fundamentally bias our inferences. Again, the results here—particularly the simulations—extend these results, showing that the way in which dual regression biases functional connectivity estimates away from the spatial correlation structure [Bijsterbosch et al. 2019] is really an inherent property of mapping between group and subject levels in this way. While this bias can be reduced with the thresholded variant of dual regression, the simulation results, and short theoretical analysis on the role of noise, suggest that the PFM model will be much more performant than this variant.

What we show here with regards to the predictive power of the PROFUMO netmats is that, in line with other work [Bijsterbosch et al. 2018; 2019; Pervaiz et al. 2020], they are relatively poor predictors of trait-like quantities. Instead, we have shown that they are much more predictive of current cognitive state. However, for analyses that try to use functional coupling to make predictions about individual subjects [Abraham et al. 2017; Dadi et al. 2019; Pervaiz et al. 2020], the ICA-DR netmats are likely to produce more accurate predictions. In that case, one has to contend with the fact that the induced biases reduce the interpretability of the findings, which may or may not be desirable depending on the specifics of the problem at hand [Stephan et al. 2015; 2017]. Of course, the presence of confounds that are themselves behaviourally relevant—such as head motion [Power et al. 2012; Satterthwaite et al. 2012; Van Dijk et al. 2012; Couvy-Duchesne et al. 2014; Hodgson et al. 2017; Laumann et al. 2017], physiological noise [Power et al. 2017; Glasser et al. 2018] or brain volume [Bartley et al. 1997; McDaniel 2005; Qing and Gong 2016]—makes this problem of interpretability very challenging in practice for any method.

The results we have presented here suggest that the spatial information encoded by PROFUMO is likely to give much better predictive performance in this context. This is similar to other work which has demonstrated increased performance of spatial features such as, for example, task-based maps [Bijsterbosch et al. 2018] or parcel topography [Kong et al. 2018], and, furthermore, that this has a close relationship with structural information [Llera et al. 2019]. The obvious questions are therefore why do spatial rearrangements of functional regions seem to be so predictive in cross-subject analyses, and how do we interpret them? One hypothesis is that this variability in spatial organisation of functional regions is simply reflecting variability in the brain’s macroscale structure, for which there are already well established links between environmental, genetic and lifestyle factors [Reiss et al. 1996; Shaw et al. 2006; Stein et al. 2012; Douaud et al. 2014; K. G. Noble et al. 2015; Elliott et al. 2018].

However, it would be an enormous surprise if this reductionist reading of these functional changes as simply reflecting structural variability is the whole story, especially after the registration approaches used. Rather, it is vitally important to understand both what mechanisms give rise to these spatial changes, and, in particular, what unique information does the functional variability carry *over and above* what can be derived from other techniques and modalities.

## 5 Conclusions

All analyses of complex, multivariate functional data require us to make simplifying assumptions, and, as such, the results we see are inevitably coloured by the modelling choices we make. This might involve, for example, deciding decide whether to run a parcel- or mode-based analysis, or when choosing which specific method to use. As such, it is essentially impossible to conclusively determine whether one method more accurately characterises the general organisational principles or subject variability from the functional data alone. However, we feel that the above results demonstrate that PROFUMO and the PFMs model are providing a novel and worthwhile perspective on the analysis and interpretation of functional MRI data. We hope that this approach—by virtue of having a model tailored to the properties of fMRI data, the enhanced spatial sensitivity and specificity, and the way spatial variability is automatically accounted for when estimating functional coupling—proves useful.

## Acknowledgements

We would like to offer our profound thanks to Tamar Makin for making the active-state data set available, and to Roser Sala-Llonch and Sasidar Madugular for the preprocessing.

We would also like to thank the anonymous reviewers, whose feedback and suggestions led to many improvements to the manuscript.

Data were provided [in part] by the Human Connectome Project, WU-Minn Consortium (Principal Investigators: David Van Essen and Kamil Ugurbil; 1U54MH091657) funded by the 16 NIH Institutes and Centers that support the NIH Blueprint for Neuroscience Research; and by the McDonnell Center for Systems Neuroscience at Washington University.

S.J.H., J.D.B. & S.M.S. were funded by the Wellcome Trust (grants 098369/Z/12/Z and 091509/Z/10/Z). S.J.H. was also supported by the grant #2017-403 of the Strategic Focal Area “Personalized Health and Related Technologies (PHRT)” of the ETH Domain. S.M.S. also received funding from an MRC Mental Health Pathfinder grant (MC_PC_17215). A.R.S. received funding for this work from the following sources: National Institute for Health Research Oxford Biomedical Research Centre, Medical Research Council of Great Britain and Northern Ireland, the Wellcome Trust (London, UK) and the Innovative Medicines Initiative Joint Undertaking (Brussels, Belgium), under grant agreement no 115007 resources of which are composed of financial contribution from the European Union’s Seventh Framework Programme (FP7/2007–2013) and EFPIA companies’ in kind contribution. A.R.S. is also a member of the Wellcome Pain Consortium (Ref. 102645), and the project was supported by a strategic award from the Wellcome (Ref. 102645). S.P.F. is supported by the European Research Council under the European Union’s Seventh Framework Programme (Developing Human Connectome Project: FP/2007–2013/ERC Grant Agreement no. 319456). E.P.D. has been supported by the Developing Human Connectome Project (European Research Council Synergy grant FP/2007-2013), and the SSNAP charity, Oxford (Support for the Sick Newborn and their Parents). M.W.W. is funded by the Wellcome Trust (092753), and is supported by the National Institute for Health Research (NIHR) Oxford Biomedical Research Centre based at Oxford University Hospitals Trust, Oxford University (the views expressed are those of the author(s) and not necessarily those of the NHS, the NIHR or the Department of Health). Finally, the Wellcome Centre for Integrative Neuroimaging is supported by core funding from the Wellcome Trust (203139/Z/16/Z).

## Appendices

### A Alternative approaches

There are now several methods that characterise resting brain activity in terms of functional modes, both at the group and subject level. The standard pipeline is essentially a two-step process, where the group-level modes are estimated before some form of back-projection is used to extract subject-specific versions of these. Dual regression and related variants thereof, typically combined or integrated with a group-level spatial independent component analysis (ICA) [Calhoun et al. 2001; Beckmann et al. 2005], have been the de facto standard for analyses of the subject variability in spatial and, more recently, temporal features of modes for at least the past decade. Dual regression proceeds by regressing the group-level spatial maps into the data to get a set of time courses—from which subject variability in temporal features may be estimated via any number of functional connectivity metrics— before regressing the time courses back into the data to get subject-specific spatial maps [Calhoun et al. 2001; Beckmann et al. 2009; Zuo et al. 2010b; Erhardt et al. 2011; Nickerson et al. 2017].

This approach has been extended over the years, with several proposed refinements to either the method for identifying group-level modes [Damoiseaux et al. 2006; Varoquaux et al. 2010; Lee et al. 2011; Smith et al. 2012; G. I. Allen et al. 2014; Hjelm et al. 2014; Karahanoğlu and Van De Ville 2015; Dohmatob et al. 2016], or to the way subject-specific information is extracted [Du and Fan 2013; Hacker et al. 2013; Zöller et al. 2019].

However, there have been several more extensive departures from the above framework that are more similar in spirit to the hierarchical PFMs model. For example, Varoquaux et al. [2011] and Abraham et al. [2013] proposed a more holistic model that finds a set of systems regularised by not only the group-level properties, but also by the consistency of both spatial and temporal information at the subject level. More recently, Li et al. [2017] introduced a model based on non-negative matrix factorisation (NMF) that jointly optimises subject-specific decompositions such that the spatial maps are both sparse and consistent over subjects, though without explicitly leveraging any information about temporal consistency.

As mentioned in the Introduction, these methods all have potential shortcomings in terms of the extent to which typical patterns of variability are learnt from the multiple subject-specific decompositions. These shortcomings are particularly apparent for dual regression type approaches, where the estimation of subject variability is completely post-hoc (and, moreover, the estimated subject variability in spatial organisation only indirectly informs the subject variability in temporal features), but it is also problematic for the more complex models which we have mentioned, for which no explicit parameterisation for the observed variability over subjects is inferred.

More recent methodological work has focused on deriving subject-specific parcellations, both based on a fixed group-level template [Dhillon et al. 2014; Wang et al. 2015; Glasser et al. 2016a; Chong et al. 2017; Gordon et al. 2017a; Salehi et al. 2017], and formulated as a hierarchical model [Liu et al. 2012; Langs et al. 2016; Kong et al. 2018]. However, while both mode- and parcel-based approaches have shown promise, our concern is that the subject variability in spatial organisation that has been reported often features reorganisations of a similar scale to our current best estimates of the sizes of distinct functional regions [Van Essen and Dierker 2007; Van Essen et al. 2012a], and as such, reliable identification at this scale is arguably beyond all but the most sophisticated, multimodal approaches utilising high quality data [Glasser et al. 2016a]. Therefore, in this work, we stick to a system-level description and base our method on a decomposition into a set of modes. Intuitively, a functional system is more protected from the deleterious effects of misalignment than a functional region in two key ways: firstly, functional systems have a greater spatial extent than parcels; secondly, a reorganisation of one region within a larger system can be straightforwardly corrected for if the other regions are relatively stable.

### B Preprocessing

The aim of the preprocessing pipeline is to normalise the data such that it has a consistent scale across subjects, and that the properties of the unstructured noise follow the assumptions that are contained in the generative model. The approach is as follows.

- *Voxelwise normalisation.* For each voxel independently, the time course (i.e.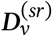) is set to zero mean and unit variance. This ensures that each voxel has a roughly equal contribution to the SVD in the next step.
- *Voxelwise normalisation of the noise subspace.* Each voxel is independently normalised such that the variance of the unstructured noise is unity. This matches the assumption of isotropic noise in the generative model. The unstructured noise subspace is estimated via the SVD. The whole data matrix is decomposed and the *M* components with the highest singular values are assumed to represent the structured signal subspace and are removed. The noise subspace is reconstructed from the remaining components, the variance is calculated in each voxel, and the data is renormalised on a voxelwise basis such that the variance becomes unity.
- *Global normalisation of the signal subspace.* There is one final degree of freedom remaining. The generative model assumes isotropic noise, but does not assume a fixed variance. Therefore we can apply a global renormalisation to set the overall variance of the modes we observe. As an approximation, if ***D*** = ***PHA*** and we assume independence over modes, then we can say that 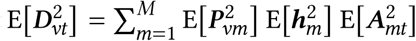. In other words, if the maps, amplitudes and time courses have unit variance then the signal variance will be equal to *M*. Therefore we use another SVD decomposition and set the overall variance of the assumed signal subspace (i.e. the first *M* components) to match the above by applying exactly the same normalisation to each voxel.

### C Data reduction

The scale of modern rfMRI studies is now such that even manipulating all the data in its raw form simultaneously is impossible. For approaches that start by inferring group-level descriptions of the data, such as ICA, it is possible to use online algorithms that work by passing over the data sequentially [Smith et al. 2014; Mensch et al. 2017], thereby removing the dependence between memory required and the number of subjects under study. However, our approach is explicitly designed to simultaneously extract group- and subject-level features, and as such we need the data from each subject to be available.

To facilitate analyses of large data-sets, we apply subject specific data reductions, but do not collapse these down further to group-level summaries. The approach we take is to approximate each run with a low-rank singular value decomposition (SVD). As our model is defined in terms of both spatial and temporal features, we have to retain both the spatial and temporal singular vectors. However, as the PFM model assumes that subject-specific spatial maps are conserved across all runs for a given subject, we make further savings by only maintaining a single set of spatial singular vectors per subject.

To do this, we calculate the SVD of the matrix formed by temporally concat-enating all data from a given subject. This combined data matrix,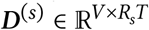,is then represented by ***U***^(*s*)^, ***S***^(*s*)^ and ***V***^(*s*)^. To approximate this with a low rank SVD, we simply only retain the singular vectors associated with the top *N* singular values. For example, assuming *V* > *R*_*s*_*T* and ignoring columns associated with singular values equal to zero, 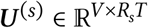 is replaced by ***Û***^(*s*)^ ∈ ℝ ^*V* ×*N*^. Finally, we can partition the temporal singular vectors, according to the order the individual runs were concatenated, in order to reconstruct the data from each run individually, or in other words, 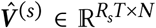is decomposed into a set of 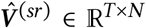. In summary, each data matrix, ***D***_(*sr*)_, hasthree approximating matrices, namely ***Û***^(*s*)^ ∈ ℝ^*V*×*N*^, ***Ŝ***^(*s*)^ ∈ ℝ^*N*×*N*^ and 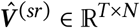.

The last thing we do is to combine these three matrices into two matrices. This simply saves some computation each time we need to calculate any expectations involving the data. The final form for the approximate data is therefore

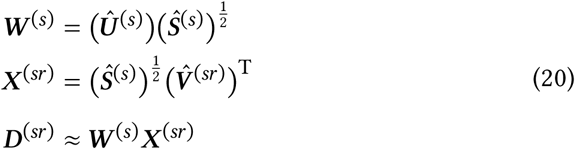

We can simply substitute this approximate expression for ***D***_(*sr*)_ any time we need access to the data in the inference procedure, and this has the added bonus of being computationally, as well as space, efficient. However, we explicitly calculate, and cache, the overall data variance from the full data, rather than ignoring the contribution from the subspace of discarded singular values^14^. This means that the estimate for the noise precision, *ψ* _(*sr*)_, will be comparable whether or not we choose to utilise this low-rank approximation, or indeed across different values of *N.*

We now have an explicit method for reducing large data-sets to a more manageable size. However, there is one final complication: computationally, calculating the SVD of every ***D***^(*s*)^ actually turns out to be prohibitively expensive in most cases. In order to circumvent this, we utilise the fact that we are explicitly looking for a low-rank approximation and implement an extremely efficient randomised algorithm to directly calculate the truncated SVD. This approach is described in the excellent review by Halko et al. [2011].

### D Degrees of freedom correction

fMRI data has an inherent spatial smoothness—such that there are non-trivial spatial autocorrelations in the noise processes—which is amplified by the spatial smoothing that is a standard pre-processing step for most analyses. As discussed earlier, this is not acknowledged in our specification of our model of the noise process. In essence, this means that the model assumes that there are more independent spatial measurements than actually exist.

Fortunately, as Groves et al. [2011] discuss, there is a simple way to mitigate some of the effects of this within the Bayesian framework. Intuitively, if we have smoothed the data then we should be able to downsample it without loss of information. At some stage, this would result in the noise becoming genuinely spatially independent again, thereby satisfying the assumptions of the generative model. However, this presents several practical problems, so rather than actually downsample the data, we simply downweight the spatial information by a factor *v*. This represents the proportion of voxels that would be retained if we were to optimally downsample. ‘This is analogous to fixing that only a random fraction of the data points will be kept, but at each stage averaging over all possible choices of decimated voxels’ [ibid.].

While this approach still does not explicitly acknowledge the relationship between noise in nearby voxels, it does counter most of the deleterious effects of this model misspecification, especially when combined with the models for noise in the subject-specific spatial maps and time courses. The main advantage of this approach, compared to a more formal model for smoothness, is that it remains particularly computationally efficient.

### E Initialisation

With a model of this complexity, it is important that the algorithm is appropriately initialised. By doing so, we can improve reliability and computational stability whilst reducing the computational time required for convergence. Our approach is to compute a consensus group-level set of modes, and use these to initialise the full model.

To do this, we mimic the temporal concatenation approach employed by most existing algorithms and compute a consensus set of spatial singular vectors (this can be done even more efficiently if we have already utilised the data reduction technique described previously [Calhoun et al. 2001]) using another randomised SVD algorithm that streams over the data. These singular vectors are reweighted via an adjusted set of singular values. More specifically, we use the properties of the Marchenko-Pastur distribution to find the noise level that ensures the SNR at the group-level decomposition is similar to the SNR at the subject level. We then run a Bayesian version of spatial ICA—with the spatial priors set to mimic the group-level priors of the full model—to generate the group-level modes. The SNR recalibration ensures we do not get over-splitting of the modes at this stage. We can then propagate this set of group-level modes through the rest of the algorithm, thereby ensuring all parameters are initialised with plausible values.

### F Simulations

The spatial model consists of two levels: parcels and modes. We simulate 100 spatially contiguous parcels within a one-dimensional space comprising 10,000 voxels. We then apply a random diffeomorphic warp to each subject separately as a model for residual misalignments after registration. We then simulate a set of 15 modes consisting of blocks of spatially adjacent parcels. There is variability in the mode weights over subjects, and we introduce overlap such that, on average, each voxel is a member of 1.4 modes.

In the temporal domain we simulate a set of sparse, correlated ‘neural’ time-courses for each of the two runs per subject. There is variability in the between-mode correlation structure at the run and subject level. These are then convolved with a random draw from the FLOBS basis of hæmodynamic response functions [Woolrich et al. 2004], which introduces variability over subjects and space. This results in 500 timepoints at a TR of 2.0 s.

The spatial maps and timecourses are combined via the outer product model, and a nonlinear saturation is applied such that the highest amplitude moments of instantaneous voxelwise activity are reduced. Finally, random noise is added with a degree of spatiotemporal smoothness such that the overall SNR (expressed in terms of variance) is 0.1. For the simulations presented in the Supplementary Material, we also add some structured, subject-specific noise components, again using an outer-product model. These can either be spatially specific or global, and are designed to contribute a similar amount of variance per component as the individual signal modes.

To allow inference algorithms to model the aforementioned artefacts, 18 modes are inferred. After inference, modes are paired to the ground truth based on the similarity between both the spatial maps and timecourses, averaged over runs and subjects. The full set of performance metrics shown in the Supplementary Material is then calculated.

### G Human Connectome Project data and analyses

For the HCP analyses, all data was from the 1,200 Subjects Data Release: humanconnectome. org/study/hcp-young-adult/document/1200-subjects-data-release. We used the 1,003 subjects for whom there was full behavioural, structural and rfMRI data (i.e. 4 runs, each of 1,200 volumes). All analyses are of the MSMAll and FIX cleaned data (i.e. `rfMRI_REST1_LR_Atlas_MSMAll_hp2000_clean.dtseries.nii` etc.).

ICA-DR results were taken from the Extensively Processed fMRI Data: humanconnectome.org/study/hcp-young-adult/document/extensively-processed-fmri-data-documentation. The amplitudes and netmats were estimated from the time courses released by the HCP: the amplitudes were taken as the standard deviations, while the netmats were the partial correlation matrices. Tikhonov regularisation was used when calculating the inverse of the full correlation matrices, with ***Γ*** = 0.1***I***.

Heritability was estimated via Falconer’s formula, 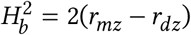 [Falconer 1960]. We calculate the correlations, *r*_*mz*_ and *r*_*dz*_, between the voxelwise spatial map weights. In other words, for each subject and each voxel we extract a length *M* vector of weights: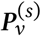 using the PFM notation, and compute the correlation between these for every pair of subjects.

For the CCA analyses, we used the full set of restricted information released by the HCP. We first removed all variables relating to study completion or quality control. The structural variables were all the remaining variables in the Free-Surfer category; all others were taken as behavioural. To preprocess the behavioural variables, we first removed any variables that were either more than 20 % NaN, or those for which more than 95 % of subjects had exactly the same entries. We then imputed any missing values using the SoftImpute method [Mazumder et al. 2010] as implemented in the `fancyimpute` Python package (github.com/ iskandr/fancyimpute). The following were regressed out as confounds (one-hot encoded where necessary) in all subsequent analyses: `Release`, `Acquisition`, `fMRI_3T_ReconVrs`, `rfMRI_motion`, `Age`, `Gender`, `Race`, `Ethnicity`, `Handedness`, `Height`, `Weight`, `BMI`, `BPSystolic`, `BPDiastolic`, `Hematocrit_1`, `Hematocrit_2`, `FS_IntraCranial_Vol`, `FS_BrainSeg_Vol`. A detailed description of all variables can be found at wiki.humanconnectome.org/display/PublicData/HCP+Data+Dictionary+Public-+Updated+for+the+1200+Subject+Release.

For the CCA, all groups of variables were normalised and then reduced to their top 25 components via the SVD, before a CCA was run on every pair of variable groups. The RV coefficient was then calculated between the top 10 paired components from each CCA.

### H Active-state data and analyses

Data was acquired from fifteen subjects, but for these analyses we excluded Subject `07` due to potential artefacts in several of their scans. Preprocessing was as previously published (i.e. brain extraction, B0 unwarping, high-pass temporal filtering, motion correction, and FIX cleaning) [Kieliba et al. 2019]. However, we did not apply mean-based intensity normalisation or low-pass filter the data. Finally, the pre-processed functional scans were then registered to MNI space and spatially smoothed (2 mm FWHM).

As with the HCP data, the ICA-DR amplitudes and netmats were estimated from the time courses: the amplitudes were taken as the standard deviations, while the netmats were the partial correlation matrices. Tikhonov regularisation was used when calculating the inverse of the full correlation matrices, with ***Γ*** = 0.1***I***. The SVM was from scikit-learn (`sklearn.svm.SVC`), and as this is a relatively small dataset parameters were left at their defaults [Varoquaux et al. 2017].

## Footnotes

1 For a more complete overview of data sharing initiatives, see the NeuroImage special issues [Eickhoff et al. 2016].

2 For the rest of this paper, when we use the term variability in relation to functional measures, it can be assumed to relate to variability in static functional connectivity over subjects or sessions. This does not consider, for example, the moment-to-moment fluctuations characterised as dynamic functional connectivity [Hutchison et al. 2013; Calhoun et al. 2014; Preti and Van De Ville 2017]. This encodes important within-subject state changes [Tagliazucchi and Laufs 2014], and there is growing evidence that this captures between-subject trait differences too [Vidaurre et al. 2017].

3 Resting-state networks, intrinsic connectivity networks, etc.

4 We make this distinction as the spatial maps, which characterise the location of functional regions, also capture aspects functional connectivity and organisation.

5 It is somewhat contentious whether (structural) registration should be held responsible for the latter two processes. Our definition of registration is somewhat broader, as we hold it responsible for bringing subjects into structural and functional correspondence. While structural registration is unlikely to be sufficient here, this is nevertheless a reasonable aim for multi-modal registration approaches. For a good discussion of these issues see e.g. Van Essen and Dierker [2007].

6 Eyes-open, eyes-closed, pre/post an intervention, various ‘active-state’ paradigms etc.

7 Cf. the noisy estimates of beta values from a GLM fit.

8 See Colclough et al. [2018] for a discussion of exactly this effect in relation to inference of functional couplings between regions.

9 See e.g. Gelman [2006] for a related discussion of redundant parameterisations of variance.

10 For adult populations, both the SPM double-gamma HRF [Friston et al. 2007] or the principal component of the FLOBS basis in FSL [Woolrich et al. 2004] are provided, though this can be replaced for different populations as appropriate e.g. [Arichi et al. 2012].

11 db.humanconnectome.org/megatrawl/index.html

12 Note also that Figure 2 from Sala-Llonch et al. [2019] uses FDR with *q* = 0.2 for the background,whereas the tests here use a more stringent FWE *p* < 0.05 test for significance.

13 Figures 4 and 5 in particular.

14 More explicitly, we use Tr((***D***^(*sr*)^)^T^***D***^(*sr*)^) rather than Tr((***W***^(*s*)^***X*** ^(*sr*)^)^T^***W*** ^(*s*)^***X*** ^(*sr*)^) whenever required.

## References

Abraham, A., Dohmatob, E., Thirion, B., Samaras, D. and Varoquaux, G. (2013). Extracting Brain Regions from Rest fMRI with Total-Variation Constrained Dictionary Learning. In: Medical Image Computing and Computer-Assisted Intervention— MICCAI 2013. Ed. by K. Mori, I. Sakuma, Y. Sato, C. Barillot and N. Navab. Vol. 8150. Lecture Notes in Computer Science. Springer Berlin Heidelberg, pp. 607–615.

Abraham, A., Milham, M. P., Di Martino, A., Craddock, R. C., Samaras, D. et al. (2017). Deriving reproducible biomarkers from multi-site resting-state data: An Autism-based example. In: NeuroImage 147 (Feb. 2017), pp. 736–745.

Allen, E. A., Erhardt, E. B., Wei, Y., Eichele, T. and Calhoun, V. D. (2012). Capturing inter-subject variability with group independent component analysis of fMRI data: A simulation study. In: NeuroImage 59.4 (Feb. 2012), pp. 4141–4159.

Allen, E. A., Erhardt, E. B., Damaraju, E., Gruner, W., Segall, J. M. et al. (2011). A baseline for the multivariate comparison of resting state networks. In: Frontiers in Systems Neuroscience 5.2.

Allen, G. I., Grosenick, L. and Taylor, J. (2014). A Generalized Least-Square Matrix Decomposition. In: Journal of the American Statistical Association 109.505, pp. 145–159. eprint: http://www.tandfonline.com/doi/pdf/10.1080/01621459.2013.852978.

Amiez, C. and Petrides, M. (2014). Neuroimaging Evidence of the Anatomo-Functional Organization of the Human Cingulate Motor Areas. In: Cerebral Cortex 24.3 (Mar. 2014), pp. 563–578. eprint: http://cercor.oxfordjournals.org/content/24/3/563.full.pdf+html.

Amunts, K., Schleicher, A. and Zilles, K. (2007). Cytoarchitecture of the cerebral cortex—More than localization. In: NeuroImage 37.4 (Oct. 2007), pp. 1061–1065.

Andrews, T. J., Halpern, S. D. and Purves, D. (1997). Correlated Size Variations in Human Visual Cortex, Lateral Geniculate Nucleus, and Optic Tract. In: The Journal of Neuroscience 17.8 (Apr. 1997), pp. 2859–2868. eprint: http://www.jneurosci.org/content/17/8/2859.full.pdf+html.

Anzellotti, S. and Coutanche, M. N. (2018). Beyond Functional Connectivity: Investigating Networks of Multivariate Representations. In: Trends in Cognitive Sciences 22.3 (Mar. 2018), pp. 258–269.

Arichi, T., Fagiolo, G., Varela, M., Melendez-Calderon, A., Allievi, A. et al. (2012). Development of BOLD signal hemodynamic responses in the human brain. In: NeuroImage 63.2 (Nov. 2012), pp. 663–673.

Attias, H. (2000). A Variational Bayesian Framework for Graphical Models. In: Advances in Neural Information Processing Systems 12. Ed. by S. A. Solla, T. K. Leen and K. Müller. MIT Press, pp. 209–215.

Bamberg, F., Kauczor, H.-U., Weckbach, S., Schlett, C. L., Forsting, M. et al. (2015). Whole-Body MR Imaging in the German National Cohort: Rationale, Design, and Technical Background. In: Radiology 277.1 (Oct. 2015), pp. 206–220. eprint: https://doi.org/10.1148/radiol.2015142272.

Bartley, A. J., Jones, D. W. and Weinberger, D. R. (1997). Genetic variability of human brain size and cortical gyral patterns. In: Brain 120.2, pp. 257–269. eprint: http://brain.oxfordjournals.org/content/120/2/257.full.pdf+html.

Beckmann, C. F., DeLuca, M., Devlin, J. T. and Smith, S. M. (2005). Investigations into resting-state connectivity using independent component analysis. In: Philosophical Transactions of the Royal Society B: Biological Sciences 360.1457, pp. 1001–1013. eprint: http://rstb.royalsocietypublishing.org/content/360/1457/1001.full.pdf+html.

Beckmann, C. F., Mackay, C., Filippini, N. and Smith, S. M. (2009). Group comparison of resting-state FMRI data using multi-subject ICA and dual regression. In: Organization for Human Brain Mapping.

Beckmann, C. F. and Smith, S. M. (2004). Probabilistic independent component analysis for functional magnetic resonance imaging. In: Medical Imaging, IEEE Transactions on 23.2 (Feb. 2004), pp. 137–152.

Bijsterbosch, J. D., Beckmann, C. F., Woolrich, M. W., Smith, S. M. and Harrison, S. J. (2019). The relationship between spatial configuration and functional connectivity of brain regions revisited. In: eLife 8 (May 2019), e44890.

Bijsterbosch, J. D., Harrison, S. J., Duff, E. P., Alfaro-Almagro, F., Woolrich, M. W. et al. (2017). Investigations into within- and between-subject resting-state amplitude variations. In: NeuroImage 159 (Oct. 2017), pp. 57–69.

Bijsterbosch, J. D., Woolrich, M. W., Glasser, M. F., Robinson, E. C., Beckmann, C. F. et al. (2018). The relationship between spatial configuration and functional connectivity of brain regions. In: eLife 7 (Feb. 2018). Ed. by C. Honey, e32992.

Biswal, B. B., Mennes, M., Zuo, X.-N., Gohel, S., Kelly, C. et al. (2010). Toward discovery science of human brain function. In: Proceedings of the National Academy of Sciences 107.10 (Mar. 2010), pp. 4734–4739. eprint: http://www.pnas.org/content/107/10/4734.full.pdf.

Blei, D. M., Kucukelbir, A. and McAuliffe, J. D. (2017). Variational Inference: A Review for Statisticians. In: Journal of the American Statistical Association 112.518, pp. 859–877. eprint: http://dx.doi.org/10.1080/01621459.2017.1285773.

Braga, R. M. and Buckner, R. L. (2017). Parallel Interdigitated Distributed Networks within the Individual Estimated by Intrinsic Functional Connectivity. In: Neuron 95.2 (July 2017), 457–471.e5.

Breteler, M. M. B., Stöcker, T., Pracht, E., Brenner, D. and Stirnberg, R. (2014). MRI in the Rhineland Study: A Novel Protocol for Population Neuroimaging. In: Alzheimer’s & Dementia 10.4, p. 92.

Brett, M., Johnsrude, I. S. and Owen, A. M. (2002). The problem of functional localization in the human brain. In: Nature Reviews Neuroscience 3.3 (Mar. 2002), pp. 243–249.

Bright, M. G. and Murphy, K. (2015). Is fMRI “noise” really noise? Resting state nuisance regressors remove variance with network structure. In: NeuroImage 114 (July 2015), pp. 158–169.

Buckner, R. L., Andrews-Hanna, J. R. and Schacter, D. L. (2008). The Brain’s Default Network: Anatomy, Function, and Relevance to Disease. In: Annals of the New York Academy of Sciences 1124.1, pp. 1–38.

Buckner, R. L., Krienen, F. M. and Yeo, B. T. T. (2013). Opportunities and limitations of intrinsic functional connectivity MRI. In: Nature Neuroscience 16.7 (July 2013), pp. 832–837.

Calhoun, V. D., Adali, T., Pearlson, G. D. and Pekar, J. J. (2001). A method for making group inferences from functional MRI data using independent component analysis. In: Human Brain Mapping 14.3 (Nov. 2001), pp. 140–151.

Calhoun, V. D., Miller, R., Pearlson, G. and Adali, T. (2014). The Chronnectome: Time-Varying Connectivity Networks as the Next Frontier in fMRI Data Discovery. In: Neuron 84.2 (Oct. 2014), pp. 262–274.

Calhoun, V. D., Potluru, V. K., Phlypo, R., Silva, R. F., Pearlmutter, B. A. et al. (2013). Independent Component Analysis for Brain fMRI Does Indeed Select for Maximal Independence. In: PLoS ONE 8.8 (Aug. 2013), e73309.

Chong, M., Bhushan, C., Joshi, A. A., Choi, S., Haldar, J. P. et al. (2017). Individual parcellation of resting fMRI with a group functional connectivity prior. In: NeuroImage 156, pp. 87–100.

Coalson, T. S., Van Essen, D. C. and Glasser, M. F. (2018). The impact of traditional neuroimaging methods on the spatial localization of cortical areas. In: Proceedings of the National Academy of Sciences 115.27, E6356–E6365.

Colclough, G. L., Smith, S. M., Nichols, T. E., Winkler, A. M., Sotiropoulos, S. N. et al. (2017). The heritability of multi-modal connectivity in human brain activity. In: eLife 6 (July 2017). Ed. by J. L. Gallant, e20178.

Colclough, G. L., Woolrich, M. W., Harrison, S. J., Rojas López, P. A., Valdes-Sosa, P. A. et al. (2018). Multi-subject hierarchical inverse covariance modelling improves estimation of functional brain networks. In: NeuroImage 178 (Sept. 2018), pp. 370–384.

Conroy, B. R., Singer, B. D., Guntupalli, J. S., Ramadge, P. J. and Haxby, J. V. (2013). Inter-subject alignment of human cortical anatomy using functional connectivity. In: NeuroImage 81 (Nov. 2013), pp. 400–411.

Couvy-Duchesne, B., Blokland, G. A. M., Hickie, I. B., Thompson, P. M., Martin, N. G. et al. (2014). Heritability of head motion during resting state functional MRI in 462 healthy twins. In: NeuroImage 102, Part 2 (Nov. 2014), pp. 424–434.

Dadi, K., Rahim, M., Abraham, A., Chyzhyk, D., Milham, M. et al. (2019). Benchmarking functional connectome-based predictive models for resting-state fMRI. In: NeuroImage 192 (May 2019), pp. 115–134.

Damoiseaux, J. S., Rombouts, S. A. R. B., Barkhof, F., Scheltens, P., Stam, C. J. et al. (2006). Consistent resting-state networks across healthy subjects. In: Proceedings of the National Academy of Sciences 103.37 (Sept. 2006), pp. 13848–13853. eprint: http://www.pnas.org/content/103/37/13848.full.pdf+html.

Devlin, J. T. and Poldrack, R. A. (2007). In praise of tedious anatomy. In: NeuroImage 37.4 (Oct. 2007), pp. 1033–1041.

Dhillon, P. S., Wolk, D. A., Das, S. R., Ungar, L. H., Gee, J. C. et al. (2014). Subject-specific functional parcellation via Prior Based Eigenanatomy. In: NeuroImage 99 (Oct. 2014), pp. 14–27.

Dohmatob, E., Mensch, A., Varoquaux, G. and Thirion, B. (2016). Learning brain regions via large-scale online structured sparse dictionary learning. In: Advances in Neural Information Processing Systems 29. Ed. by D. D. Lee, M. Sugiyama, U. V. Luxburg, I. Guyon and R. Garnett. Curran Associates, Inc., pp. 4610–4618.

Douaud, G., Groves, A. R., Tamnes, C. K., Westlye, L. T., Duff, E. P. et al. (2014). A common brain network links development, aging, and vulnerability to disease. In: Proceedings of the National Academy of Sciences 111.49 (Dec. 2014), pp. 17648–17653.

Dougherty, R. F., Koch, V. M., Brewer, A. A., Fischer, B., Modersitzki, J. et al. (2003). Visual field representations and locations of visual areas V1/2/3 in human visual cortex. In: Journal of Vision 3.10 (Oct. 2003), pp. 586–598. eprint: /data/Journals/JOV/933499/jov-3-10-1.pdf.

Du, Y. and Fan, Y. (2013). Group information guided ICA for fMRI data analysis. In: NeuroImage 69 (Apr. 2013), pp. 157–197.

Dubois, J. and Adolphs, R. (2016). Building a Science of Individual Differences from fMRI. In: Trends in Cognitive Sciences 20.6 (June 2016), pp. 425–443.

Duff, E. P., Johnston, L. A., Xiong, J., Fox, P. T., Mareels, I. et al. (2008). The power of spectral density analysis for mapping endogenous BOLD signal fluctuations. In: Human Brain Mapping 29.7 (July 2008), pp. 778–790. eprint: https://onlinelibrary.wiley.com/doi/pdf/10.1002/hbm.20601.

Duff, E. P., Makin, T., Cottaar, M., Smith, S. M. and Woolrich, M. W. (2018). Disambiguating brain functional connectivity. In: NeuroImage 173 (June 2018), pp. 540–550. eprint: https://www.biorxiv.org/content/early/2017/01/25/103002.full.pdf.

Eickhoff, S., Nichols, T. E., Van Horn, J. D. and Turner, J. A. (2016). Sharing the wealth: Neuroimaging data repositories. In: NeuroImage 124.Part B (Jan. 2016), pp. 1065–1068.

Eklund, A., Nichols, T. E. and Knutsson, H. (2016). Cluster failure: Why fMRI inferences for spatial extent have inflated false-positive rates. In: Proceedings of the National Academy of Sciences 113.28 (July 2016), pp. 7900–7905.

Elliott, L. T., Sharp, K., Alfaro-Almagro, F., Shi, S., Miller, K. L. et al. (2018). Genomewide association studies of brain imaging phenotypes in UK Biobank. In: Nature 562.7726 (Oct. 2018), pp. 210–216.

Erhardt, E. B., Rachakonda, S., Bedrick, E. J., Allen, E. A., Adali, T. et al. (2011). Comparison of multi-subject ICA methods for analysis of fMRI data. In: Human Brain Mapping 32.12, pp. 2075–2095.

Esposito, F., Scarabino, T., Hyvärinen, A., Himberg, J., Formisano, E. et al. (2005). Independent component analysis of fMRI group studies by self-organizing clustering. In: NeuroImage 25.1 (Mar. 2005), pp. 193–205.

Fagerland, M. W., Lydersen, S. and Laake, P. (2013). The McNemar test for binary matched-pairs data: mid-p and asymptotic are better than exact conditional. In: BMC Medical Research Methodology 13, p. 91.

Falconer, D. S. (1960). Introduction to Quantitative Genetics. Oliver & Boyd, Edinburgh/London.

Finn, E. S., Shen, X., Scheinost, D., Rosenberg, M. D., Huang, J. et al. (2015). Functional connectome fingerprinting: identifying individuals using patterns of brain connectivity. In: Nature Neuroscience 18.11 (Nov. 2015), pp. 1664–1671.

Friston, K. J. (2011). Functional and Effective Connectivity: A Review. In: Brain Connectivity 1.1 (June 2011), pp. 13–36.

Friston, K. J., Ashburner, J., Kiebel, S., Nichols, T. and Penny, W., eds. (2007). Statistical Parametric Mapping: The Analysis of Functional Brain Images. Academic Press.

Friston, K. J., Frith, C. D., Liddle, P. F. and Frackowiak, R. S. J. (1993). Functional Connectivity: The Principal-Component Analysis of Large (PET) Data Sets. In: Journal of Cerebral Blood Flow & Metabolism 13.1 (Jan. 1993), pp. 5–14. eprint: http://dx.doi.org/10.1038/jcbfm.1993.4.

Geerligs, L., Cam-CAN and Henson, R. N. (2016). Functional connectivity and structural covariance between regions of interest can be measured more accurately using multivariate distance correlation. In: NeuroImage 135 (July 2016), pp. 16–31.

Gelman, A. (2006). Prior distributions for variance parameters in hierarchical models (Comment on Article by Browne and Draper). In: Bayesian Analysis 1.3 (Sept. 2006), pp. 515–534.

George, E. I. and McCulloch, R. E. (1993). Variable Selection via Gibbs Sampling. In: Journal of the American Statistical Association 88.423, pp. 881–889. eprint: http://amstat.tandfonline.com/doi/pdf/10.1080/01621459.1993.10476353.

Glasser, M. F., Coalson, T. S., Bijsterbosch, J. D., Harrison, S. J., Harms, M. P. et al. (2018). Using temporal ICA to selectively remove global noise while preserving global signal in functional MRI data. In: NeuroImage 181 (Nov. 2018), pp. 692–717.

Glasser, M. F., Coalson, T. S., Robinson, E. C., Hacker, C. D., Harwell, J. et al. (2016a). A multi-modal parcellation of human cerebral cortex. In: Nature 536 (Aug. 2016), pp. 171–178.

Glasser, M. F., Smith, S. M., Marcus, D. S., Andersson, J. L. R., Auerbach, E. J. et al. (2016b). The Human Connectome Project’s neuroimaging approach. In: Nature Neuroscience 19.9 (Sept. 2016), pp. 1175–1187.

Glasser, M. F., Sotiropoulos, S. N., Wilson, J. A., Coalson, T. S., Fischl, B. et al. (2013). The minimal preprocessing pipelines for the Human Connectome Project. In: NeuroImage 80 (Oct. 2013), pp. 105–124.

Glasser, M. F. and Van Essen, D. C. (2011). Mapping Human Cortical Areas In Vivo Based on Myelin Content as Revealed by T1- and T2-Weighted MRI. In: The Journal of Neuroscience 31.32 (Aug. 2011), pp. 11597–11616. eprint: http://www.jneurosci.org/content/31/32/11597.full.pdf+html.

Gonzalez-Castillo, J., Saad, Z. S., Handwerker, D. A., Inati, S. J., Brenowitz, N. et al. (2012). Whole-brain, time-locked activation with simple tasks revealed using massive averaging and model-free analysis. In: Proceedings of the National Academy of Sciences 109.14 (Apr. 2012), pp. 5487–5492. eprint: http://www.pnas.org/content/109/14/5487.full.pdf.

Gordon, E. M., Laumann, T. O., Adeyemo, B., Gilmore, A. W., Nelson, S. M. et al. (2016). Individual-specific features of brain systems identified with resting state functional correlations. In: NeuroImage 146 (Feb. 2016), pp. 918–939.

Gordon, E. M., Laumann, T. O., Adeyemo, B. and Petersen, S. E. (2017a). Individual Variability of the System-Level Organization of the Human Brain. In: Cerebral Cortex 27.1 (Jan. 2017), pp. 386–399.

Gordon, E. M., Laumann, T. O., Gilmore, A. W., Newbold, D. J., Greene, D. J. et al. (2017b). Precision Functional Mapping of Individual Human Brains. In: Neuron 95.4 (Aug. 2017), pp. 791–807.

Gorgolewski, K. J., Esteban, O., Schaefer, G., Wandell, B. and Poldrack, R. A. (2017). OpenNeuro - a free online platform for sharing and analysis of neuroimaging data. In: Organization for Human Brain Mapping.

Gratton, C., Laumann, T. O., Nielsen, A. N., Greene, D. J., Gordon, E. M. et al. (2018). Functional Brain Networks Are Dominated by Stable Group and Individual Factors, Not Cognitive or Daily Variation. In: Neuron 98.2 (Apr. 2018), pp. 439–452.

Greicius, M. D., Krasnow, B., Reiss, A. L. and Menon, V. (2003). Functional connectivity in the resting brain: A network analysis of the default mode hypothesis. In: Proceedings of the National Academy of Sciences 100.1 (Jan. 2003), pp. 253–258. eprint: http://www.pnas.org/content/100/1/253.full.pdf+html.

Groves, A. R., Beckmann, C. F., Smith, S. M. and Woolrich, M. W. (2011). Linked independent component analysis for multimodal data fusion. In: NeuroImage 54.3, pp. 2198–2217.

Guntupalli, J. S., Feilong, M. and Haxby, J. V. (2018). A computational model of shared fine-scale structure in the human connectome. In: PLOS Computational Biology 14.4 (Apr. 2018), pp. 1–26.

Guntupalli, J. S., Hanke, M., Halchenko, Y. O., Connolly, A. C., Ramadge, P. J. et al. (2016). A Model of Representational Spaces in Human Cortex. In: Cerebral Cortex 26.6 (June 2016), pp. 2919–2934. eprint: /oup/backfile/Content_public/Journal/cercor/26/6/10.1093_cercor_bhw068/3/bhw068.pdf.

Haak, K. V., Marquand, A. F. and Beckmann, C. F. (2018). Connectopic mapping with resting-state fMRI. In: NeuroImage 170 (Apr. 2018), pp. 83–94.

Hacker, C. D., Laumann, T. O., Szrama, N. P., Baldassarre, A., Snyder, A. Z. et al. (2013). Resting state network estimation in individual subjects. In: NeuroImage 82 (Nov. 2013), pp. 616–633.

Haldane, J. B. S. (1932). A note on inverse probability. In: Mathematical Proceedings of the Cambridge Philosophical Society 28.1 (Jan. 1932), pp. 55–61.

Halko, N., Martinsson, P. G. and Tropp, J. A. (2011). Finding Structure with Randomness: Probabilistic Algorithms for Constructing Approximate Matrix Decompositions. In: SIAM Review 53.2, pp. 217–288. eprint: http://dx.doi.org/10.1137/090771806.

Harrison, S. J., Woolrich, M. W., Robinson, E. C., Glasser, M. F., Beckmann, C. F. et al. (2015). Large-scale Probabilistic Functional Modes from resting state fMRI. In: NeuroImage 109 (Apr. 2015), pp. 217–231.

Hasan, A., McIntosh, A. M., Droese, U.-A., Schneider-Axmann, T., Lawrie, S. M. et al. (2011). Prefrontal cortex gyrification index in twins: an MRI study. English. In: European Archives of Psychiatry and Clinical Neuroscience 261.7, pp. 459–465.

Hjelm, R. D., Calhoun, V. D., Salakhutdinov, R., Allen, E. A., Adali, T. et al. (2014). Restricted Boltzmann machines for neuroimaging: An application in identifying intrinsic networks. In: NeuroImage 96 (Aug. 2014), pp. 245–260.

Hodgson, K., Poldrack, R. A., Curran, J. E., Knowles, E. E., Mathias, S. et al. (2017). Shared Genetic Factors Influence Head Motion During MRI and Body Mass Index. In: Cerebral Cortex 27.12 (Dec. 2017), pp. 5539–5546.

Horien, C., Shen, X., Scheinost, D. and Constable, R. T. (2019). The individual functional connectome is unique and stable over months to years. In: NeuroImage 189, pp. 676–687.

Hotelling, H. (1936). Relations between two sets of variates. In: Biometrika 28.3-4 (Dec. 1936), pp. 321–377.

Hutchison, R. M., Womelsdorf, T., Allen, E. A., Bandettini, P. A., Calhoun, V. D. et al. (2013). Dynamic functional connectivity: Promise, issues, and interpretations. In: NeuroImage 80 (Oct. 2013), pp. 360–378.

Huth, A. G., Heer, W. A. de, Griffiths, T. L., Theunissen, F. E. and Gallant, J. L. (2016). Natural speech reveals the semantic maps that tile human cerebral cortex. In: Nature 532.7600 (Apr. 2016), pp. 453–458.

Insel, T. R. and Cuthbert, B. N. (2015). Brain disorders? Precisely. In: Science 348.6234 (May 2015), pp. 499–500. eprint: http://science.sciencemag.org/content/348/6234/499.full.pdf.

Ishwaran, H. and Rao, J. S. (2005). Spike and Slab Variable Selection: Frequentist and Bayesian Strategies. In: The Annals of Statistics 33.2, pp. 730–773.

Jenkinson, M., Beckmann, C. F., Behrens, T. E. J., Woolrich, M. W. and Smith, S. M. (2012). FSL. In: NeuroImage 62.2, pp. 782–790.

Johansen-Berg, H., Behrens, T. E. J., Robson, M. D., Drobnjak, I., Rushworth, M. F. S. et al. (2004). Changes in connectivity profiles define functionally distinct regions in human medial frontal cortex. In: Proceedings of the National Academy of Sciences 101.36 (Sept. 2004), pp. 13335–13340. eprint: http://www.pnas.org/content/101/36/13335.full.pdf.

Karahanoglu, F. I. and Van De Ville, D. (2015). Transient brain activity disentangles fMRI resting-state dynamics in terms of spatially and temporally overlapping networks. In: Nature Communications 6.7751 (July 2015).

Kennedy, D. N., Haselgrove, C., Riehl, J., Preuss, N. and Buccigrossi, R. (2016). The NITRC image repository. In: NeuroImage 124.Part B (Jan. 2016), pp. 1069–1073.

Kieliba, P., Madugula, S., Filippini, N., Duff, E. P. and Makin, T. R. (2019). Large-scale intrinsic connectivity is consistent across varying task demands. In: PLOS ONE 14.4 (Apr. 2019), pp. 1–21.

Kiviniemi, V., Starck, T., Remes, J., Long, X., Nikkinen, J. et al. (2009). Functional segmentation of the brain cortex using high model order group PICA. In: Human Brain Mapping 30.12, pp. 3865–3886.

Kong, R., Li, J., Orban, C., Sabuncu, M. R., Liu, H. et al. (2018). Spatial Topography of Individual-Specific Cortical Networks Predicts Human Cognition, Personality, and Emotion. In: Cerebral Cortex, bhy123. eprint: /oup/backfile/content_public/journal/cercor/pap/10.1093_cercor_bhy123/1/bhy123.pdf.

Kriegeskorte, N., Bodurka, J. and Bandettini, P. (2008). Artifactual time-course correlations in echo-planar fMRI with implications for studies of brain function. In: International Journal of Imaging Systems and Technology 18.5-6, pp. 345–349.

Krienen, F. M., Yeo, B. T. T. and Buckner, R. L. (2014). Reconfigurable task-dependent functional coupling modes cluster around a core functional architecture. In: Philosophical Transactions of the Royal Society of London B: Biological Sciences 369.1653 (Oct. 2014), p. 20130526. eprint: http://rstb.royalsocietypublishing.org/content/369/1653/20130526.full.pdf.

Kruschke, J. (2014). Doing Bayesian Data Analysis: A Tutorial with R, JAGS, and Stan. 2nd ed. Academic Press.

Langs, G., Tie, Y., Rigolo, L., Golby, A. and Golland, P. (2010). Functional Geometry Alignment and Localization of Brain Areas. In: Advances in Neural Information Processing Systems 23. Ed. by J. D. Lafferty, C. K. I. Williams, J. Shawe-Taylor, R. S. Zemel and A. Culotta. Curran Associates, Inc., pp. 1225–1233.

Langs, G., Wang, D., Golland, P., Mueller, S., Pan, R. et al. (2016). Identifying Shared Brain Networks in Individuals by Decoupling Functional and Anatomical Variability. In: Cerebral Cortex 26.10 (Sept. 2016), pp. 4004–4014. eprint:/oup/backfile/Content_public/Journal/cercor/26/10/10.1093_cercor_bhv189/2/bhv189.pdf.

Laumann, T. O., Gordon, E. M., Adeyemo, B., Snyder, A. Z., Joo, S. J. et al. (2015). Functional System and Areal Organization of a Highly Sampled Individual Human Brain. In: Neuron 87.3 (Aug. 2015), pp. 657–670.

Laumann, T. O., Snyder, A. Z., Mitra, A., Gordon, E. M., Gratton, C. et al. (2017). On the Stability of BOLD fMRI Correlations. In: Cerebral Cortex 27.10 (Oct. 2017), pp. 4719–4732.

Lee, J.-H., Hashimoto, R., Wible, C. G. and Yoo, S.-S. (2011). Investigation of spectrally coherent resting-state networks using non-negative matrix factorization for functional MRI data. In: International Journal of Imaging Systems and Technology 21.2, pp. 211–222.

Li, H., Satterthwaite, T. D. and Fan, Y. (2017). Large-scale sparse functional networks from resting state fMRI. In: NeuroImage 156 (Aug. 2017), pp. 1–13.

Liu, W., Awate, S. P. and Fletcher, P. T. (2012). Group Analysis of Resting-State fMRI by Hierarchical Markov Random Fields. In: Medical Image Computing and Computer-Assisted Intervention—MICCAI 2012. Ed. by N. Ayache, H. Delingette, P. Golland and K. Mori. Vol. 7512. Lecture Notes in Computer Science. Springer Berlin Heidelberg, pp. 189–196.

Llera, A., Wolfers, T., Mulders, P. and Beckmann, C. F. (2019). Inter-individual differences in human brain structure and morphology link to variation in demographics and behavior. In: eLife 8 (July 2019). Ed. by M. Helmstaedter, R. B. Ivry, F. Pestilli and J. P. Lerch, e44443.

MacKay, D. J. C. (2003). Information Theory, Inference and Learning Algorithms. Cambridge University Press.

Marcus, D. S., Harms, M. P., Snyder, A. Z., Jenkinson, M., Wilson, J. A. et al. (2013). Human Connectome Project informatics: Quality control, database services, and data visualization. In: NeuroImage 80 (Oct. 2013), pp. 202–219.

Marrelec, G., Krainik, A., Duffau, H., Pélégrini-Issac, M., Lehéricy, S. et al. (2006). Partial correlation for functional brain interactivity investigation in functional MRI. In: NeuroImage 32.1 (Aug. 2006), pp. 228–237.

Mazumder, R., Hastie, T. and Tibshirani, R. (2010). Spectral Regularization Algorithms for Learning Large Incomplete Matrices. In: Journal of Machine Learning Research 11, pp. 2287–2322.

McDaniel, M. A. (2005). Big-brained people are smarter: A meta-analysis of the relationship between in vivo brain volume and intelligence. In: Intelligence 33.4, pp. 337–346.

Mennes, M., Biswal, B. B., Castellanos, F. X. and Milham, M. P. (2013). Making data sharing work: The FCP/INDI experience. In: NeuroImage 82 (Nov. 2013), pp. 683–691.

Mensch, A., Mairal, J., Thirion, B. and Varoquaux, G. (2017). Stochastic Subsampling for Factorizing Huge Matrices. In: ArXiv e-prints (Jan. 2017). 1701.05363 [stat.ML].

Miller, K. L., Alfaro-Almagro, F., Bangerter, N. K., Thomas, D. L., Yacoub, E. et al. (2016). Multimodal population brain imaging in the UK Biobank prospective epidemiological study. In: Nature Neuroscience 19 (Nov. 2016), pp. 1523–1536.

Mitchell, T. J. and Beauchamp, J. J. (1988). Bayesian Variable Selection in Linear Regression. In: Journal of the American Statistical Association 83.404, pp. 1023–1032. eprint: http://www.tandfonline.com/doi/pdf/10.1080/01621459.198810478694.

Mueller, S., Wang, D., Fox, M. D., Yeo, B. T. T., Sepulcre, J. et al. (2013). Individual Variability in Functional Connectivity Architecture of the Human Brain. In: Neuron 77.3 (Feb. 2013), pp. 586–595.

Nickerson, L. D., Smith, S. M., Öngür, D. and Beckmann, C. F. (2017). Using Dual Regression to Investigate Network Shape and Amplitude in Functional Connectivity Analyses. In: Frontiers in Neuroscience 11.115 (Mar. 2017).

Noble, K. G., Houston, S. M., Brito, N. H., Bartsch, H., Kan, E. et al. (2015). Family income, parental education and brain structure in children and adolescents. In: Nature Neuroscience 18 (Mar. 2015), pp. 773–778.

Noble, S., Scheinost, D., Finn, E. S., Shen, X., Papademetris, X. et al. (2017). Multisite reliability of MR-based functional connectivity. In: NeuroImage 146 (Feb. 2017), pp. 959–970.

Pervaiz, U., Vidaurre, D., Woolrich, M. W. and Smith, S. M. (2020). Optimising network modelling methods for fMRI. In: NeuroImage 211 (May 2020), p. 116604.

Poldrack, R. A., Barch, D., Mitchell, J., Wager, T., Wagner, A. et al. (2013). Toward open sharing of task-based fMRI data: the OpenfMRI project. In: Frontiers in Neuroinformatics 7.12 (July 2013).

Poldrack, R. A., Laumann, T. O., Koyejo, O., Gregory, B., Hover, A. et al. (2015). Long-term neural and physiological phenotyping of a single human. In: Nature Communications 6.8885 (Dec. 2015).

Power, J. D., Barnes, K. A., Snyder, A. Z., Schlaggar, B. L. and Petersen, S. E. (2012). Spurious but systematic correlations in functional connectivity MRI networks arise from subject motion. In: NeuroImage 59.3 (Feb. 2012), pp. 2142–2154.

Power, J. D., Plitt, M., Laumann, T. O. and Martin, A. (2017). Sources and implications of whole-brain fMRI signals in humans. In: NeuroImage 146 (Feb. 2017), pp. 609–625.

Preti, M. G. and Van De Ville, D. (2017). Dynamics of functional connectivity at high spatial resolution reveal long-range interactions and fine-scale organization. In: Scientific Reports 7.1 (Oct. 2017), p. 12773.

Qing, Z. and Gong, G. (2016). Size matters to function: Brain volume correlates with intrinsic brain activity across healthy individuals. In: NeuroImage 139 (Oct. 2016), pp. 271–278.

Raemaekers, M., Schellekens, W., Wezel, R. J. A. van, Petridou, N., Kristo, G. et al. (2014). Patterns of resting state connectivity in human primary visual cortical areas: A 7 T fMRI study. In: NeuroImage 84 (Jan. 2014), pp. 911–921.

Raichle, M. E., MacLeod, A. M., Snyder, A. Z., Powers, W. J., Gusnard, D. A. et al. (2001). A default mode of brain function. In: Proceedings of the National Academy of Sciences 98.2 (Jan. 2001), pp. 676–682. eprint: http://www.pnas.org/content/98/2/676.full.pdf+html.

Reiss, A. L., Abrams, M. T., Singer, H. S., Ross, J. L. and Denckla, M. B. (1996). Brain development, gender and IQ in children: A volumetric imaging study. In: Brain 119.5, pp. 1763–1774. eprint: http://brain.oxfordjournals.org/content/119/5/1763.full.pdf+html.

Robert, P. and Escoufier, Y. (1976). A Unifying Tool for Linear Multivariate Statistical Methods: The RV-Coefficient. English. In: Journal of the Royal Statistical Society. Series C (Applied Statistics) 25.3, pp. 257–265.

Robinson, E. C., Garcia, K., Glasser, M. F., Chen, Z., Coalson, T. S. et al. (2018). Multimodal surface matching with higher-order smoothness constraints. In: NeuroImage 167 (Feb. 2018), pp. 453–465.

Robinson, E. C., Jbabdi, S., Glasser, M. F., Andersson, J., Burgess, G. C. et al. (2014). MSM: A new flexible framework for Multimodal Surface Matching. In: NeuroImage 100, pp. 414–426.

Sala-Llonch, R., Smith, S. M., Woolrich, M. W. and Duff, E. P. (2019). Spatial parcellations, spectral filtering, and connectivity measures in fMRI: Optimizing for discrimination. In: Human Brain Mapping 40.2 (Feb. 2019), pp. 407–419. eprint: https://onlinelibrary.wiley.com/doi/pdf/10.1002/hbm.24381.

Salehi, M., Greene, A. S., Karbasi, A., Shen, X., Scheinost, D. et al. (2020). There is no single functional atlas even for a single individual: Functional parcel definitions change with task. In: NeuroImage 208 (Mar. 2020), p. 116366.

Salehi, M., Karbasi, A., Scheinost, D. and Constable, R. T. (2017). A Submodular Approach to Create Individualized Parcellations of the Human Brain. In: Medical Image Computing and Computer Assisted Intervention—MICCAI 2017. Ed. by M. Descoteaux, L. Maier-Hein, A. Franz, P. Jannin, D. L. Collins et al. Cham: Springer International Publishing, pp. 478–485.

Satterthwaite, T. D., Wolf, D. H., Loughead, J., Ruparel, K., Elliott, M. A. et al. (2012). Impact of in-scanner head motion on multiple measures of functional connectivity: Relevance for studies of neurodevelopment in youth. In: NeuroImage 60.1 (Mar. 2012), pp. 623–632.

Scott, A., Courtney, W., Wood, D., De la Garza, R., Lane, S. et al. (2011). COINS: An Innovative Informatics and Neuroimaging Tool Suite Built for Large Heterogeneous Datasets. In: Frontiers in Neuroinformatics 5.33 (Dec. 2011).

Shackman, A. J., Salomons, T. V., Slagter, H. A., Fox, A. S., Winter, J. J. et al. (2011). The integration of negative affect, pain and cognitive control in the cingulate cortex. In: Nature Reviews Neuroscience 12.3 (Mar. 2011), pp. 154–167.

Shaw, P., Greenstein, D., Lerch, J., Clasen, L., Lenroot, R. et al. (2006). Intellectual ability and cortical development in children and adolescents. In: Nature 440 (Mar. 2006), pp. 676–679.

Shehzad, Z., Kelly, A. M. C., Reiss, P. T., Gee, D. G., Gotimer, K. et al. (2009). The Resting Brain: Unconstrained yet Reliable. In: Cerebral Cortex 19.10 (Oct. 2009), pp. 2209–2229. eprint: http://cercor.oxfordjournals.org/content/19/10/2209.full.pdf+html.

Shirer, W. R., Ryali, S., Rykhlevskaia, E., Menon, V. and Greicius, M. D. (2011). Decoding Subject-Driven Cognitive States with Whole-Brain Connectivity Patterns. In: Cerebral Cortex 22.1 (May 2011), pp. 158–165. eprint: https://academic.oup.com/cercor/article-pdf/22/1/158/14096754/bhr099.pdf.

Shmuel, A., Yacoub, E., Chaimow, D., Logothetis, N. K. and Ugurbil, K. (2007). Spatio-temporal point-spread function of fMRI signal in human gray matter at 7 Tesla. In: NeuroImage 35.2 (Apr. 2007), pp. 539–552.

Shulman, G. L., Fiez, J. A., Corbetta, M., Buckner, R. L., Miezin, F. M. et al. (1997). Common Blood Flow Changes across Visual Tasks: II. Decreases in Cerebral Cortex. In: Journal of Cognitive Neuroscience 9.5 (Oct. 1997), pp. 648–663.

Smith, S. M., Feinberg, D. A., Griffanti, L., Harms, M. P., Kelly, M. et al. (2013a). Resting-state fMRI in the Human Connectome Project. In: NeuroImage 80 (Oct. 2013), pp. 144–168.

Smith, S. M., Hyvärinen, A., Varoquaux, G., Miller, K. L. and Beckmann, C. F. (2014). Group-PCA for very large fMRI datasets. In: NeuroImage 101 (Nov. 2014), pp. 738–749.

Smith, S. M., Miller, K. L., Moeller, S., Xu, J., Auerbach, E. J. et al. (2012). Temporally-independent functional modes of spontaneous brain activity. In: Proceedings of the National Academy of Sciences 109.8, pp. 3131–3136.

Smith, S. M., Miller, K. L., Salimi-Khorshidi, G., Webster, M., Beckmann, C. F. et al. (2011). Network modelling methods for FMRI. In: NeuroImage 54.2, pp. 875–891.

Smith, S. M., Nichols, T. E., Vidaurre, D., Winkler, A. M., Behrens, T. E. J. et al. (2015). A positive-negative mode of population covariation links brain connectivity, demographics and behavior. In: Nature Neuroscience 18.11 (Nov. 2015), pp. 1565–1567.

Smith, S. M., Vidaurre, D., Beckmann, C. F., Glasser, M. F., Jenkinson, M. et al. (2013b). Functional connectomics from resting-state fMRI. In: Trends in Cognitive Sciences 17.12 (Dec. 2013), pp. 666–682.

Stegle, O., Lippert, C., Mooij, J., Lawrence, N. and Borgwardt, K. (2011). Efficient inference in matrix-variate Gaussian models with iid observation noise. In: Advances in Neural Information Processing Systems 24. Ed. by J. Shawe-Taylor, R. Zemel, P. Bartlett, F. Pereira and K. Weinberger, pp. 630–638.

Stein, J. L., Medland, S. E., Vasquez, A. A., Hibar, D. P., Senstad, R. E. et al. (2012). Identification of common variants associated with human hippocampal and intracranial volumes. In: Nature Genetics 44.5 (May 2012), pp. 552–561.

Stephan, K. E., Iglesias, S., Heinzle, J. and Diaconescu, A. O. (2015). Translational Perspectives for Computational Neuroimaging. In: Neuron 87.4 (Aug. 2015), pp. 716–732.

Stephan, K. E., Schlagenhauf, F., Huys, Q. J. M., Raman, S., Aponte, E. A. et al. (2017). Computational neuroimaging strategies for single patient predictions. In: NeuroImage 145, Part B (Jan. 2017), pp. 180–199.

Tagliazucchi, E. and Laufs, H. (2014). Decoding Wakefulness Levels from Typical fMRI Resting-State Data Reveals Reliable Drifts between Wakefulness and Sleep. In: Neuron 82.3, pp. 695–708.

Thompson, P. M., Stein, J. L., Medland, S. E., Hibar, D. P., Vasquez, A. A. et al. (2014). The ENIGMA Consortium: large-scale collaborative analyses of neuroimaging and genetic data. In: Brain Imaging and Behavior 8.2 (June 2014), pp. 153–182.

Titsias, M. and Lázaro-Gredilla, M. (2011). Spike and Slab Variational Inference for Multi-Task and Multiple Kernel Learning. In: Advances in Neural Information Processing Systems 24. Ed. by J. Shawe-Taylor, R. Zemel, P. Bartlett, F. Pereira and K. Weinberger, pp. 2339–2347.

Tong, T., Aganj, I., Ge, T., Polimeni, J. R. and Fischl, B. (2017). Functional density and edge maps: Characterizing functional architecture in individuals and improving cross-subject registration. In: NeuroImage 158 (Sept. 2017), pp. 346–355.

Van Dijk, K. R. A., Sabuncu, M. R. and Buckner, R. L. (2012). The influence of head motion on intrinsic functional connectivity MRI. In: NeuroImage 59.1 (Jan. 2012), pp. 431–438.

Van Essen, D. C. and Dierker, D. L. (2007). Surface-Based and Probabilistic Atlases of Primate Cerebral Cortex. In: Neuron 56.2 (Oct. 2007), pp. 209–225.

Van Essen, D. C., Glasser, M. F., Dierker, D. L., Harwell, J. and Coalson, T. (2012a). Parcellations and Hemispheric Asymmetries of Human Cerebral Cortex Analyzed on Surface-Based Atlases. In: Cerebral Cortex 22.10 (Oct. 2012), pp. 2241–2262. eprint: http://cercor.oxfordjournals.org/content/22/10/2241.full.pdf+html.

Van Essen, D. C., Smith, S. M., Barch, D. M., Behrens, T. E. J., Yacoub, E. et al. (2013). The WU-Minn Human Connectome Project: An overview. In: NeuroImage 80 (Oct. 2013), pp. 62–79.

Van Essen, D. C., Ugurbil, K., Auerbach, E., Barch, D., Behrens, T. E. J. et al. (2012b). The Human Connectome Project: A data acquisition perspective. In: NeuroImage 62.4, pp. 2222–2231.

Vanderwal, T., Eilbott, J., Finn, E. S., Craddock, R. C., Turnbull, A. et al. (2017). Individual differences in functional connectivity during naturalistic viewing conditions. In: NeuroImage 157 (Aug. 2017), pp. 521–530.

Varoquaux, G., Gramfort, A., Pedregosa, F., Michel, V. and Thirion, B. (2011). Multi-subject Dictionary Learning to Segment an Atlas of Brain Spontaneous Activity. In: Information Processing in Medical Imaging. Ed. by G. Székely and H. K. Hahn. Vol. 6801. Lecture Notes in Computer Science. Springer Berlin Heidelberg, pp. 562–573.

Varoquaux, G., Raamana, P. R., Engemann, D. A., Hoyos-Idrobo, A., Schwartz, Y. et al. (2017). Assessing and tuning brain decoders: Cross-validation, caveats, and guidelines. In: NeuroImage 145, Part B (Jan. 2017), pp. 166–179.

Varoquaux, G., Sadaghiani, S., Pinel, P., Kleinschmidt, A., Poline, J. et al. (2010). A group model for stable multi-subject ICA on fMRI datasets. In: NeuroImage 51.1, pp. 288–299.

Vidaurre, D., Smith, S. M. and Woolrich, M. W. (2017). Brain network dynamics are hierarchically organized in time. In: Proceedings of the National Academy of Sciences 114.48 (Nov. 2017), pp. 12827–12832.

Wang, D., Buckner, R. L., Fox, M. D., Holt, D. J., Holmes, A. J. et al. (2015). Parcellating cortical functional networks in individuals. In: Nature Neuroscience 18.12 (Dec. 2015), pp. 1853–1860.

Winkler, A. M., Ridgway, G. R., Douaud, G., Nichols, T. E. and Smith, S. M. (2016). Faster permutation inference in brain imaging. In: NeuroImage 141 (Nov. 2016), pp. 502–516.

Winkler, A. M., Ridgway, G. R., Webster, M. A., Smith, S. M. and Nichols, T. E. (2014). Permutation inference for the general linear model. In: NeuroImage 92 (May 2014), pp. 381–397.

Winn, J. and Bishop, C. M. (2005). Variational message passing. In: Journal of Machine Learning Research 6, pp. 661–694.

Woolrich, M. W., Behrens, T. E. J. and Smith, S. M. (2004). Constrained linear basis sets for HRF modelling using Variational Bayes. In: NeuroImage 21.4, pp. 1748–1761.

Worsley, K. J., Evans, A. C., Marrett, S. and Neelin, P. (1996). A Three-Dimensional Statistical Analysis for CBF Activation Studies in Human Brain. In: Journal of Cerebral Blood Flow & Metabolism 12.6 (Nov. 1996), pp. 900–918.

Zang, Y.-F., He, Y., Zhu, C.-Z., Cao, Q.-J., Sui, M.-Q. et al. (2007). Altered baseline brain activity in children with ADHD revealed by resting-state functional MRI. In: Brain and Development 29.2 (Mar. 2007), pp. 83–91.

Zöller, D. M., Bolton, T. A. W., Karahanoglu, F. I., Eliez, S., Schaer, M. et al. (2019). Robust recovery of temporal overlap between network activity using transient-informed spatio-temporal regression. In: IEEE Transactions on Medical Imaging 38.1 (Jan. 2019), pp. 291–302.

Zou, Q.-H., Zhu, C.-Z., Yang, Y., Zuo, X.-N., Long, X.-Y. et al. (2008). An improved approach to detection of amplitude of low-frequency fluctuation (ALFF) for restingstate fMRI: Fractional ALFF. In: Journal of Neuroscience Methods 172.1 (July 2008), pp. 137–141.

Zuo, X.-N., Di Martino, A., Kelly, C., Shehzad, Z. E., Gee, D. G. et al. (2010a). The oscillating brain: Complex and reliable. In: NeuroImage 49.2 (Jan. 2010), pp. 1432–1445.

Zuo, X.-N., Kelly, C., Adelstein, J. S., Klein, D. F., Castellanos, F. X. et al. (2010b). Reliable intrinsic connectivity networks: Test-retest evaluation using ICA and dual regression approach. In: NeuroImage 49.3 (Feb. 2010), pp. 2163–2177.

