## Supplementary Material for "Modelling Subject Variability in the Spatial and Temporal Characteristics of Functional Modes"

26th June 2020

#### 1 Model

##### 1.1 Implementation

The code that does the probabilistic inference is written in **C++** (isocpp.org). The bulk of the computation is done using single precision floats as this reduces the memory footprint and results in a significant computational speed-up. This is handled by the **Armadillo** linear algebra library (Sanderson and Curtin [2016], arma.sourceforge.net) and **OpenBLAS** (github.com/xianyi/OpenBLAS). The inference procedure parallelises very neatly over subjects: holding the group parameters fixed, all the subject specific updates can be performed independently of each other. The code is parallelised using **OpenMP** (openmp.org) and also relies on several functions from the **Boost** libraries (boost.org).

Postprocessing and visualisation of the results is done in **Python** (python.org). This relies on **NumPy** (numpy.org), **SciPy** (scipy.org) and **NiBabel** (nipy.org/nibabel), and all plots are generated by **matplotlib** (matplotlib.org).

##### 1.2 Model Parameters

This manuscript, including the model description and analyses performed, is based on version 0.11.1 of PROFUMO. The full graphical model is shown in Supplementary Figure S1. The specific values that the various hyperpriors take can be found in the following file, git.fmrib.ox.ac.uk/samh/profumo/-/blob/0.11.1/C++/Source/ModelManager.c++, where the exact definition of the key distributions is available in git.fmrib.ox.ac.uk/samh/profumo/-/blob/0.11.1/Documentation/UpdateRules/

[a]FMRIB, Wellcome Centre for Integrative Neuroimaging, University of Oxford, Oxford, UK
[b]OHBA, Wellcome Centre for Integrative Neuroimaging, University of Oxford, Oxford, UK
[c]Translational Neuromodeling Unit, University of Zurich & ETH Zurich, Zurich, Switzerland
[d]Department of Radiology, Washington University Medical School, Saint Louis, USA
[e]Department of Paediatrics, University of Oxford, Oxford, UK

KeyDistributions/KeyDistributions.pdf. The key update rules can be found at git.fmrib.ox.ac.uk/samh/profumo/-/blob/0.11.1/Documentation/UpdateRules.

While there are a relatively large number of hyperparameters, there are several ways to simplify their specification. Firstly, the data normalisation procedure (see Appendices) is based on the spatial maps, amplitudes and time courses being approximately unit variance. Secondly, many the parameters of many conjugate priors have an interpretation as a number of 'pseudo-observations', and we set many of the group-level priors to be equivalent in strength to observing a small number of 'pseudo-subjects'. Taken together, these drastically reduce the number of free parameters we need to specify.

### 2 Results

#### 2.1 Simulations

Supplementary Figures S2 to S10 demonstrate the performance of the different approaches on simulated data, both with and without subject-specific structured artefacts. As per the manuscript itself, as well as PROFUMO and ICA-DR, we test dual regression starting with the ground-truth spatial maps (GTg-DR) and thresholded dual regression (ICA-DRt, GTg-DRt). For each metric, optimal performance is shown by the horizontal green line.

### References

Sanderson, C. and Curtin, R. (2016). *Armadillo: a template-based C++ library for linear algebra*. In: *The Journal of Open Source Software* 1.2 (June 2016).

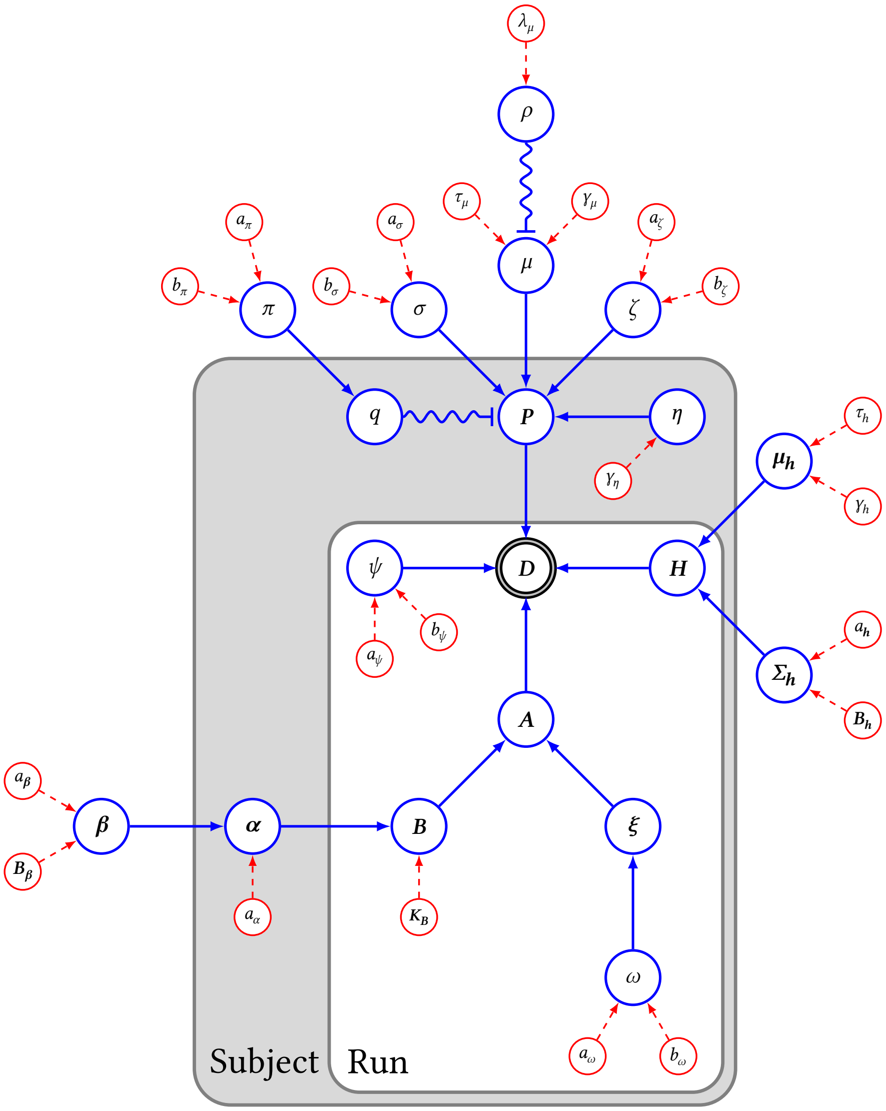


Supplementary Figure S1: Graphical representation of the full model structure. Variables are shown in the blue circles and the dependencies are depicted as the arrows joining them, with switches conditioned on indicator variables shown as wiggly lines. Pre-specified prior parameters are shown in red. The plates indicate which parameters are inferred at the subject and run levels, though we have omitted other dimensions and numbers of variables for simplicity.

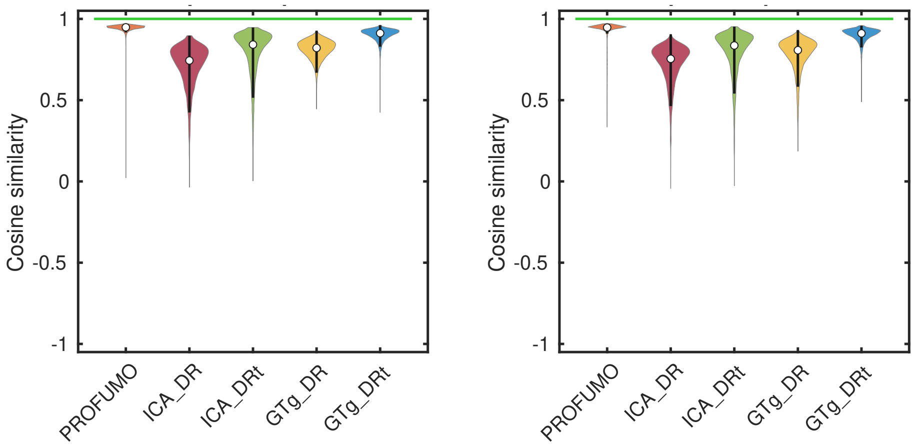


**(a)** Without artefacts. **(b)** With artefacts.

**Supplementary Figure S2:** Similarity of the ground truth and inferred subject-specific spatial maps. Inferred modes are paired to their ground-truth counterparts, and the cosine similarity is computed between each pair of full spatial maps. The distribution is over modes, subjects, and repeats.

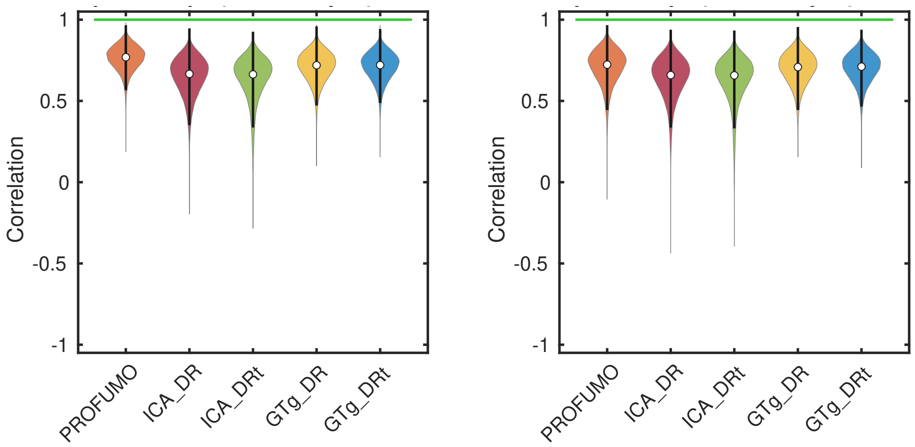


**(a)** Without artefacts. **(b)** With artefacts.

**Supplementary Figure S3:** Cross-subject similarity of the ground truth and inferred subject-specific spatial maps. Inferred modes are paired to their ground-truth counterparts, and for each voxel the correlation is computed between the true and inferred changes in spatial weights over subjects. The distribution is over non-zero voxels, modes, and repeats.

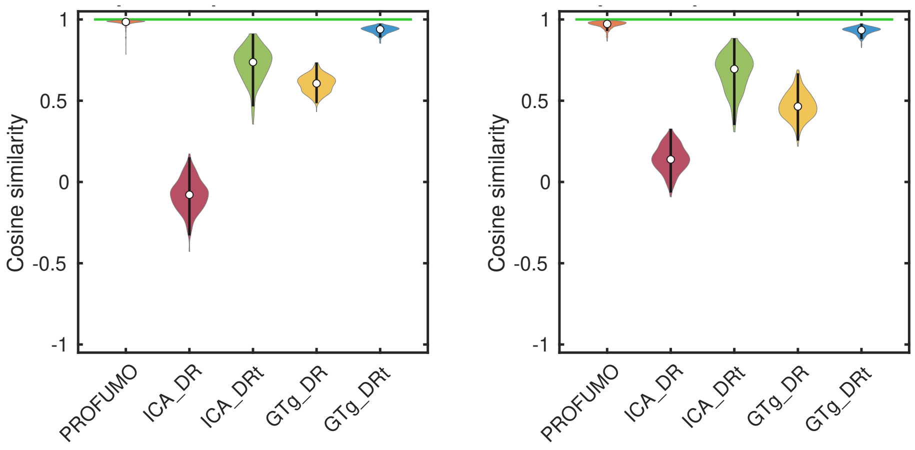


(a) Without artefacts. (b) With artefacts.

Supplementary Figure S4: Similarity of the ground truth and inferred subject-specific spatial map interactions. After inferred modes are paired to their ground-truth counterparts, the cosine similarity is computed between all the spatial maps for a given subject. The cosine similarity between the unwrapped upper-triangle of this similarity matrix and the ground truth is reported here. The distribution is over subjects and repeats.

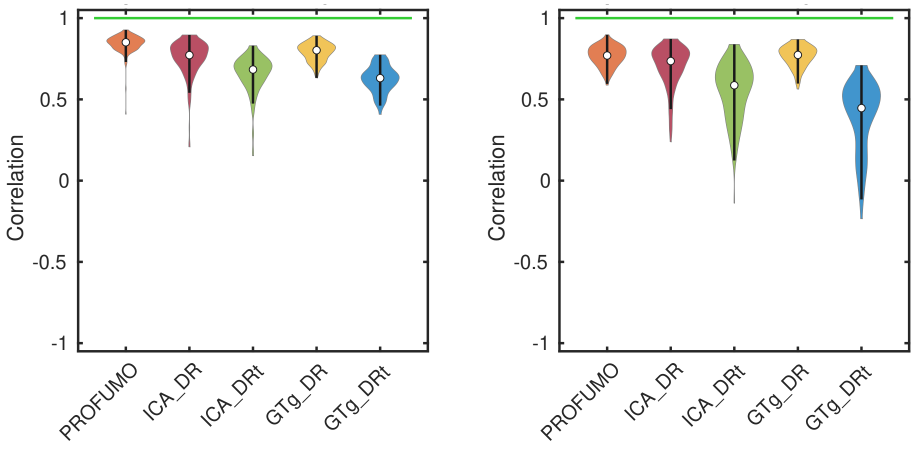


(a) Without artefacts. (b) With artefacts.

Supplementary Figure S5: Cross-subject similarity of the ground truth and inferred run-specific amplitudes. Inferred modes are paired to their ground-truth counterparts, and for each mode the correlation is computed between the true and inferred changes in amplitude over subjects. The distribution is over modes and repeats.

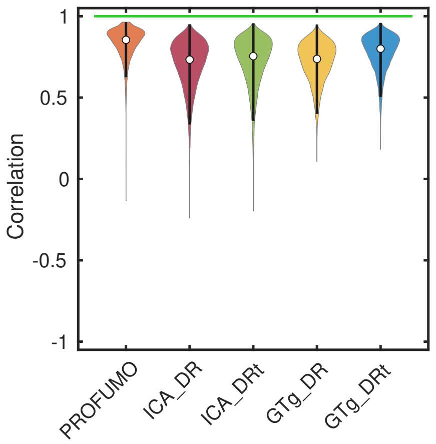


(a) Without artefacts.

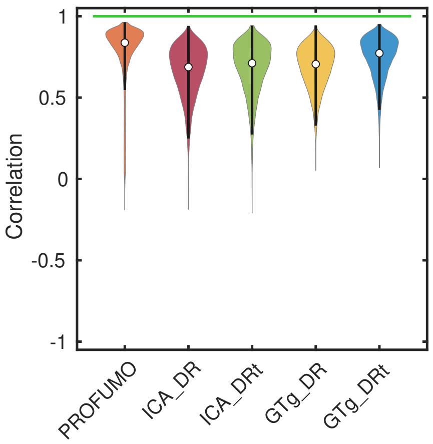


(b) With artefacts.

**Supplementary Figure S6:** Similarity of the ground truth and inferred run-specific timecourses. Inferred modes are paired to their ground-truth counterparts, and the correlation is computed between each pair of timecourses. The distribution is over modes, runs, and repeats.

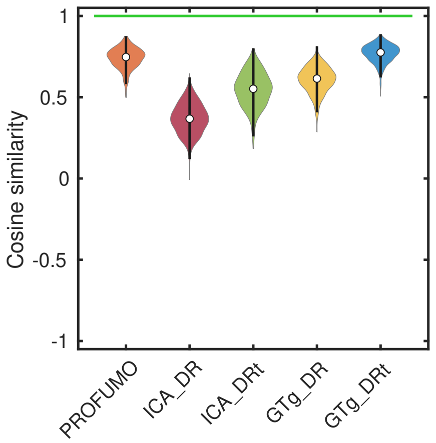


(a) Without artefacts.

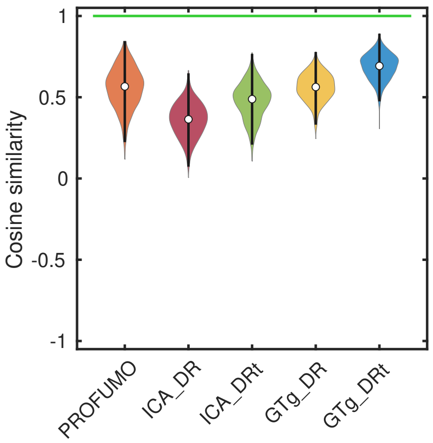


(b) With artefacts.

**Supplementary Figure S7:** Similarity of the ground truth and inferred run-specific netmats. After inferred modes are paired to their ground-truth counterparts, the regularised partial correlation is computed between all the timecourses for a given run. The cosine similarity between the unwrapped upper-triangle of this network matrix and the ground truth is reported here. The distribution is over runs and repeats.

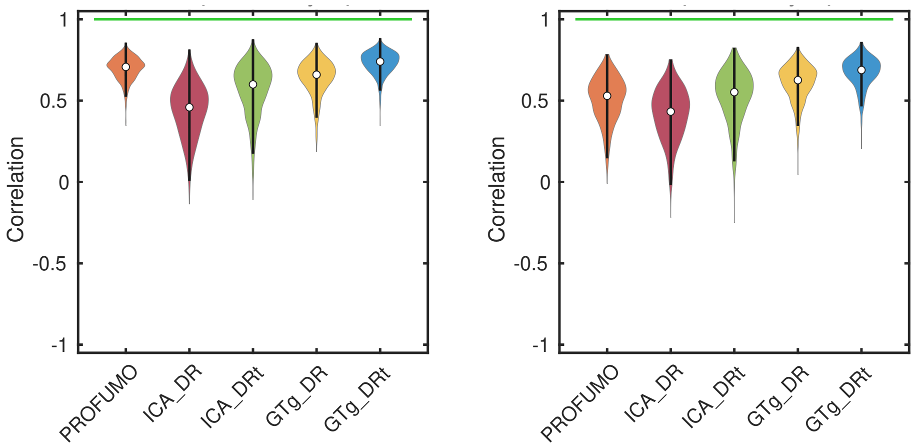


(a) Without artefacts. (b) With artefacts.

Supplementary Figure S8: Cross-subject similarity of the ground truth and inferred run-specific netmats. After inferred modes are paired to their ground-truth counterparts, the regularised partial correlation is computed between all the timecourses for a given run. For each network edge (i.e. element of the netmat) the correlation is computed between the true and inferred changes in edge strength over subjects (i.e. changes in functional coupling). The distribution is over edges and repeats.

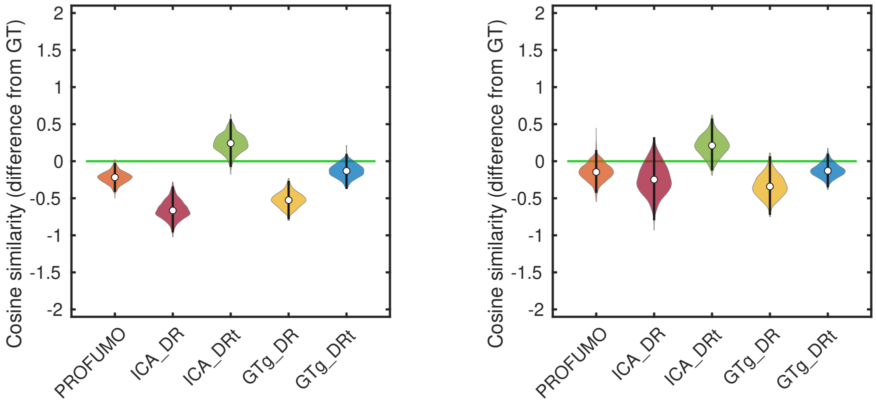


(a) Without artefacts. (b) With artefacts.

Supplementary Figure S9: Biases in the inferred patterns of spatial and temporal interactions between modes. Inferred modes are paired to their ground-truth counterparts, and the run-specific spatial interactions and temporal netmats are calculated as before. We calculate the cosine similarity between the unwrapped upper-triangle of these matrices and report the difference between this and the ground truth value (i.e. the bias). A negative value implies that a strong spatial correlation between any two modes tends to reduce the strength of the inferred functional coupling. Alternatively, a positive value implies that the overall pattern of functional coupling between modes looks more like the spatial pattern than it should. The distribution is over runs and repeats.

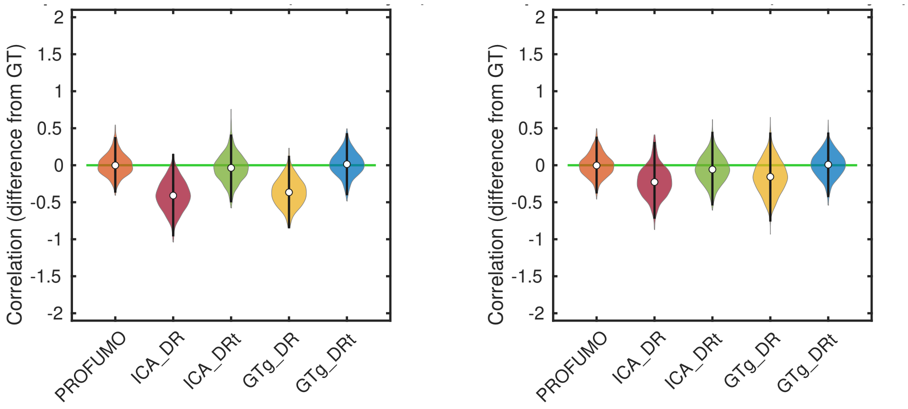


**(a)** Without artefacts. **(b)** With artefacts.

**Supplementary Figure S10:** Biases in the inferred patterns of spatial and temporal interactions across subjects. Inferred modes are paired to their ground-truth counterparts, and the run-specific spatial interactions and temporal netmats are calculated as before. For each edge, we calculate the correlation between changes in the strength of the spatial and temporal similarities across runs, and we report the difference between this and the ground truth value (i.e. the bias). A negative value implies that, for a given edge, an above average spatial correlation in a specific subject tends to result in the strength of that subject's inferred functional coupling being below average. The distribution is over edges and repeats.
